# Comparison of Rosetta flexible-backbone computational protein design methods on binding interactions

**DOI:** 10.1101/674291

**Authors:** Amanda L. Loshbaugh, Tanja Kortemme

## Abstract

Computational design of binding sites in proteins remains difficult, in part due to limitations in our current ability to sample backbone conformations that enable precise and accurate geometric positioning of side chains during sequence design. Here we present a benchmark framework for comparison between flexible-backbone design methods applied to binding interactions. We quantify the ability of different flexible backbone design methods in the widely used protein design software Rosetta to recapitulate observed protein sequence profiles assumed to represent functional protein/protein and protein/small molecule binding interactions. The CoupledMoves method, which combines backbone flexibility and sequence exploration into a single acceptance step during the sampling trajectory, better recapitulates observed sequence profiles than the BackrubEnsemble and FastDesign methods, which separate backbone flexibility and sequence design into separate acceptance steps during the sampling trajectory. Flexible-backbone design with the CoupledMoves method is a powerful strategy for reducing sequence space to generate targeted libraries for experimental screening and selection.

## INTRODUCTION

Computational protein design searches for sequences that adopt desired structures and functions. Most generally, computational design methods require (i) algorithms to efficiently search the vast sequence and conformational space accessible to proteins, and (ii) effective energy functions to rank the solutions. Both of these requirements necessitate approximations. Design energy functions are often simplified while considering atomic detail (Alford et al., 2017; Gordon et al., 1999), and the search space of sequences and conformations is typically limited by reducing degrees of freedom in a design simulation. One early approximation was to leave the backbone fixed while sampling rotameric side chain conformations during sequence design (Dahiyat & Mayo, 1997; Ponder & Richards, 1987). While the fixed backbone approximation is useful for computational efficiency, it is rarely sufficiently accurate as flexibility is a hallmark of naturally occurring functional proteins and backbones shift to accommodate side chain mutations arising during evolution or design (Baldwin et al., 1993; Davis et al., 2006; Desjarlais & Handel, 1999; Keedy et al., 2012). Highly stable, idealized folds can be designed *de novo* (Koga et al., 2012; Rocklin et al., 2017), but design of proteins with new functions remains challenging. In most cases where new functions have been designed computationally, the designed protein is modeled on natural “scaffold” proteins with minimal changes in backbone conformation (Correia et al., 2014; Jiang et al., 2008; Procko et al., 2014; Rothlisberger et al., 2008; Tinberg et al., 2013), although there are recent notable exceptions of functions designed into *de novo* proteins (Dou et al., 2018; Silva et al., 2019). Moreover, attaining sufficiently high activities typically requires optimization of the desired function by directed evolution (Fleishman et al., 2011; Rothlisberger et al., 2008; Tinberg et al., 2013). Function often depends on hydrogen bonds, which require precise backbone and side chain geometry, which remains difficult to design (Dou et al., 2017) especially when a novel function requires “reshaping” of an existing protein conformation (Kundert & Kortemme, 2019).

Various strategies have been proposed to model backbone flexibility in design, such as small random perturbations in torsional or Cartesian space (Davey & Chica, 2014, 2017; Desjarlais & Handel, 1999; Larson et al., 2002), normal mode analysis (Fu et al., 2007), or “backrub” motions shown to underlie commonly observed protein structural heterogeneity in high resolution crystal structures (Davis et al., 2006) that have proven useful to model structural changes in response to mutations (Georgiev et al., 2008; Keedy et al., 2012; Smith & Kortemme, 2008). Several strategies have used protein backbone ensembles for design, which are typically generated computationally employing various sampling methods (Davey & Chica, 2014, 2017; Ding & Dokholyan, 2006) including backrub motions (Friedland et al., 2009; Friedland et al., 2008; Humphris & Kortemme, 2008; Humphris-Narayanan et al., 2012) but can also be taken from crystal structures (Friedland et al., 2009; Humphris-Narayanan et al., 2012; Larson et al., 2002).

Within the structure modeling and design program Rosetta (Leaver-Fay et al., 2011), backbone flexibility has been treated in a number of ways. These include (i) generation of new protein backbones by assembly from peptide fragments which demonstrated success in *ab initio* structure prediction (Simons et al., 1999; Simons et al., 1997), (ii) iterating between sequence design via Monte Carlo search and structure optimization via minimization (Kuhlman et al., 2003; Saunders & Baker, 2005), (iii) a robotics-inspired kinematic closure (KIC) algorithm (Coutsias et al., 2004) shown to model loop conformations with sub-Angstrom accuracy (Mandell et al., 2009), and (iv) the Backrub algorithm benchmarked on recapitulation of known sequences (Friedland et al., 2009; Humphris & Kortemme, 2008; Ollikainen & Kortemme, 2013; Smith & Kortemme, 2010, 2011). Most flexible backbone design methods iterate between sequence design on a fixed backbone and structural optimization on a fixed sequence (Kuhlman et al., 2003), which effectively uncouples sequence changes from direct influence on backbone structure. In contrast, the “CoupledMoves” method in Rosetta (Ollikainen et al., 2015), combines side chain and backbone moves using Rosetta backrub sampling (Smith & Kortemme, 2008) in a single design step.

While Rosetta flexible backbone design has been successfully applied to forward engineering of new functions (Chevalier et al., 2017; Dou et al., 2018; Kapp et al., 2012), different methods have not been directly compared for accuracy using common benchmark datasets. Here, we describe such a benchmark comparison of three different flexible-backbone design methods in Rosetta: CoupledMoves (Ollikainen et al., 2015), BackrubEnsemble (Ollikainen & Kortemme, 2013; Smith & Kortemme, 2010, 2011), and FastDesign, which combines sequence design with the Rosetta FastRelax method (Khatib et al., 2011; Tyka et al., 2011) to move the backbone. We focus on methods within the openly available Rosetta framework because they use the same energy function, which allows us to directly compare different methods of sampling backbone flexibility. We evaluate each of the methods on its ability to recapitulate “tolerated sequence space” for binding interactions. We define tolerated sequence space as experimentally selected or naturally occurring sequences consistent with a functional binding interaction with a small molecule or protein binding partner.

We find that CoupledMoves recapitulates tolerated sequence space and individual stabilizing mutations more accurately than FastDesign or BackrubEnsemble. We introduce an updated version of the CoupledMoves algorithm (CM-KIC) that uses kinematic closure (KIC) in place of the original backrub backbone mover, which leads to further small improvements in performance. The coupled algorithm allows subtle conformational shifts in backbone torsions which accommodate favorable side chain rotamers, in turn leading to more accurate prediction of side chain interactions. We also analyze shortcomings of the design methods that highlight areas for improvement.

## MATERIALS AND METHODS

### Design methods

Design protocols used Rosetta revision number 60351 and energy function ref2015 (Alford et al., 2017; Park et al., 2016). For each method, we used standard parameters and settings previously reported in benchmarks or design applications, except for the new coupled moves (CM) methods (CM-FKIC and CM-WKIC, see below) reported here. Command lines for each method can be found in the supplement.

#### FastDesign

FastDesign is based on the FastRelax protocol in Rosetta described in (Khatib et al., 2011; Tyka et al., 2011). Briefly, FastRelax consists of inner cycles of rotamer repacking and backbone and side chain torsion minimization with progressively higher weight on the repulsive part of the van der Waals energy function component (from 2% to 100% of its total value). FastDesign uses an analogous protocol but allows side chain design in addition to repacking. During FastDesign, we used harmonic coordinate constraints to keep backbone heavy atoms close to their starting position, and the weight of the constraints was ramped down from 1.0 to 0.0 during the course of each inner simulated annealing cycle. Constraint and repulsive weights were ramped five times, during five outer cycles. For each input protein structure, 400 designs were generated in independent design trajectories. Command line arguments are provided in the Supplement.

#### BackrubEnsemble

The BackrubEnsemble method is described in (Smith & Kortemme, 2010, 2011). Briefly, the method generates a structural ensemble with backbone conformational variation using the backrub algorithm implemented in Rosetta (Smith & Kortemme, 2008), and then carries out fixed-backbone side chain design on each member of the ensemble. 400 ensemble members were generated using 10,000 backrub trials, a temperature of 1.2, and a backbone segment length of 3-12 atoms. Command line arguments are shown in the Supplement.

#### Forced BackrubEnsemble design

“Forced” BackrubEnsemble design forces sequence design to choose the known consensus side chain at certain positions. Forced design was applied to Glutathione Reductase positions E50 and D331, and DIG10 positions Y34, Y101, and Y115. For each protein, 100 forced simulations were run, using as input the first 100 members of the same BackrubEnsemble on which typical design was performed.

#### CoupledMoves

The CoupledMoves method was used as described in (Ollikainen et al., 2015). Briefly, each coupled move had a 90% probability of being a backbone and side-chain move, and a 10% probability of being a ligand move. Each simulation was run for 1,000 moves and 400 simulations were run for each protein-ligand or protein-protein complex. All unique amino acid sequences accepted during each simulation were output into a FASTA file, and the resulting 400 FASTA files were pooled, including redundancy, for analysis. Command line arguments are provided in the Supplement.

#### CoupledMoves with Kinematic Closure (CM-KIC)

Two different methods of modeling backbone flexibility are implemented in CoupledMoves. The first method uses the Backrub algorithm (Smith & Kortemme, 2008) and was originally described in (Ollikainen et al., 2015). The second method uses kinematic closure (KIC) (Coutsias et al., 2004; Mandell et al., 2009) and is implemented in CoupledMoves here. Kinematic closure in Rosetta (Mandell et al., 2009) generates conformations of backbone segments by sampling non-pivot torsions in the segment and then analytically determining values for 6 pivot torsions to close the loop. For the CM-FKIC (KIC using fragments) method, non-pivot torsions are sampled from peptide fragments (Simons et al., 1997) taken from the protein structure database (PDB). For the CM-WKIC (KIC using a “walking” perturbation) method, non-pivot torsions are adjusted by a random value from a Gaussian distribution centered around zero and with a standard deviation of 3°. In each case, the remaining six pivot torsions are then solved analytically to close the loop. Command line arguments are provided in the Supplement.

#### Ligand handling

Rosetta requires ligands to be described by a params file, which contains information defining the ligand’s atom types, bond geometry, and chemical connectivity. We generated params files from PDB structures using Rosetta’s `molfile_to_params.py` utility script. We did not model multiple ligand conformers except for DIG, for which the DIG ligand conformer library used during DIG10 design (Tinberg et al., 2013) was obtained from the authors.

CoupledMoves samples ligand rigid-body translation and rotation in all cases. FastDesign minimizes ligand torsional degrees of freedom in addition to backbone torsion angles during its minimization step. BackrubEnsemble and FixBB do not sample ligand movement.

### Benchmark datasets

#### Cofactor binding sites

This dataset is described in detail in (Ollikainen et al., 2015). Briefly, the dataset is comprised of seven protein families, each containing a conserved small molecule cofactor binding site (**Table S1**). The crystal structure with the highest resolution available was chosen as the starting point for design. As in (Ollikainen et al., 2015), positions with a side-chain heavy atom within 6 Å of any heavy atom in the cofactor ligand were allowed to design to any amino acid identity, and positions that could clash with designable positions were allowed to repack (change conformation but not identity) (**Table S2**). Known profiles were obtained from natural sequences of these binding sites as described in (Ollikainen et al., 2015).

#### Enzyme specificity

This dataset is described in detail in (Ollikainen et al., 2015). Briefly, the dataset is comprised of 10 enzymes for which there are experimentally validated specificity-altering mutations in the ligand binding sites (**Table S3**). As in (Ollikainen et al., 2015), design was carried out with either the native or the non-native substrate/substrate analog. Positions with heavy atoms within 4.5 Å of any ligand atoms differing between the native and non-native substrate were allowed to design to any amino acid identity, and positions that could clash with designable positions (as described in (Ollikainen et al., 2015)) were allowed to repack (**Table S4**). Structures were prepared as described in (Ollikainen et al., 2015). Briefly, for each enzyme four types of structures were prepared: 1) the native enzyme with the native ligand, 2) the mutant enzyme(s) with the non-native ligand, 3) the native enzyme with the non-native ligand, and 4) the mutant enzyme with the native ligand.

#### DIG10

The DIG10 dataset was taken from (Tinberg et al., 2013). Briefly, DIG10 is a computationally designed protein that has been engineered to bind the small molecule digoxigenin (DIG) (Tinberg et al., 2013). A computational design, DIG10, was subjected to selection by yeast surface display, first of a single-site saturation mutagenesis library, then of a combinatorial library of beneficial mutations identified in the first selection, yielding variant DIG10.1. DIG10.1 was then subjected to site saturation mutagenesis (SSM) and selections using yeast surface display to probe the effect of mutations, which our computational design protocol seeks to recapitulate. For input, we used the crystal structure of wild-type protein (PDB: 1Z1S) on which DIG10 was designed, the sequence of DIG10.1 (which we placed onto the 1Z1S scaffold using the Rosetta fixed backbone (FixBB) design protocol), and the digoxigenin conformation from the DIG10.2/digoxigenin complex (PDB: 4J8T), where DIG10.2 is a DIG10.1 variant containing additional mutations from the SSM selection). Digoxigenin was placed into the 1Z1S scaffold by using PyMOL to align 4J8T and 1Z1S, then combining the digoxigenin molecule from 4J8T and the protein structure from 1Z1S into a new PDB file. The known profile represents the frequency equivalent (*F_equiv_*, described below) of the selection experiment on the DIG10.1 SSM library. The 39 positions selected for experimental site saturation in (Tinberg et al., 2013) were allowed to design to only those amino acid identities with high enough sequencing counts to be included in the enrichment and depletion calculations in (Tinberg et al., 2013) (**Table S5, Table S6**). We note that the experimental screen mutated 1-2 position at a time, whereas we design multiple positions simultaneously. In CoupledMoves design, 30 positions were allowed to repack based on the possibility of clashes with designed positions; in design by non-coupled methods (FastDesign, BackrubEnsemble, FixBB), all positions were allowed to repack (**Table S5**).

#### Fen49

The Fen49 dataset was taken from (Bick et al., 2017). Fen49 is a computationally designed protein that has been engineered to bind the small molecule fentanyl (Fen). The original computational design, Fen49, has an affinity of 6.9 µM for a fentanyl-bovine serum albumin conjugate. After four rounds of selection starting from Fen49, a combination of two substitutions, A78V and A172I, was identified to produce a variant with a 100-fold improved affinity of 64 nM. We used the wild-type protein (PDB: 2QZ3), on which the sequence of Fen49 was modeled using Rosetta FixBB as input to our design simulations. The fentanyl conformation from designed fentanyl binder Fen49*/fentanyl complex (PDB: 5TZO, where Fen49* is a Fen49 Y88A point mutant that was more suitable for complex structure determination (Bick et al., 2017)) was placed into the 2QZ3 scaffold using PyMOL. Fentanyl was placed into the 2QZ3 scaffold by using PyMOL to align 2QZ3 and 5TZO, then combining the fentanyl molecule from 5TZO and the protein structure from 2QZ3 into a new PDB file. While all positions of Fen49 were subjected to SSM, for our study we designed only the 18 residues defined as binding site in (Bick et al., 2017) (**Table S7**). Design was allowed only to those amino acids with high enough sequencing counts to be included in the enrichment and depletion calculations in (Bick et al., 2017) (**Table S8**). Finally, four positions (37, 64, 69, 71) in the input structure were set to alanine (using Rosetta’s FixBB protocol), because the wild-type residue was disallowed due to low counts (**Table S8**). In CoupledMoves design, 22 positions were allowed to repack based on the possibility of clashes with designed positions; in design by non-coupled methods, all positions were allowed to repack (**Table S7**). The known profile represents the frequency equivalent (*F_equiv_*, described below) of the final round of selection (obtained from the authors). Note that the experimental screen mutated one position at a time, whereas we design multiple positions simultaneously.

#### Frequency equivalent

Experimental data from the DIG10 and Fen49 datasets are deep sequencing counts before and after selection. To allow direct comparison between the experimental data and sequence profiles from Rosetta design for each mutation *x* at position *i*, we derived a frequency equivalent *F_equiv_* from the experimental frequency data:

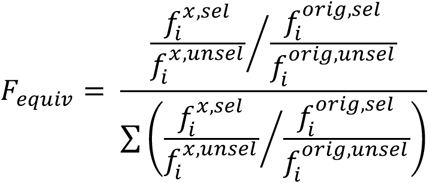

where 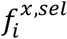 and 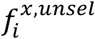 are the frequency of that mutation, and 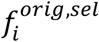 and 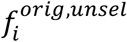 are the frequency of the original amino acid identity, in the selected and unselected populations, respectively, and *F_equiv_* is normalized by dividing over the sum across all amino acid identities found in the sequencing results. *F_equiv_* is then used in comparison to Rosetta design results.

#### hGH/hGHR

The hGH/hGHR dataset was taken from (Humphris & Kortemme, 2008). The protein-protein interface between human growth hormone (hGH) and human growth hormone receptor (hGHR) is high affinity, with a K_D_ reported as 0.9 nM (Cunningham & Wells, 1993) and 1.56 nM (Pal et al., 2006). As input for design, we used a crystal structure (PDB: 1A22). The known sequence profiles were taken from a phage display selection experiment, wherein 35 key residues from the ∼1300 Å^2^ hGH/hGHR interface were divided into six combinatorial libraries of five or six positions (Pal et al., 2006). To minimize potential cooperative interactions, positions were grouped into libraries that maximized the three-dimensional distance between residues. Our computational workflow mimicked this strategy, using the same designable residues and running independent design trajectories for each of the six libraries. As in (Humphris & Kortemme, 2008), residues within 4 Å of designed residues were allowed to repack (**Table S9**).

#### Herceptin-HER2

The Herceptin-HER2 dataset was taken from (Babor et al., 2011). The protein-protein interface between therapeutic antibody Herceptin and its target, human epidermal growth factor 2 (HER2), is high affinity (KD = 0.35nM (Gerstner et al., 2002)). We used a crystal structure (PDB: 1N8Z) as input structure for design, truncated as in (Babor et al., 2011) to include only chain A positions 1-106, chain B positions 1-119, and chain C positions 511-607. The known sequence profiles were taken from phage display selection experiments that used five combinatorial libraries containing five to seven positions each after four rounds of selection (Gerstner et al., 2002). We mimicked the experimental strategy in our computation, with five separate design runs, one for each experimental library, and allowing repacking of residues within 4 Å of designed residues, as in (Babor et al., 2011) (**Table S10**). Herceptin/HER2 sidechains were repacked from the crystal structure before design.

### Performance metrics

#### Position profile similarity

Position profile similarity (PPS) was computed as described in (Ollikainen et al., 2015). Briefly, PPS represents the similarity in the side chain amino acid identity distributions between the predicted and known sequences at a given position:

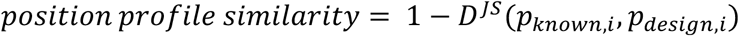

where *p_known,i_* and *p_design,i_* are the probability distributions over the 20 amino acids for the known (natural or experimental) and designed sequences, respectively, at position *i* and *D^js^*(*x*, *y*) is the Jensen-Shannon divergence between two distributions *x* and *y*, as in.

#### RankTop

For the profile datasets, mutations were ranked according to their frequency in the predicted and known (experimental/natural) sequence profile. RankTop is the rank, in the predicted profile, of the top ranked amino acid from the known profile. If the amino acid is not found, its rank is set to 20.

#### Percent Enrichment

As in (Ollikainen et al., 2015) the percent enrichment (PE) for each specificity-altering mutation in the enzyme specificity dataset was calculated as follows:

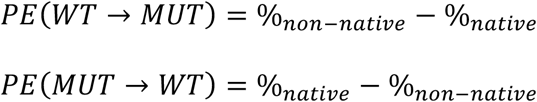

where %*_native_* is the percent occurrence of the mutation in sequences designed for the native ligand and %*_non-native_* is the percent occurrence of the mutation in sequence designed for the non-native ligand. *PE*(*WT* → *MUT*) was used for predictions that start with the wild-type structure and *PE*(*MUT* → *WT*) was used for predictions that start with the mutant structure. As in (Ollikainen et al., 2015), a prediction was considered correct if it obtained a positive percent enrichment value.

#### Rank

For the enzyme specificity dataset, mutations were ranked by descending order of their percent enrichment values, as described in (Ollikainen et al., 2015).

#### Sequence Entropy

The sequence entropy *H_i_* was computed as in (Ollikainen et al., 2015):

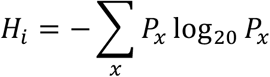

where *P_x_* is the percent of sequences with amino acid *x* at position *i*.

#### Distance from input sequence

Distance from input sequence is a variation of the profile similarity metric, where distance is calculated as:

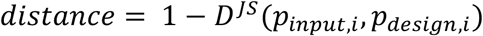

where *p_input,i_* and *p_design,i_* are the probability distributions of the single input side chain and the designed sequence profiles, respectively, at position *i*.

## RESULTS

### Design methods

We set out to compare four flexible-backbone design methods (**Figure 1, Methods**) using a common set of benchmarks (described below): (i) FastDesign utilizing the Rosetta FastRelax method (Khatib et al., 2011; Tyka et al., 2011) for backbone flexibility, (ii) BackrubEnsemble Design (Smith & Kortemme, 2010, 2011), (iii) CoupledMoves with Backrub (CM-BR) (Ollikainen et al., 2015), and (iv) the new CoupledMoves with Kinematic Closure (CM-KIC) method introduced here. We also compare to fixed-backbone design (FixBB) and a null model where all amino acid frequencies are set to 5%.

**Figure 1.**
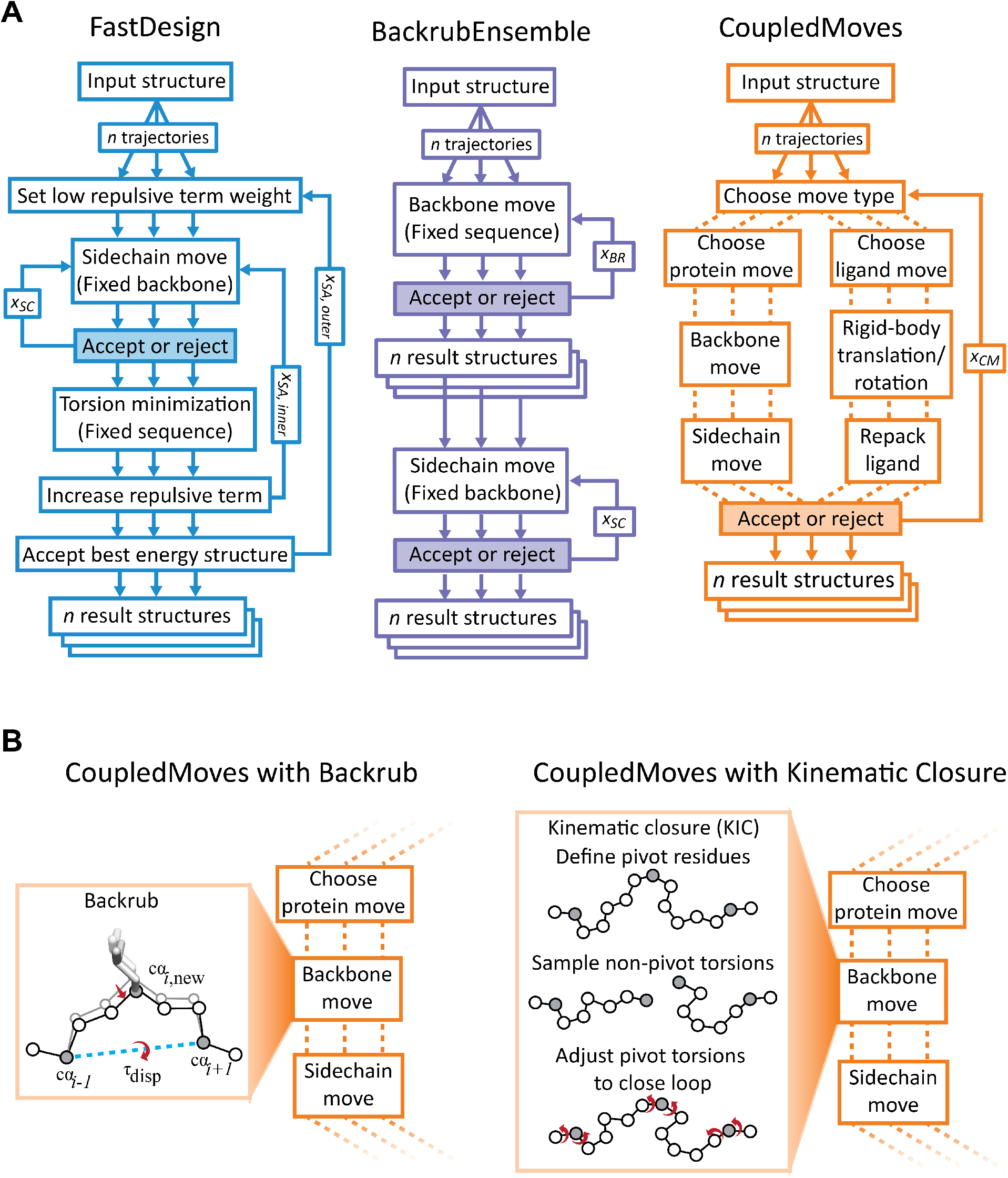
Design methods. **(A)** Design method comparison. The FastDesign (left, blue) and BackrubEnsemble (middle, purple) methods separate sequence design steps (using a fixed backbone) from backbone optimization steps (using a fixed sequence). CoupledMoves (right, orange) evaluates combined moves that sample both backbone conformation and amino acid sequence (or, alternatively, combine ligand translations/rotations with changes of ligand conformers). FastDesign performs five outer (*x_SA,outer_*) simulated annealing cycles, during which the weight of the Lennard-Jones repulsive energy term is ramped from 2% to 100%. For each ramped weight, an inner cycle (*x_SA,inner_*) consists of a complete round of sequence design with *x_SC_* steps on a fixed backbone, followed by a step that minimizes backbone, sidechain, and ligand torsion angles. BackrubEnsemble performs 10,000 (*x_BR_*) Backrub moves to generate each ensemble member. For both FastDesign and BackrubEnsemble, *x_SC_* scales with the number of possible moves, and is equal to 10 times the number of possible rotamers at all designable or repackable positions. CoupledMoves performs 1000 trials (*x_CM_*) per trajectory. **(B)** Original and updated backbone mover in CoupledMoves. The original CoupledMoves method (Ollikainen et al., 2015) uses the Backrub algorithm to make backbone moves. A backrub move (Davis et al., 2006; Smith & Kortemme, 2008) rotates a segment as a rigid body by displacement angle τ_disp_ around an axis between two pivot Cα atoms 2-11 residues apart (shown is a 2-residue move). In the updated versions of CoupledMoves introduced here, backbone moves are made using a Kinematic Closure algorithm (Mandell et al., 2009). Backbone torsion angles for non-pivot Cα atoms are perturbed either using fragment insertion (FKIC) or by small perturbations away from the existing angles (WKIC), then the loop is closed by analytical determination of Φ and Ψ angles (red) at three pivot Cα atoms (grey).

The main algorithmic differences between the methods are illustrated in **Figure 1A**. FastDesign (**Fig. 1A, left**) iterates between two steps. In the first step (fixed-backbone sequence design), amino acid side chain identities and rotameric conformations are optimized using Monte-Carlo simulated annealing but the backbone is kept fixed. In the second step (fixed-sequence torsion minimization), the entire structure is minimized using backbone and side chain torsion degrees of freedom while keeping the sequence fixed. These steps are iterated through cycles of simulated annealing, during which the weight of the repulsive component of the Lennard-Jones potential is increased stepwise to enable amino acid changes that may introduce unfavorable clashes but can be subsequently relaxed in the minimization step. FastDesign has been used in a variety of application to design new functions (Chevalier et al., 2017; Dou et al., 2018; Silva et al., 2019).

The BackrubEnsemble method (Smith & Kortemme, 2010, 2011) (**Figure 1A, middle**) also proceeds in two steps. The first step generates an ensemble of backbones through application of Backrub moves. Each Backrub move (Smith & Kortemme, 2008) selects two pivot backbone Cα atoms and rotates the entire segment between them (2-11 residues) as a rigid body. Backrub moves are made throughout the protein structure (or a predefined region) by randomly selecting pivot points. The second step performs fixed-backbone sequence design on each member of the ensemble using Monte-Carlo simulated annealing. The BackrubEnsemble method in Rosetta has been shown to recapitulate protein conformational fluctuations (Friedland et al., 2009; Friedland et al., 2008; Ollikainen et al., 2013) and tolerated sequence space (Humphris & Kortemme, 2008; Smith & Kortemme, 2010, 2011) and has been successfully applied to the redesign of protein recognition specificity (Kapp et al., 2012).

In contrast to FastDesign and BackrubEnsemble that separate fixed-backbone sequence design from fixed-sequence backbone sampling, CoupledMoves combines backbone and side chain moves, which can include sequence changes, into a single “coupled” Monte-Carlo step (**Figure 1A, right**). In this fashion, the backbone can respond to a designed sequence change more directly than in the non-coupled FastDesign and BackrubEnsemble methods. However, coupling backbone and side chain moves could artificially collapse designed structures. Because replacing a larger with a smaller amino acid side chain is less likely to lead to clashes, the change is more likely to be accepted. In subsequent steps it is harder to recover from such a collapse as the backbone will have moved to accommodate the smaller side chain. To alleviate this problem, each side chain move in CoupledMoves considers all rotamers for allowed amino acids and chooses a likely side-chain rotamer and identity based on its Boltzmann-weighted Rosetta score. This change led to a considerable decrease in the number designed alanine or glycine side chains (Ollikainen et al., 2015). Finally, coupled moves can also be performed for the ligand, where rotation and translation of the ligand can be combined with ligand conformer changes. Coupled moves has been shown to better recapitulate amino acid preferences in small molecule binding sites and mutations that switch enzyme specificity (Ollikainen et al., 2015), but has not yet been tested in a forward-engineering application.

The original version of the Coupled moves method uses Backrub moves to sample backbone degrees of freedom. Here we introduce an updated version of the CoupledMoves algorithm that performs backbone moves with the kinematic closure (KIC) algorithm (Mandell et al., 2009) (**Figure 1B**). KIC selects two pivot Cα atoms that define a segment, and a third pivot Cα atom within the segment. The algorithm next perturbs the backbone torsion angles around all non-pivot Cα atoms in the segment, breaking the loop. Finally, the torsion angles of the three pivot atoms are solved analytically to close the loop. The original implementation of KIC samples backbone phi/psi torsion angles at the non-pivot Cα atoms probabilistically from Ramachandran space (Mandell et al., 2009). Our implementation here allows phi/psi sampling by substitution of peptide fragments derived from the protein structure databank (FKIC) or random “walk” perturbation of backbone torsion angles by values from a Gaussian distribution centered around zero with a standard deviation of 3° (WKIC) (see Methods).

### Benchmark datasets

We evaluate the performance of the different methods on six benchmark datasets (**Table 1**, **Figure 2**). Each benchmark contains information on functional sequence variants. We chose binding as a proxy for function because the engineering of binding interactions is a common task with many important applications. Moreover, the stability of a binding interaction is a functional constraint that can be more easily explicitly modeled and scored by Rosetta than for example requirements for efficient enzyme catalysis that are often incompletely understood. The datasets comprise both small molecule binding sites and protein-protein interaction interfaces.

**Figure 2.**
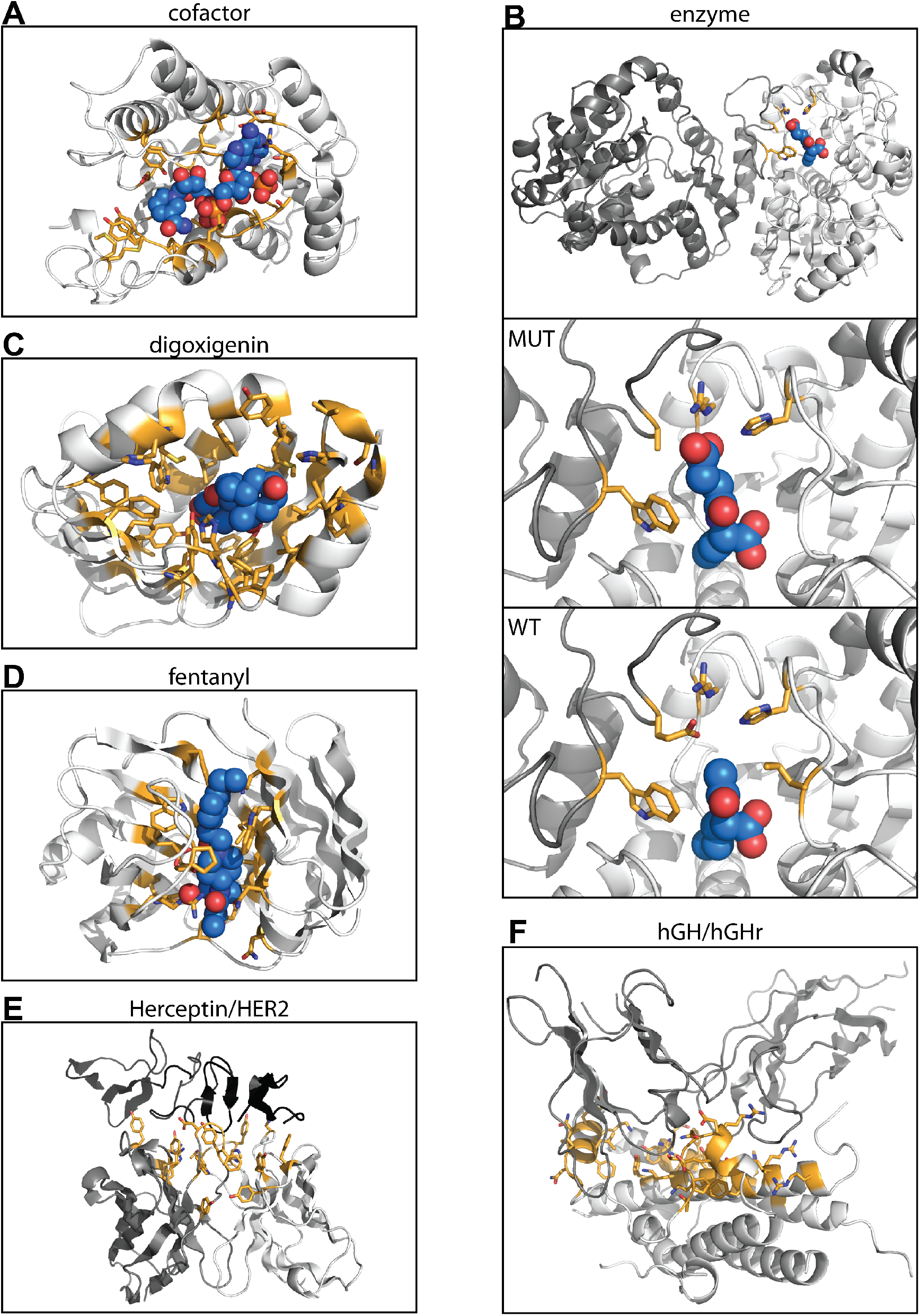
Benchmark dataset structures. Side chains at designed positions are highlighted in orange and shown as sticks. Ligands are colored light blue and shown in sphere representation. The structures shown are those used as input for design, as described in Methods. Nitrogen atoms are shown in dark blue, and oxygen atoms are shown in red. (**A**) Representative structure from the cofactor dataset, Alcohol Dehydrogenase with cofactor NAP. Structures for other six protein families are not shown. (**B**) Representative structure from the enzyme dataset, N-acetylornithine carbamoyltransferase. The full structure of the mutant enzyme, with ligand N-(3-carboxypropanoyl-L-norvaline (SN0), is shown in the left panel. The middle panel shows the binding site. The right panel shows the binding site of the wild-type protein, with ligand N-acetyl-L-norvaline (AN0). The other nine enzymes are not shown. (**C**) DIG10.1, the designed digoxigenin binder on which the SSM library was generated and selected, with digoxigenin. (**D**) The wild-type protein used for design of fentanyl binding protein, with fentanyl placed in the binding pocket. (**E**) Herceptin/HER2. Designable positions on the Herceptin antibody light chain (light gray) and heavy chain (dark gray) interact with target HER2 (black). The combination of designable positions from all libraries are shown. (**F**) hGH/hGHr. Designable positions on hGH (light gray) interact with target hGHr. The combination of designable positions from all libraries is shown.

Four of the datasets contain small molecule binding sites (**Table 1**, **Figure 2**). The first two datasets were taken from (Ollikainen et al., 2015). Dataset 1 comprises evolutionary sequence alignments for seven naturally occurring protein families (**Figure 2A** shows one structure as a representative) that each bind a specific cofactor (“cofactor” set, **Tables S1-S2**). Dataset 2 was curated from experimentally-characterized substrate specificity-altering point mutations for ten different enzymes (“enzyme specificity” set, **Figure 2B, Table S3-S4**). Datasets 3 and 4 were compiled from site saturation mutagenesis (SSM) experiments performed on two different proteins designed by Rosetta to bind small molecules (sets “DIG10” (digoxigenin) (Tinberg et al., 2013), **Figure 2C** and **Table S5-S6**, and “Fen49” (fentanyl) (Bick et al., 2017), **Figure 2D** and **Table S7-S8**). The SSM libraries were screened for binding to the target small molecule (digoxigenin or fentanyl, respectively) using yeast display followed by deep sequencing of naive and selected populations.

**Table 1.**
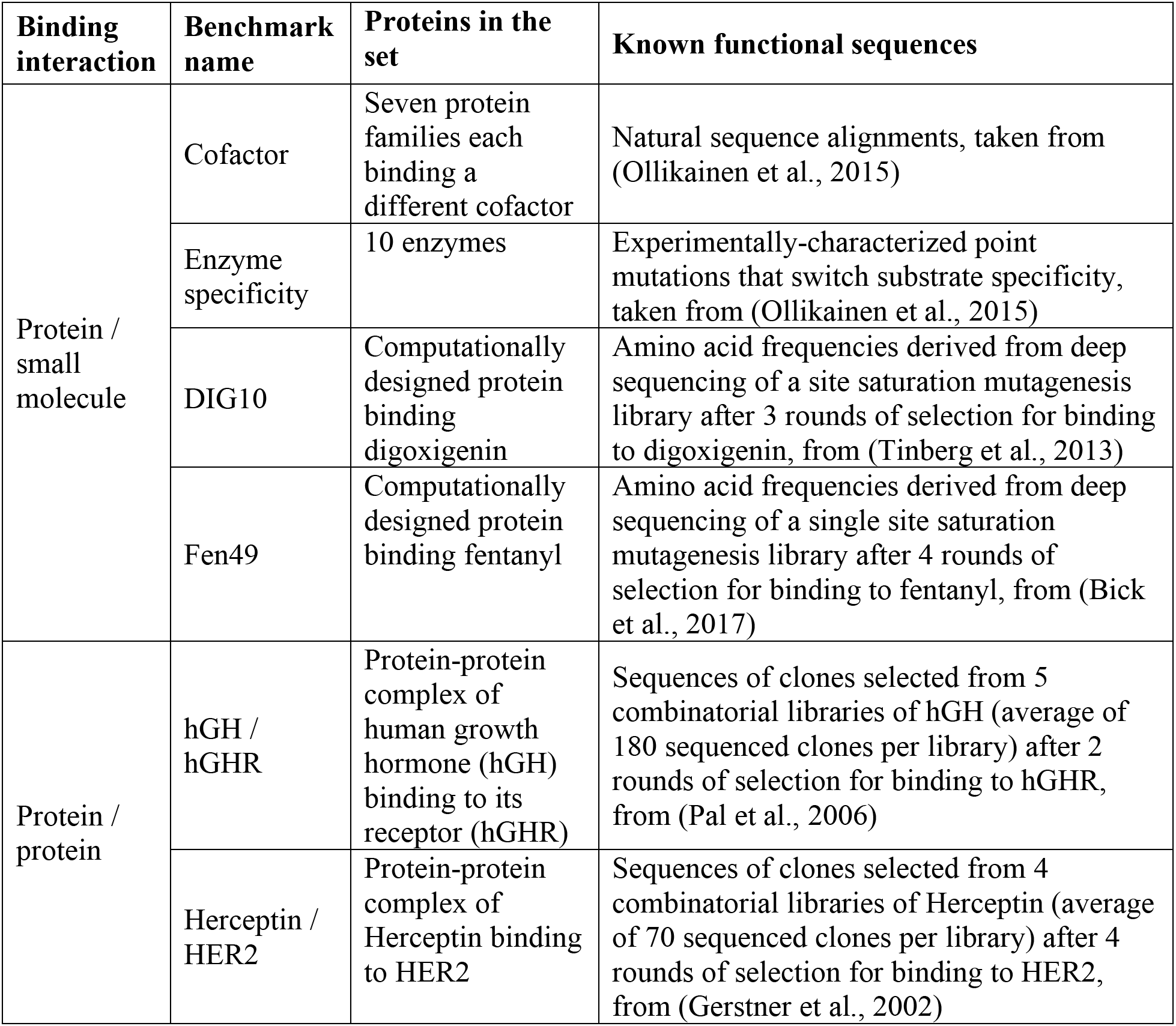
Benchmark datasets.

The two protein-protein interface datasets contain sequences selected from combinatorial libraries (allowing all 20 naturally occurring amino acids at 5 to 7 sequence positions) by phage display and subsequent sequencing of individual clones (**Table 1**). Dataset 5 comprises sequences from 5 phage display libraries of Herceptin (17 positions total) selected for binding to HER2 (“Herceptin/HER2” set (Gerstner et al., 2002), **Figure 2E** and **Table S9**). Dataset 6 comprises sequences from 6 libraries of human growth hormone (hGH) (35 positions total) selected for binding to human growth hormone reception (hGHR) (“hGH/hGHR” set (Pal et al., 2006), **Figure 2F** and **Table S10**).

### Performance metrics

Five of the datasets contain sequences from either experimental selection (DIG10, Fen49, Herceptin-HER2, hGH/hGHR) or natural sequence alignments of evolutionary families (cofactor), reflecting the diversity of amino acids at each position compatible with the protein’s function (tolerated sequence space) (Humphris & Kortemme, 2008). We refer to this diversity as the “known sequence profile” for each position. We evaluate the ability of our design methods to recapitulate these known sequence profiles by quantifying two metrics used previously (Ollikainen et al., 2015; Smith & Kortemme, 2010), profile similarity and rank top, both calculated per position (see Methods). Position profile similarity (PPS) measures the similarity of the probability distribution of amino acid frequencies between the known profile and the profile generated by Rosetta design at each position. Rank top measures the rank, in the design profile, of the amino acid most frequently observed at a given position in the known profile.

The enzyme specificity benchmark (Ollikainen et al., 2015) contains individual point mutations (rather than sequence profiles) experimentally characterized to switch enzyme substrate specificity. In this case, in contrast to the analysis for the sequence profile datasets, we do not assume knowledge of positions mutated in the experiment. Instead, we evaluate how the experimentally characterized specificity switching mutation ranks across designed mutations at all positions in the vicinity of the changed substrate, to approximate an actual design project where it is not clear a priori which position should be mutated. In addition to the absolute rank we also evaluate the percentile (Ollikainen et al., 2015), of the experimentally characterized mutation among all design predictions (see Methods).

Each metric has a different experimental interpretation. The tolerated sequence space captured by the PPS metric is useful for the design of libraries, which can be screened for criteria in addition to binding affinity and specificity, such as protein stability and solubility. RankTop is useful for cases where a few mutations or design sequences are selected for individual experimental tests. Percentile gives information on how many predictions would need to be tested in order to find a successful mutation when making predictions for a range of positions.

### CoupledMoves improves prediction of tolerated sequence space

We first evaluated the overall performance of each flexible backbone design method on the five sequence profile datasets. **Figure 3A** shows the distributions of position profile similarities across all designed positions in each benchmark, with the median indicated by a white dot. CoupledMoves attains higher median PPS values than FastDesign and FixBB for all datasets, and higher median PPS values than BackrubEnsemble for all datasets except Fen49 and DIG10 where Coupled Moves and BackrubEnsemble perform similarly. Somewhat surprisingly, using this global metric CoupledMoves is worse than the null model for the Fen49 set. Even more strikingly, FastDesign and FixBB do not attain a higher median PPS values than the null model for any of the datasets (except cofactor), and are considerably worse than the null model for the hGH/hGHR and DIG10 datasets. As discussed below, the comparatively poor overall PPS of all methods for the hGH/hGHR, DIG10, and Fen49 datasets is due to low similarity between the input sequence and the known profile. In these cases, the null model scores as well or better than the design methods; of the flexible-backbone design methods, CoupledMoves performs best.

**Figure 3.**
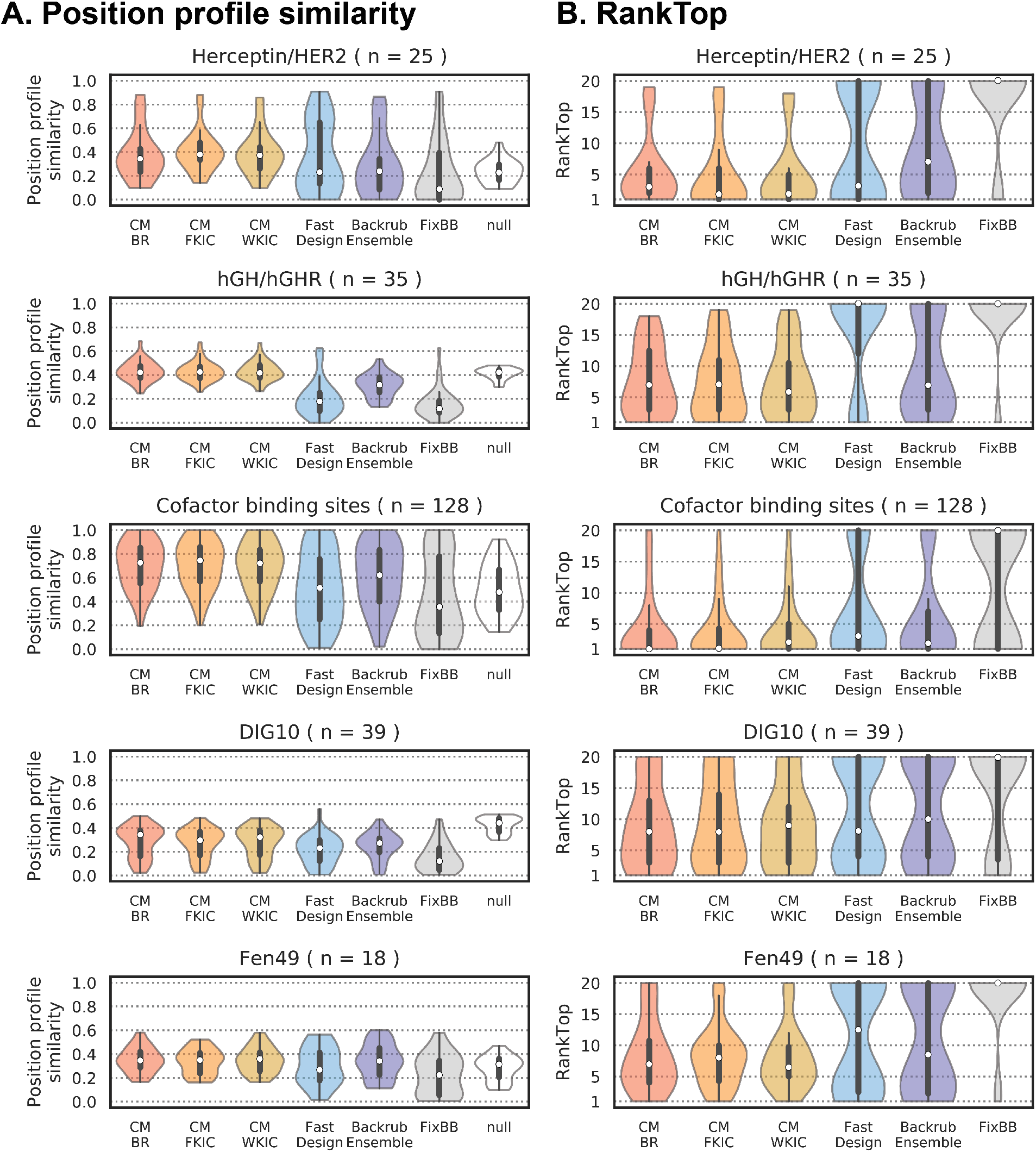
Comparison of design method performance on sequence profile datasets. PPS (**A**) and RankTop (**B**) distributions. A rank of 1 means that the design method correctly identified the most frequent amino acid side chain observed in the experimental/natural profile, whereas a RankTop of 20 means that the side chain was observed with zero frequency, or that all other side chains were modeled with some frequency and the top known side chain was the least frequent. The median of the distributions is marked with a white dot. Second and third quartiles are marked by the thick black bar, and the thin bar marks 1.5 times the inter-quartile range. The width of the violins is determined by the number of observations in each bin, and bins are defined using Scott’s normal reference rule. The number of sequence positions in each set is described by n.

We next evaluated the RankTop values for all five datasets (**Figure 3B**). Here, all flexible backbone methods (except FastDesign for the hGH/hGHR dataset) perform better than fixed backbone design, which in the majority of the cases misses the most frequent amino acid side chain from the known profiles (the null model by definition ranks all amino acids the same so is not relevant here). The rank top values are lowest (best) for the Herceptin/HER2 and cofactor sets. CoupledMoves performs better than BackrubEnsemble and FastDesign for the Herceptin/HER2, hGH/hGHR and cofactor datasets, similar to FastDesign for the Dig10 set and similar to BackrubEnsemble for the Fen49 set. Moreover, for several benchmarks (hGH/hGHR, Herceptin/HER2, Fen49), CM-WKIC leads to small but noticeable improvement in RankTop values over CM-BR. Taken together, when considering both PPS and RankTop over all datasets, CoupledMoves and in particular CM-WKIC perform best overall.

We also considered PPS and RankTop for each protein family comprising the Cofactor dataset (**Figure S1**), and found that CoupledMoves outperforms FastDesign for all families, and outperforms BackrubEnsemble for six of the seven families, with the exceptions of the flavin binding site of Flavodoxins. Performance for individual libraries of the Herceptin/HER2 (**Figure S2**) and hGH/hGHR (**Figure S3**) leads to similar conclusions.

To determine if methods were more predictive for different groups of positions, we plotted the PPS values for the different methods against each other (**Figure 4A,B**). CoupledMoves achieves similar or better PPS for nearly all positions when compared to the non-coupled methods (**Figure 4A**, CM-KIC shown as example). BackrubEnsemble achieves PPS values better or similar than FastDesign (**Figure 4B, left**), and better than FixBB (**Figure 4B, middle**), for almost all positions. FastDesign, compared to FixBB, achieves better PPS for some positions, but worse PPS for others (**Figure 4B, right**). **Figure 4C** quantifies the number of positions for which CoupledMoves is better, worse, or similar to the non-coupled methods. A prediction for a position is classified as “better” or “worse” by a given method relative to a comparison method when the difference in performance is above or below, respectively, a threshold of ± 0.1 for PPS or ± 5 for RankTop. When the difference is within the threshold, the predictions are classed as “similar.” CoupledMoves achieves better PPS values than BackrubEnsemble for 65 ± 1 positions, better than FastDesign for 119 ± 2 positions, and better than FixBB for 143 ± 2 positions. Standard deviation represents the average across CM-BR, CM-FKIC, and CM-WKIC. CoupledMoves also achieves better RankTop for more positions than BackrubEnsemble, FastDesign, and FixBB (39 ± 3, 67 ± 3 and 126 ± 4 positions, respectively), (**Figure 4C**). Moreover, CoupledMoves performs worse than non-coupled methods for very few positions (**Figure 4C**, red bars).

**Figure 4.**
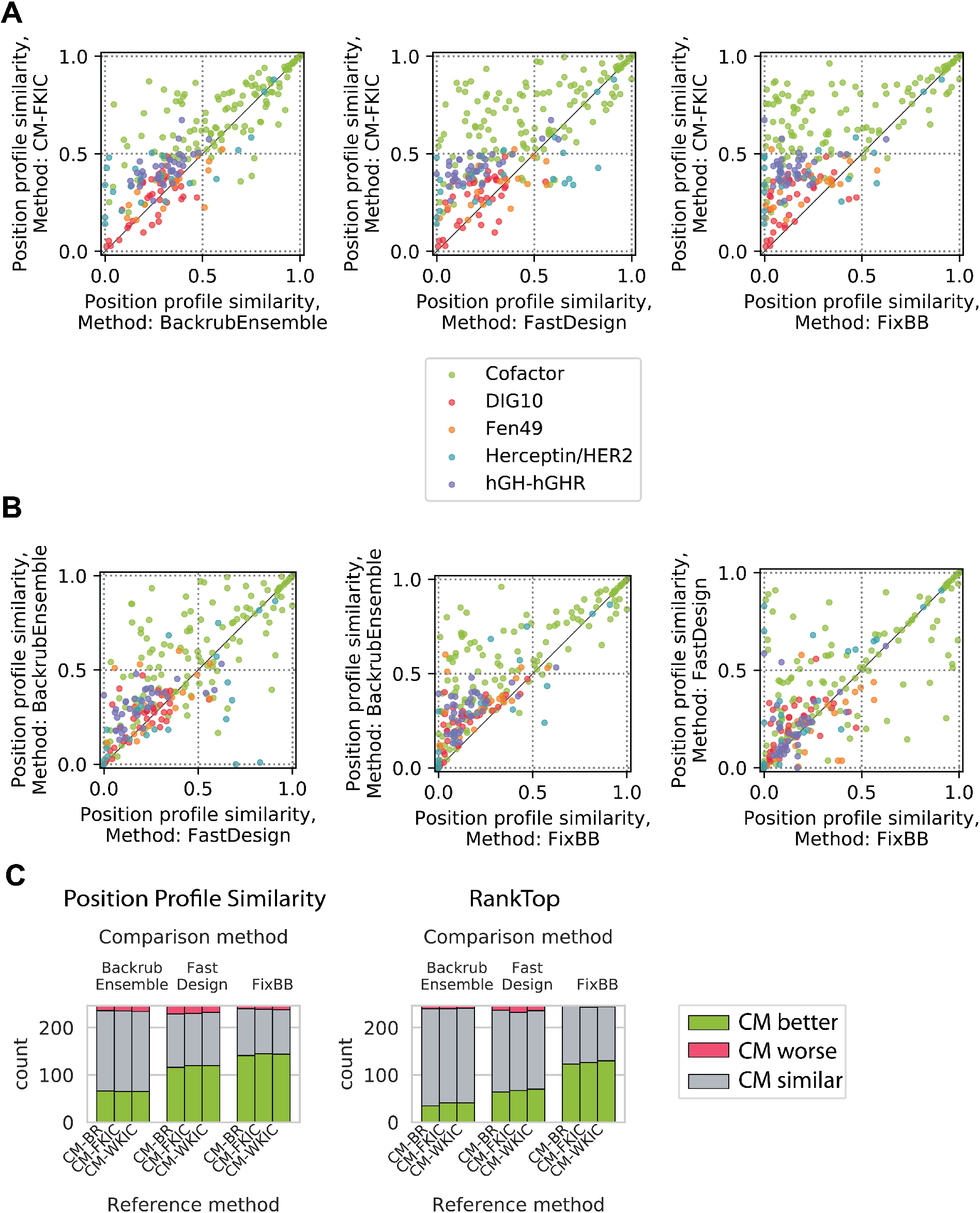
Method performance comparison for profile datasets by sequence position. Shown are the same data as in Figure 3, but plotting individual sequence positions instead of distributions. Colors indicate different datasets. (**A**) Comparison between CM-FKIC and non-coupled methods. Points above the diagonal represent positions where CM-FKIC outperforms the non-coupled method. (**B**) Comparison between non-coupled methods, where points above the diagonal represent positions where BackrubEnsemble outperforms FastDesign (left) or FixBB (middle), or where FastDesign outperforms FixBB (right). (**C**) Summary of position counts classified by whether CoupledMoves (“Reference method”) performs better (green), worse (red) or similar (gray) compared to coupled methods (“Comparison method”). The CoupledMoves reference method is “better” or “worse” than the comparison method when the difference in performance is above or below, respectively, a threshold of ± 0.1 for PPS or ± 5 for RankTop. When the difference is within the threshold, the methods are classed as “similar.”

### CoupledMoves outperforms the other methods at recapitulating key affinity-determining side chains

We next sought to evaluate the ability of the different methods to predict amino acid preferences for the positions that are most functionally important in the 5 profile datasets. Sequence logo representations of the tolerated sequence space for each of our datasets (**Figures S4-S8**) indicated considerable differences in sequence entropies between individual positions, and we reasoned that conserved side chain residues at low sequence-entropy positions are more likely to be important for protein function than residues at position with higher entropy. We hence split the positions in each dataset into three sequence entropy groups (see Methods) and evaluated median PPS and RankTop for the cofactor and Herceptin/HER2 datasets, which have the most consensus positions (**Figure 5, Figure S9**). Positions with low (entropy ≤ 0.33) or medium (0.33 < entropy ≤ 0.67) entropy were defined as consensus positions. The top known side chain for these positions was defined as the consensus side chain. We find that CoupledMoves achieves better PPS than the null model for consensus positions in the Herceptin/HER2 and cofactor datasets. FastDesign is better than the null model for only low-entropy positions for both datasets. BackrubEnsemble is better than the null model for low entropy positions in the cofactor dataset, but not Herceptin/HER2. In contrast, the null model has the highest PPS for the high entropy bin, which might be expected for positions with high mutational tolerance.

**Figure 5.**
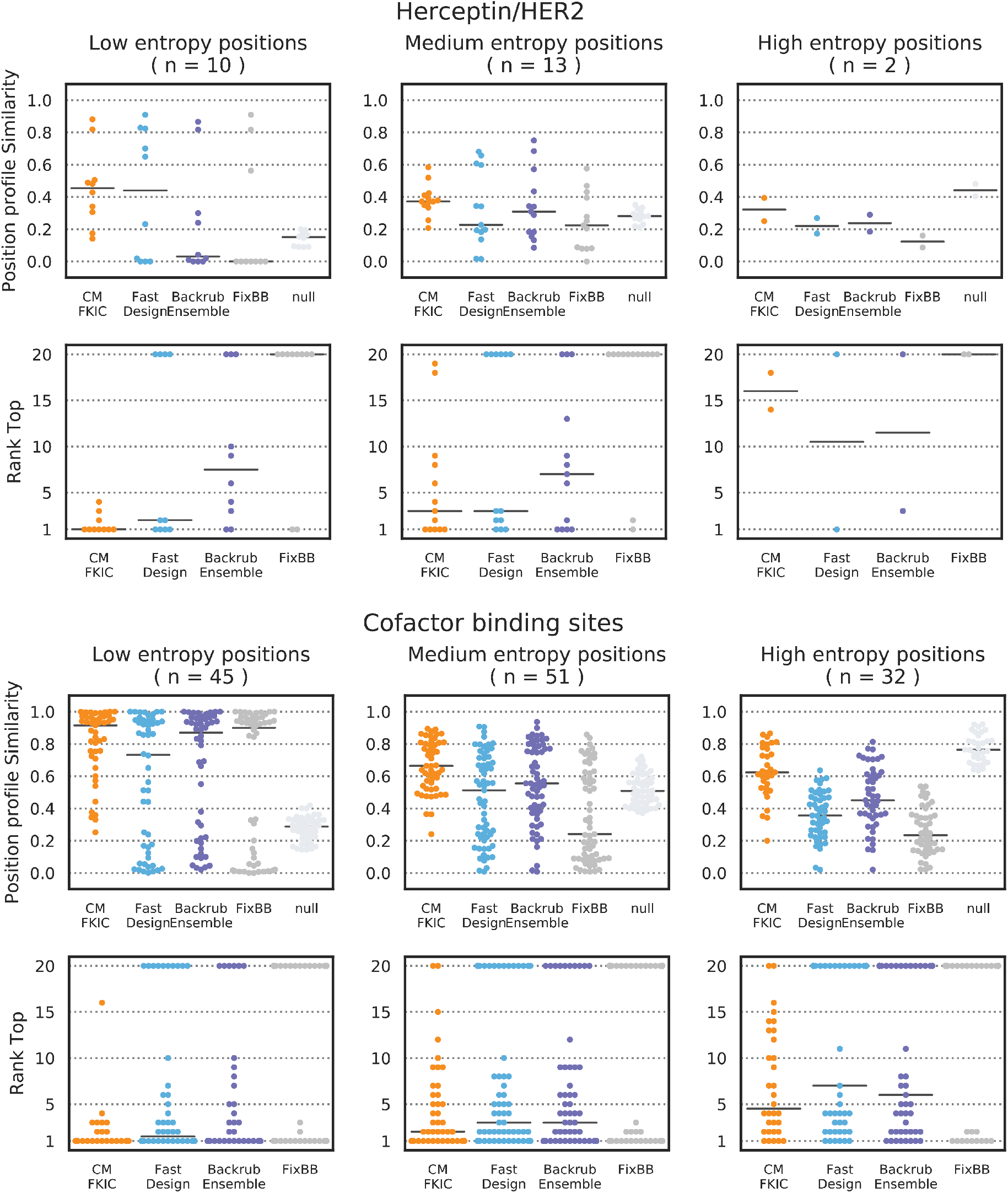
PPS and RankTop as a function of known sequence entropy. Each point represents one sequence position. Shown here are the Herceptin/HER2 (top) and Cofactor (bottom) datasets, which have the highest number of medium and low entropy positions. The remaining datasets are shown in **Figure S9**. For each dataset, PPS and RankTop are binned by entropy of the known sequence profile at each position (low: entropy ≤ 0.33, medium: 0.33 < entropy ≤ 0.67, and high: entropy > 0.67). The median is marked with a horizontal black line.

Similar to PPS, CoupledMoves achieves the best (lowest) RankTop values for consensus positions, predicting the correct amino acid residue with at least some frequency at most positions, as opposed to non-coupled methods which frequently do not identify the consensus side amino acid identity at all (rank of 20) (**Figure 5**). CoupledMoves predictions typically have the highest entropy (**Figure S10**), which leads on average to higher similarity at variable positions. Nevertheless, PPS and RankTop at low-entropy positions (**Figure 5, Figure S9**), and energetic rankings of consensus positions (see Discussion) indicates that CoupledMoves is the most accurate method for functionally relevant interactions.

In addition to low-entropy positions determined from known sequence profiles, we also considered experimentally-characterized affinity-improving mutations, which were available for the Herceptin/HER2, Fen49, and enzyme specificity datasets (the latter set is discussed below). For Herceptin, the most important affinity-improving mutation, D VH98 W, resulted in 3-fold improvement of binding affinity and was found in 23% of sequences resulting from phage display (Gerstner et al., 2002). Contrary to previous findings (Babor et al., 2011) where BackrubEnsemble recapitulated D VH98 W as the top mutation, the non-coupled methods tested in this study did not identify tryptophan (**Figure S7**), but CoupledMoves methods selected the tryptophan mutation at low frequency (CM-BR 1.1%, CM-FKIC 1.3%, CM-WKIC 1.5%). We note that this position is surface exposed in the original structure, leading to high entropy in the design profiles where many side chains are tolerated. It is possible that a structural rearrangement in the D VH98 W mutant adds additional interactions across the interface but that these structural changes are not correctly modeled in our simulations.

For the Fen49 dataset, the authors identified two key mutations, A77V and A171I, that led to ∼100-fold improvement in binding affinity to fentanyl, but none of the design methods tested here found both mutations (**Figure S6**). These two positions are located in the binding pocket and enriched in larger hydrophobic residues in the selection, presumably to provide additional surface complementarity with fentanyl (Bick et al., 2017). While all design methods did substitute larger hydrophobic side chains, only FastDesign ranked 171I highly, and only BackrubEnsemble ranked 77V highly. CoupledMoves selected 77V at a lower frequency. No method identified the combination of A77V and A171I. While there is no crystal structure with these mutations, we hypothesize that packing I171 against the phenyl ring of fentanyl may be inaccessible to the fentanyl conformer of Fen49, and modeling ligand flexibility might enable design to converge on I171. Unlike position 171, which is an ideal distance for van der Waals interactions with fentanyl, there is an almost 6 Å distance between the closest heavy atoms of position 77 and the ligand, and the position has a relatively large solvent-accessible surface area. It is therefore unsurprising that Rosetta is unable to arrive at a consensus for this position. The inability of all methods to find the key mutations in Fen49 may represent shortcomings in modeling ligand flexibility. In addition, the Fen49 deep sequencing results are incomplete due to experimental limitations (for example, the original Fen49 side chains were present in the selection but did not have frequency counts (Bick et al., 2017)).

### CoupledMoves improves prediction of substrate specificity-altering mutations

The Enzyme Specificity dataset provides an opportunity to analyze functionally important mutations, as the dataset is made up of pairs of structures where individual point mutations have been experimentally characterized that switch ligand-binding specificity between two ligands (Ollikainen et al., 2015). To determine to what extent the different flexible backbone methods can recapitulate these experimentally characterized specificity-switching mutations, we carried out design simulations on structures with either the original or the new ligand in the binding pocket and designing positions in the vicinity of the ligand substructure change, as described previously (Ollikainen et al., 2015) (**Table S3, Table S4**). To design for mutations switching specificity to the new ligand, we prepared the input structure by computationally substituting the new ligand into the binding pocket of the wild-type protein crystal structure. For the inverse, we swapped the wild-type ligand into the binding pocket of the mutant crystal structure (see Methods).

Some enzymes in this dataset have multiple experimentally-characterized mutations, either a single position to multiple identities (Protein Data Bank (PDB) codes: 1K70, 3KZO), or multiple positions (PDB: 1A80, 3HG5), for a total of 29 cases (12 wild-type and 17 mutant side chains). The CoupledMoves methods (CM-BR, CM-FKIC, CM-WKIC) correctly identify (positive percent enrichment, see Methods) 14, 11, and 12 mutations specificity-determining mutations, respectively, while the non-coupled methods (FastDesign, BackrubEnsemble, FixBB) identify only 7, 7, and 5 mutations, respectively (**Table 2**). All CoupledMoves methods identify specificity-altering mutations with a better percentile and rank than the non-coupled methods (**Tables 2, 3**), with the original CM-BR attaining the best median and quartile performance, and FastDesign and BackrubEnsemble performing similarly poorly.

**Table 2.** Performance summary for the enzyme specificity benchmark. Shown are median values across the benchmark. Percentile and rank were computed from sorted percent enrichment values (as defined in Methods). If the correct amino acid is not sampled, rank is the maximum possible number of amino acids for designed positions (number of designable positions*20). A mutation is considered correctly identified if a method identifies it with a positive percent enrichment.

|  | Find mutant amino acid starting from wild-type structure & mutant ligand |  |  | Find wild-type amino acid starting from mutant structure & wild-type ligand |  |  |
| --- | --- | --- | --- | --- | --- | --- |
|  | Median Percentile | Median Rank | Number of mutations identified | Median Percentile | Median Rank | Number of mutations identified |
| CM-BR | 79 | 15 | 14 | 76 | 22 | 13 |
| CM-FKIC | 70 | 25 | 10 | 76 | 22 | 13 |
| CM-WKIC | 78 | 19 | 12 | 80 | 22 | 12 |
| BackrubEnsemble | 0 | 60 | 7 | 74 | 32 | 9 |
| FastDesign | 0 | 60 | 7 | 0 | 100 | 4 |
| FixBB | 0 | 80 | 5 | 0 | 80 | 5 |

### Non-coupled methods more frequently make incorrect predictions where correct side chain residues are lost, while CoupledMoves most frequently makes predictions where correct residues are gained

We next considered how the sequence of the input structure influences method performance. Only positions with low and medium entropy (≤ 0.67) in the known profile are considered. Three broad scenarios can be distinguished (**Figure 6, top panels**). In the first scenario (“loss”), the input side chain (the residue in the starting structure used for design) is present or even preferred in the known sequence profile but is depleted in the design simulations. In the second scenario (“gain”), the input side chain and the known position profile are dissimilar, but preferred side chains are enriched by design. The third scenario occurs when design results in little change of similarity to the known profile (“neutral”). When plotting the PPS values for each method as a function of profile similarity to the input, loss occurs more frequently for positions designed by BackrubEnsemble, FastDesign and FixBB, whereas gain occurs more frequently for positions designed by CoupledMoves and BackrubEnsemble (**Figure 6a**, middle and bottom panels, **Table S11**).

**Figure 6.**
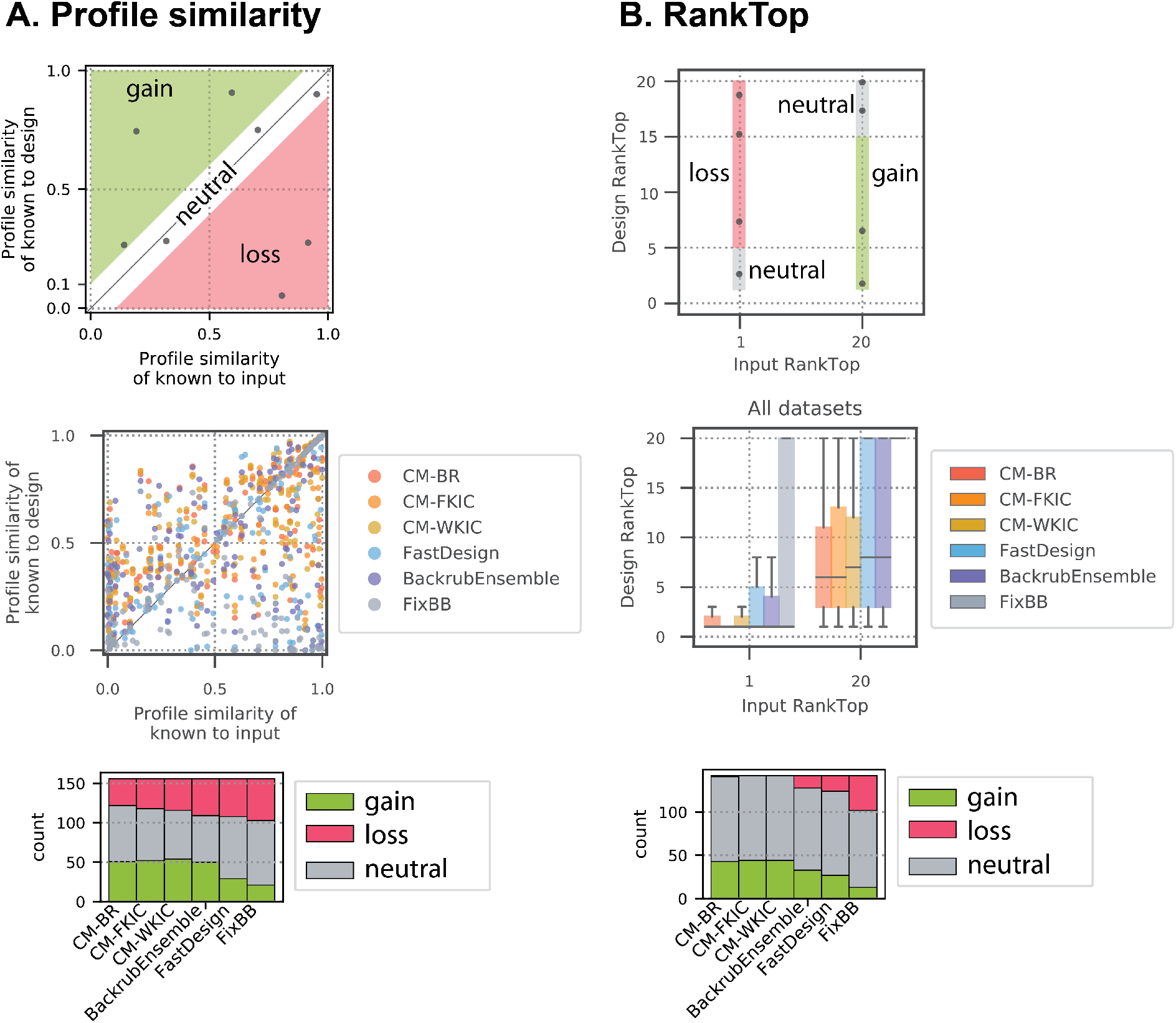
PPS and RankTop as a function of similarity to input. Gain (green) and loss (red) as defined in the main text. Only positions with low and medium entropy (≤ 0.67) are considered. This figure combines all datasets; individual datasets are shown in **Figure S11**. (**A**) PPS as a function of similarity to the input sequence for all profile datasets. Top: Gain and loss zones are defined by a threshold of 0.1 difference between input-known PPS and design-known PPS. Middle: Each point represents one position in the protein sequence, colored by design method. CoupledMoves results (yellow, orange, red) are enriched in the gain zone and FastDesign (blue) and FixBB (grey) results are enriched in the loss zone. Bottom: Quantifications of number of designed sequence positions in gain, loss, and neutral zones for each method. (**B**) RankTop as a function of similarity to the input sequence for all profile datasets, except Fen49, which is omitted because the fentanyl deep sequencing data do not include the input sequence. The top amino acid from the known profile is assigned a rank of 1 if it is present in the input sequence, or a rank of 20 if it is not. Top: A threshold of 5 in the difference in RankTop between input and designed sequences defines the gain and loss zones. Middle: Box plots represent all positions in all datasets, except fentanyl. The median of the distributions is marked with a horizontal line. Second and third quartiles are marked by the box, and the whiskers extend to 1.5 times the inter-quartile range. Bottom: Quantification of sequence positions in gain, loss, and neutral zones for RankTop values.

We also performed a similar analysis for the RankTop values. We defined “loss” as the case where a correct starting amino acid side chain is ranked worse than 5 in the final profile and gain as the case when the known top amino acid side chain is not present in the starting sequence and design models it with a rank of 15 or better (**Figure 6b, top panel**). We only observed loss for positions designed by BackrubEnsemble, FastDesign, and FixBB (**Figure 6b, middle and bottom panels**). CoupledMoves achieves gain with the best median and quartile RankTop values (**Figure 6b, middle panel**), and for the greatest number of positions (**Figure 6b, bottom panel**). Positions are more likely to remain neutral than to experience gain or loss (**Figure 6, bottom panels, Table S11**), thus positions with near-correct input sequence tend to maintain higher PPS values. This observation offers an explanation for the comparatively poor PPS and RankTop values of all methods for the DIG10, Fen49 and hGH/hGHR datasets (**Figure 3**), which are characterized by low similarity between each dataset’s input sequence and known profile (**Figure S11**).

We then asked which methods best predict positions deemed both functionally relevant (consensus) and difficult (requiring gain). We find that CoupledMoves is more likely than non-coupled methods to enrich for correct side chains not present in the input, with 1.2-and 1.5-fold increase in number of positions experiencing gain, compared to BackrubEnsemble and FastDesign, respectively (**Table S11**). In addition, CoupledMoves most consistently avoids loss (0.22- and 0.30-fold decrease in number of positions experiencing loss, compared to BackrubEnsemble and FastDesign, respectively), and retention of correct input side chains (neutral scenario) contributes to overall performance. Taken together, the overall best performance of CoupledMoves arises both from increasing the number of positions with gain and decreasing the number of positions experiencing loss.

We also classified positions as polar/charged or hydrophobic based on the most preferred side chain in the known sequence profile, and use this classification to evaluate performance in recapitulating polar contacts versus hydrophobic packing. CoupledMoves outperforms BackrubEnsemble and FastDesign in discovering and retaining both polar/charged and hydrophobic positions (**Table S11**).

### Selected structural examples

At the Herceptin/HER2 interface, arginine at position VH50 (RVH50) is one of four positions (the other three are YV_H_56, WV_H_95, and YV_H_100a) where CoupledMoves maintains a consensus side chain that is completely lost by one or more non-coupled methods (**Figure S7**). In the crystal structure, RV_H_50 forms a hydrogen bond network across the Herceptin/HER2 interface by interacting with Herceptin TV_L_94 and HER2 E273 and D275. CoupledMoves is able to retain RV_H_50, while FastDesign and BackrubEnsemble replace this residue with hydrophobic residues, predominantly methionine and glycine, respectively (**Figure 7A**).

**Figure 7.**
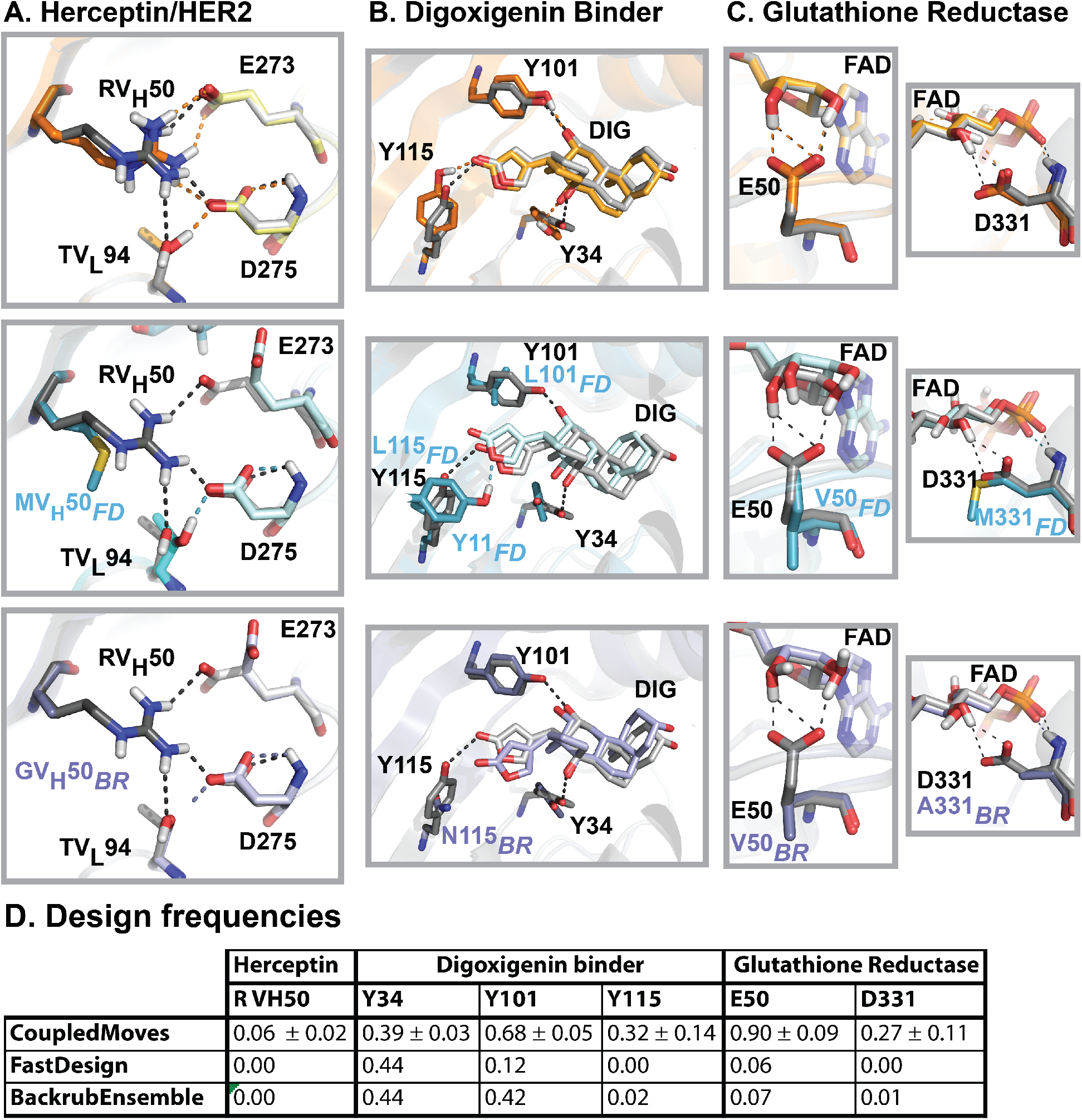
Examples of structural models generated by different design methods. Comparison of crystal structures used as input for design (gray) to models generated by CoupledMoves (top, orange), FastDesign (middle, blue), and BackrubEnsemble (bottom, purple). **(A)** The crystal structure of Herceptin/HER2 (PDB: 1N8Z) shows a hydrogen bond network (black dashed lines) spanning the interface between Herceptin residues RV_H_50 (dark color) and TV_L_94 (medium color), and HER2 residues E273 and D275 (light color). Key designable residue RV_H_50 is retained by CoupledMoves, which models a native-like hydrogen bond network (orange dashed lines). In contrast, FastDesign and BackrubEnsemble model reduced networks (blue and purple dashed lines, respectively). Hydrogen atoms for 1N8Z were added using Rosetta. (**B**) Three tyrosines (Y34, Y101, Y115) form a hydrogen bond network (black dashed lines) with digoxigenin (DIG) in the crystal structure of digoxigenin binder DIG10.2 (PDB: 4J8T). CoupledMoves most frequently retains all three tyrosines and form a similar network (orange dashed lines). FastDesign frequently models leucines at positions 101 and 115, and instead frequently models tyrosine at position 11, forming a hydrogen bond with the ester oxygen rather than carbonyl oxygen of the nearby digoxigenin ring (blue dashed line). BackrubEnsemble most frequently models asparagine at position 115, while retaining the other two contacts (purple dashed lines). (**C**) In crystal structures, glutamate E50 (left column, PDB: 3DK9) and aspartate D331 (right column, PDB: 6FTC) form a hydrogen bond network with flavin-adenine dinucleotide (FAD) (black dashed lines). CoupledMoves retains E50 and D331 in geometries that maintain the network (orange dashed lines). FastDesign and BackrubEnsemble frequently model hydrophobic residues at these positions, abolishing the network. Hydrogen atoms for 3DK9 were added using Rosetta. (**D**) The frequencies of the top known side chain for each position as designed by the different methods. Values for CoupledMoves represent averages and standard deviations across CM-BR, CM-FKIC, and CM-WKIC.

Hydrogen bonds between digoxigenin and the designed protein are most frequently retained by CoupledMoves. In the crystal structure of DIG10.2 (the digoxigenin binder designed with knowledge from the results of the experimental library screen (Tinberg et al., 2013)), tyrosines 34, 101, and 115 hydrogen bond with digoxigenin, as designed (Tinberg et al., 2013). CoupledMoves frequently chooses Tyrosine at all three positions (**Figure 7B, top**), whereas FastDesign often models only one interaction correctly (**Figure 7B, middle**), and BackrubEnsemble models two (**Figure 7B, bottom**). At position 115, BackrubEnsemble most frequently models asparagine, which is too short to hydrogen bond with digoxigenin. FastDesign most frequently models leucine, not tyrosine, at position 115, and instead models Tyrosine at nearby position 11 (alanine consensus in experiment), forming an alternative hydrogen bond with the ester oxygen rather than carbonyl oxygen of the nearby digoxigenin ring.

A third structural example for loss is found in the binding site for cofactor flavin-adenine dinucleotide (FAD) binding site in glutathione reductase (**Figure 7C**). The majority of natural glutathione reductases coordinate FAD with glutamate at position 50 (E50) and aspartate at position 331 (D331). These side chains are frequently maintained by CoupledMoves, but not by FastDesign or BackrubEnsemble (**Figure 7D**). Models generated by CoupledMoves agree with the input crystal structure (3DK9), in which E50 forms a hydrogen bond network with two hydroxyl groups of the 3-4-dihydroxy-furan moiety of FAD. CoupledMoves also predicts a hydrogen bond between evolutionarily conserved residue D331 and a hydroxyl group of FAD. The non-coupled design methods frequently replace both polar side chains with apolar side chains, valine at position E50, and alanine or methionine at D331, eliminating the hydrogen bonds between the protein and the ligand.

### Computational time

We also evaluated the relative compute time for each of the different methods. We first analyzed how performance depended on the number of trajectories run (**Figure S12**). This analysis suggested that performance is optimal for Coupled Moves, BackrubEnsemble and FastDesign at 400, 200 and 100 trajectories, respectively, with slight variation between datasets (**Figure S12**). Since each BackrubEnsemble and FastDesign trajectory takes approximately 2-fold and 20-fold more time than CoupledMoves, respectively, CoupledMoves requires substantially less compute time than FastDesign and about equal compute time to BackrubEnsemble (**Table S12**).

## DISCUSSION

We demonstrate that CoupledMoves recapitulates known sequence profiles at designed positions more accurately than the FastDesign and BackrubEnsemble methods. We consider two conceptual categories of positions: (i) important for function and (ii) difficult to design. For the first category, we classify positions as important for function (in this case binding) either by proxy of low sequence entropy in the known sequence profile, or if specific mutations have been experimentally determined to be important, as in the Enzyme Specificity dataset. CoupledMoves most accurately predicts low entropy consensus positions for all profile benchmarks (**Figure 5**) and outperforms the other methods in correctly identifying specificity-switching mutations in the enzyme specificity set (**Tables 2, 3**). For the second category, we designate positions as difficult to design if the most frequent amino acid side chain in the known profile is not present in the structure used as input for design. Considering both low and medium entropy positions, CoupledMoves is more likely than BackrubEnsemble and FastDesign to correctly identify both charged/polar and hydrophobic side chain residues at higher frequency than in the input sequence (gain), while FastDesign is least likely model a preferred side chain residue present in the input sequence (loss) (**Figure 6**, **Table S11, Figure S11**). We conclude that CoupledMoves is best able to predict both residues that are important for function and difficult to design in our datasets.

**Table 3.** Detailed performance for enzyme specificity benchmark. Percentile and rank are defined as in Table 2.

|  |  |  |  |  |  | Find mutant amino acid starting from<br>wild-type protein structure & mutant ligand |  |  |  |  |  |  |  |  |  |  |  |
| --- | --- | --- | --- | --- | --- | --- | --- | --- | --- | --- | --- | --- | --- | --- | --- | --- | --- |
|  |  |  |  |  |  | CM<br>BR |  | CM<br>FKIC |  | CM<br>WKIC |  | Backrub<br>Ensemble |  | Fast<br>Design |  | FixBB<br>control |  |
| Wild-type<br>PDB ID | Mutant<br>PDB ID | Wild-type<br>Ligand | Mutant<br>Ligand | Desired<br>Mutation | # Design<br>Positions | Percentile | Rank | Percentile | Rank | Percentile | Rank | Percentile | Rank | Percentile | Rank | Percentile | Rank |
| 1A80 | 1M9H | NDP | NAD | K232G | 5 | 79 | 22 | - | 100 | 64 | 37 | - | 100 | - | 100 | - | 100 |
| 1A80 | 1M9H | NDP | NAD | R238H | 5 | 66 | 35 | - | 100 | - | 100 | - | 100 | - | 100 | - | 100 |
| 1FCB | 1SZE | PYR | 173 | L230A | 5 | 98 | 3 | 99 | 2 | 100 | 1 | 97 | 4 | 89 | 12 | 100 | 1 |
| 1K70 | 1RA0 | HPY | FPY | D314A | 4 | 100 | 1 | 98 | 3 | 99 | 2 | 100 | 1 | 98 | 3 | 99 | 2 |
| 1K70 | 1RA5 | HPY | FPY | D314G | 4 | 98 | 3 | 99 | 2 | 98 | 3 | - | 80 | - | 80 | 98 | 3 |
| 1K70 | 1RAK | HPY | FPY | D314S | 4 | - | 80 | - | 80 | - | 80 | - | 80 | 94 | 6 | - | 80 |
| 1PK7 | 1OUM | ADN | TAL | M64V | 3 | 77 | 15 | - | 60 | - | 60 | - | 60 | - | 60 | - | 60 |
| 1ZK4 | 1ZK1 | NAP | NAD | G37D | 7 | 99 | 2 | 98 | 4 | 100 | 1 | 99 | 2 | 100 | 1 | 100 | 1 |
| 2FZN | 3E2Q | PRO | HYP | Y540S | 2 | 95 | 3 | 100 | 1 | 98 | 2 | - | 40 | - | 40 | - | 40 |
| 2H6F | 2H6G | FAR | GER | W602T | 9 | 63 | 68 | 50 | 91 | 66 | 63 | 97 | 6 | 93 | 14 | - | 180 |
| 2O7B | 2O78 | HC4 | TCA | H89F | 4 | 79 | 18 | 70 | 25 | 78 | 19 | - | 80 | - | 80 | - | 80 |
| 3HG5 | 3LX9 | GLA | A2G | E203S | 7 | - | 140 | - | 140 | - | 140 | 92 | 12 | - | 140 | - | 140 |
| 3HG5 | 3LX9 | GLA | A2G | L206A | 7 | 72 | 40 | 93 | 11 | - | 140 | 99 | 2 | 98 | 4 | 100 | 1 |
| 3KZO | 3L02 | AN0 | SN0 | E92A | 5 | 99 | 2 | 99 | 2 | 99 | 2 | - | 100 | 100 | 1 | - | 100 |
| 3KZO | 3L04 | AN0 | SN0 | E92P | 5 | - | 100 | - | 100 | 54 | 47 | - | 100 | - | 100 | - | 100 |
| 3KZO | 3L05 | AN0 | SN0 | E92S | 5 | 94 | 7 | 90 | 11 | 89 | 12 | - | 100 | - | 100 | - | 100 |
| 3KZO | 3L06 | AN0 | SN0 | E92V | 5 | 86 | 15 | 61 | 40 | 82 | 19 | 92 | 9 | - | 100 | - | 100 |

|  |  |  |  |  |  | Find wild-type amino acid starting from<br>mutant protein structure & wild-type ligand |  |  |  |  |  |  |  |  |  |  |  |
| --- | --- | --- | --- | --- | --- | --- | --- | --- | --- | --- | --- | --- | --- | --- | --- | --- | --- |
|  |  |  |  |  |  | CM<br>BR |  | CM<br>FKIC |  | CM<br>WKIC |  | Backrub<br>Ensemble |  | Fast<br>Design |  | FixBB<br>control |  |
| Wild-type<br>PDB ID | Mutant<br>PDB ID | Wild-type<br>Ligand | Mutant<br>Ligand | Desired<br>Mutation | # Design<br>Positions | Percentile | Rank | CMFKIC | Rank | Percentile | Rank | Percentile | Rank | Percentile | Rank | Percentile | Rank |
| 1A80 | 1M9H | NDP | NAD | G232K | 5 | - | 100 | - | 100 | - | 100 | - | 100 | - | 100 | - | 100 |
| 1A80 | 1M9H | NDP | NAD | H238R | 5 | - | 100 | - | 100 | - | 100 | - | 100 | - | 100 | - | 100 |
| 1FCB | 1SZE | PYR | 173 | A230L | 5 | 97 | 5 | 96 | 7 | 96 | 6 | 88 | 18 | - | 100 | - | 100 |
| 1K70 | 1RA0 | HPY | FPY | A314D | 4 | 99 | 2 | 95 | 6 | 98 | 3 | 93 | 8 | 83 | 18 | - | 80 |
| 1K70 | 1RA5 | HPY | FPY | G314D | 4 | 73 | 23 | 74 | 22 | 74 | 22 | - | 80 | - | 80 | - | 80 |
| 1K70 | 1RAK | HPY | FPY | S314D | 4 | - | 80 | - | 80 | - | 80 | - | 80 | - | 80 | - | 80 |
| 1PK7 | 1OUM | ADN | TAL | V64M | 3 | 80 | 13 | 85 | 10 | 82 | 12 | - | 60 | - | 60 | - | 60 |
| 1ZK4 | 1ZK1 | NAP | NAD | D37G | 7 | 100 | 1 | 100 | 1 | 100 | 1 | 99 | 2 | - | 140 | 99 | 2 |
| 2FZN | 3E2Q | PRO | HYP | S540Y | 2 | - | 40 | - | 40 | - | 40 | - | 40 | 88 | 22 | - | 40 |
| 2H6F | 2H6G | FAR | GER | T602W | 9 | 59 | 75 | 69 | 57 | 57 | 79 | 85 | 28 | - | 180 | - | 180 |
| 2O7B | 2O78 | HC4 | TCA | F89H | 4 | 85 | 13 | 74 | 22 | - | 80 | - | 80 | - | 80 | - | 80 |
| 3HG5 | 3LX9 | GLA | A2G | S203E | 7 | 80 | 29 | 76 | 35 | 80 | 29 | 92 | 12 | 91 | 8 | 97 | 5 |
| 3HG5 | 3LX9 | GLA | A2G | A206L | 7 | 59 | 59 | 58 | 60 | 59 | 58 | - | 140 | 88 | 13 | - | 140 |
| 3KZO | 3L02 | AN0 | SN0 | A92E | 5 | 90 | 9 | 94 | 6 | 95 | 5 | 98 | 3 | - | 100 | 100 | 1 |
| 3KZO | 3L04 | AN0 | SN0 | P92E | 5 | 89 | 12 | 88 | 13 | 92 | 9 | 91 | 10 | - | 100 | 95 | 6 |
| 3KZO | 3L05 | AN0 | SN0 | S92E | 5 | 84 | 17 | 85 | 16 | 86 | 15 | 95 | 6 | - | 100 | - | 100 |
| 3KZO | 3L06 | AN0 | SN0 | V92E | 5 | 86 | 18 | 93 | 10 | 89 | 14 | 74 | 32 | - | 100 | 95 | 7 |

To provide insights into why the different methods model consensus side chains with different frequencies, despite using the same energy function, we analyzed how the correct amino acid at these positions was ranked by energy for each of the different methods. **Figure 8a** shows distributions of percentiles for predicted total Rosetta energy of instances where a method models the known top ranked amino acid side chain. These distributions are shifted towards higher percentiles for CoupledMoves compared to the other methods. CM-FKIC predicts the consensus side chain for 51 positions with total energy above the 75^th^ percentile, while BackrubEnsemble and FastDesign predict 37 and 27 positions in the same category. CoupledMoves models the consensus side chain for a total of 132 designable positions in the datasets, compared to 111 and 95 positions for BackrubEnsemble and FastDesign, respectively. The high sequence entropy of CoupledMoves design compared to other methods (**Figure S10**) makes it even more remarkable that CoupledMoves ranks the energetics of consensus side chains so favorably among many options. We conclude that, for side chains modeled with > 0.33 frequency and > 75^th^ energy percentile, CoupledMoves predictions are likely correct.

**Figure 8.**
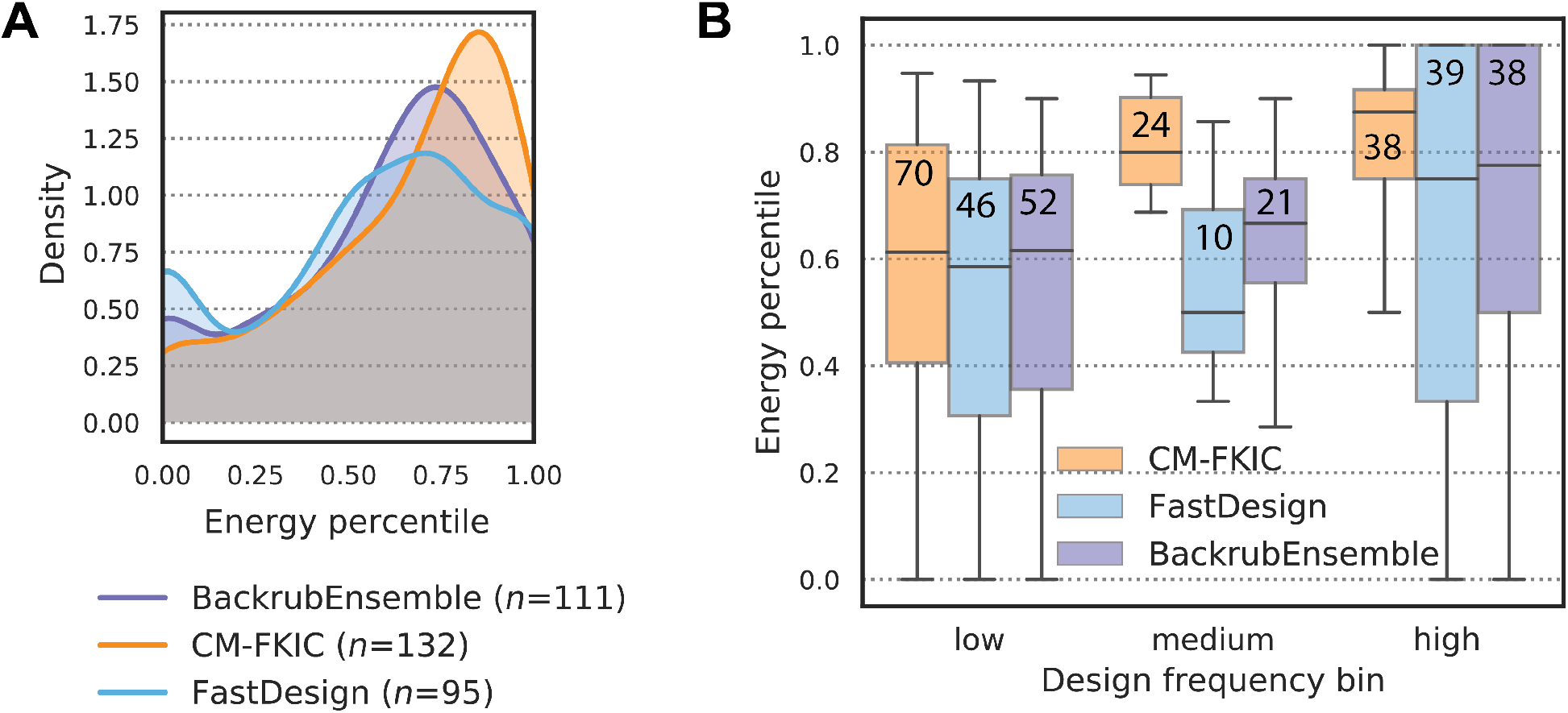
Distribution of energy percentiles for correctly modeled positions. “Energy percentile” refers to the percentile of the average total Rosetta energy of the correctly modeled side chain compared to that of all other side chains modeled by the design method at that position. Energy percentile was calculated for consensus (entropy ≤ 0.67) positions for which a method modeled the consensus at least once. (**A**) Distribution of energy percentiles. Count *n* indicates the number of positions for which each method modeled the consensus side chain at least once. (**B**) Energy percentiles as a function of design frequency are shown as boxplots. Values from (**A**) are binned by design frequency (low: frequency ≤ 0.33, medium: 0.33 < frequency ≤ 0.67, and high: frequency > 0.67). The number of values in each bin is shown on each boxplot. The median of the distributions is marked with a horizontal line. Second and third quartiles are marked by the box, and the whiskers extend to 1.5 times the inter-quartile range.

In cases where the BackrubEnsemble method does model the consensus side chain during design, the energetics rank favorably (**Figure 8a**). One possible reason for the overall worse performance of BackrubEnsemble over CoupledMoves is that cases correctly predicted by BackrubEnsemble might be derived from only a subset of ensemble members whose backbone conformations are compatible with energetically favorable placement of the consensus side chain. In these cases, the input/consensus side chain is compatible with the ensemble, but during sequence design another amino acid side chain has more favorable Rosetta energy. Indeed, forcing the consensus side chains onto all ensemble members results in a greater proportion of models with unfavorable (positive) Rosetta energy, and a smaller proportion of models with highly favorable energy (**Figure 9**, shown are glutathione reductase and digoxigenin binder, which are examples of loss by the BackrubEnsemble method). This behavior suggests that ensemble members are not uniformly compatible with consensus sidechains, highlighting a limitation of the BackrubEnsemble method. Backbone moves are sampled only once, at the beginning of the trajectory during ensemble creation (**Figure 1a**). Sidechains are subsequently modeled onto each ensemble member by finding an energetically favorable rotamer for the pre-determined backbone conformation. In contrast, the CoupledMoves design trajectory cycles small backbone adjustments in response to sequence change moves, which allows switching from non-consensus to consensus side chains. Without cycles of backbone and sidechain sampling, the BackrubEnsemble method is limited to snapshots of the allowed backbone conformational diversity defined by the initial ensemble members.

**Figure 9.**
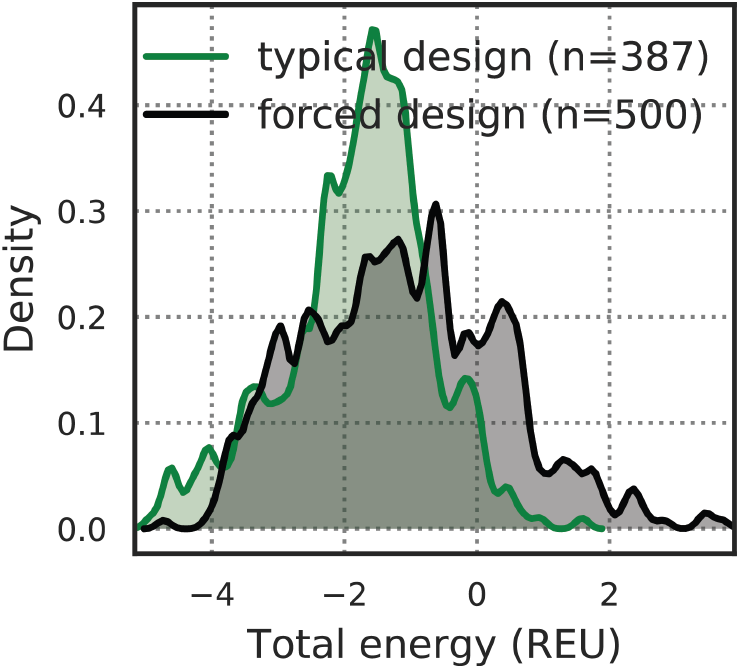
Comparison between typical and forced design of consensus side chains onto Backrub ensemble. Distribution of Rosetta energies (REU) for consensus amino acid side chains at five positions: Glutathione Reductase positions E50 and D331, and DIG10 positions Y34, Y101, and Y115. For each position, we show 100 models forced to adopt the consensus side chain during sequence design (black), and typical models (green) that arrived at the consensus side chain though they were allowed to design to multiple side chain identities. For typical models, n corresponds to the number of models with the known consensus side chain, out of a total of 2000 (400 models for each of the five positions; design frequencies are shown in Figure 7d). Density represents a Gaussian kernel density estimate using a bin width of 0.1 REU.

For CoupledMoves, the design frequency increases with energy percentile for consensus side chains (**Figure 8b**), which is expected - side chains with a higher (more favorable) energy percentile should be chosen more frequently. However, this trend is less pronounced for both BackrubEnsemble and FastDesign. For BackrubEnsemble, this behavior is possibly due to the limitations enforced by the backbones conformations of the ensemble. In the case of FastDesign, it is possible that the minimization step in FastDesign is prone to trapping the design simulations in local minima and hence that the frequency of chosen amino acids poorly reflect their actual fitness rank. This hypothesis is supported by the low entropy of FastDesign design sequence profiles (**Figure S10**). FastDesign may be less likely to escape local minima than the other methods, despite the use of a reduced Lennard-Jones repulsive term in the early cycles of the simulation (**Figure 1a**).

In addition to limitations in sampling methods (as well as the energy function used to rank designs), there are also potential limitations inherent to the benchmark datasets. For example, in the case of the enzyme specificity dataset, we can only compare to the point mutations that were experimentally tested, but we do not have sequence profiles. The enzymes have not been subject to saturation mutagenesis, so it is unknown whether additional specificity-altering mutations could exist.

Sequence profiles in the cofactor dataset result from natural evolution, rather than experimental screening. Natural evolution includes selection pressures beyond affinity (catalysis, specificity, kinetics etc.), so that the sequence profiles for natural binding site positions may be influenced by factors beyond those modeled here. In addition, our analysis does not evaluate covariation between residue positions. However, evolutionary sequence profiles have the advantage of clearly identifying consensus binding positions, and we observe considerable agreement between Rosetta predicted and known sequence profiles for this set.

Finally, all methods tested perform most poorly at consensus positions in the deep sequencing datasets, DIG10 and Fen49, and the design methods perform worse than the null model on DIG10. Initiating design from the crystal structure corresponding to the result of the library selection (PDB: 4J8T) did not improve performance. It is possible that the selection experiments report on additional considerations such as expression and display on the yeast surface that are not considered in the design simulations, or that the sensitivity range of the selection is tuned to primarily differentiate between functional versus deleterious mutations but is less capable of quantitatively ranking binding affinity. Alternatively, critical adjustments of both backbones and the ligand, in addition to ligand strain and ligand flexibility, are not correctly captured in the Rosetta simulations.

Apart from suggesting individual point mutations such as in the enzyme specificity set, our results on recapitulating position-specific sequence profiles highlight the utility of CoupledMoves for generating libraries. CoupledMoves will be most useful in design cases where protein backbones are supplied with existing side chains, such as natural or previously-characterized designed proteins (rather than the *de novo* design of new structures). Computation has long been used to reduce the sequence space queried by library screens (Hayes et al., 2002; Treynor et al., 2007; Voigt et al., 2001), and it is well established that flexible-backbone protein design can generate sequences similar to observed natural and experimental sequences (Ding & Dokholyan, 2006; Friedland et al., 2009; Humphris & Kortemme, 2008; Humphris-Narayanan et al., 2012; Koehl & Levitt, 2002; Larson et al., 2003; Saunders & Baker, 2005; Smith & Kortemme, 2011). As the design results obtained with CoupledMoves most accurately reflect tolerated sequence space in comparison to other methods using the same energy function, CoupledMoves represents a powerful flexible backbone strategy for generating combinatorial libraries for screening and selection, and optimizing proteins for new and useful functions.

Computational design of binding sites remains difficult, in part due to limitations in current ability to realistically sample backbone conformations that enable side chains to make realistic contacts during sequence design. The benchmarking framework presented here can be adapted to different types of design applications, such as sequence design on parametrically-generated rather than natural protein backbones, or *de novo* designed rather than reengineered binding sites.

## Supporting information

Supplementary Infomation

## Acknowledgments

This work was supported by grants from the National Institutes of Health (NIH R01-GM110089) and the National Science Foundation (NSF DBI-1564692) to TK. The authors would like to thank Kale Kundert and Noah Ollikainen for helpful discussions, and Christy Tinberg and Matt Bick for sharing data.

## REFERENCES

Alford, R. F., Leaver-Fay, A., Jeliazkov, J. R., et al. (2017). The Rosetta All-Atom Energy Function for Macromolecular Modeling and Design. J Chem Theory Comput, 13(6), 3031–3048.

Babor, M., Mandell, D. J., & Kortemme, T. (2011). Assessment of flexible backbone protein design methods for sequence library prediction in the therapeutic antibody Herceptin-HER2 interface. Protein Sci, 20(6), 1082–1089.

Baldwin, E. P., Hajiseyedjavadi, O., Baase, W. A., et al. (1993). The role of backbone flexibility in the accommodation of variants that repack the core of T4 lysozyme. Science, 262(5140), 1715–1718.

Bick, M. J., Greisen, P. J., Morey, K. J., et al. (2017). Computational design of environmental sensors for the potent opioid fentanyl. Elife, 6.

Chevalier, A., Silva, D. A., Rocklin, G. J., et al. (2017). Massively parallel de novo protein design for targeted therapeutics. Nature, 550(7674), 74–79.

Correia, B. E., Bates, J. T., Loomis, R. J., et al. (2014). Proof of principle for epitope-focused vaccine design. Nature, 507(7491), 201–206.

Coutsias, E. A., Seok, C., Jacobson, M. P., et al. (2004). A kinematic view of loop closure. J Comput Chem, 25(4), 510–528.

Cunningham, B. C., & Wells, J. A. (1993). Comparison of a structural and a functional epitope. J Mol Biol, 234(3), 554–563.

Dahiyat, B. I., & Mayo, S. L. (1997). De novo protein design: fully automated sequence selection. Science, 278(5335), 82–87.

Davey, J. A., & Chica, R. A. (2014). Improving the accuracy of protein stability predictions with multistate design using a variety of backbone ensembles. Proteins, 82(5), 771–784.

Davey, J. A., & Chica, R. A. (2017). Multistate Computational Protein Design with Backbone Ensembles. Methods Mol Biol, 1529, 161–179.

Davis, I. W., Arendall, W. B., 3rd, Richardson, D. C., et al. (2006). The backrub motion: how protein backbone shrugs when a sidechain dances. Structure, 14(2), 265–274.

Desjarlais, J. R., & Handel, T. M. (1999). Side-chain and backbone flexibility in protein core design. J Mol Biol, 290(1), 305–318.

Ding, F., & Dokholyan, N. V. (2006). Emergence of protein fold families through rational design. PLoS Comput Biol, 2(7), e85.

Dou, J., Doyle, L., Jr Greisen, P., et al. (2017). Sampling and energy evaluation challenges in ligand binding protein design. Protein Sci, 26(12), 2426–2437.

Dou, J., Vorobieva, A. A., Sheffler, W., et al. (2018). De novo design of a fluorescence-activating beta-barrel. Nature, 561(7724), 485–491.

Fleishman, S. J., Whitehead, T. A., Ekiert, D. C., et al. (2011). Computational design of proteins targeting the conserved stem region of influenza hemagglutinin. Science, 332(6031), 816–821.

Friedland, G. D., Lakomek, N. A., Griesinger, C., et al. (2009). A correspondence between solution-state dynamics of an individual protein and the sequence and conformational diversity of its family. PLoS Comput Biol, 5(5), e1000393.

Friedland, G. D., Linares, A. J., Smith, C. A., et al. (2008). A simple model of backbone flexibility improves modeling of side-chain conformational variability. J Mol Biol, 380(4), 757–774.

Fu, X., Apgar, J. R., & Keating, A. E. (2007). Modeling backbone flexibility to achieve sequence diversity: the design of novel alpha-helical ligands for Bcl-xL. J Mol Biol, 371(4), 1099–1117.

Georgiev, I., Keedy, D., Richardson, J. S., et al. (2008). Algorithm for backrub motions in protein design. Bioinformatics, 24(13), i196–204.

Gerstner, R. B., Carter, P., & Lowman, H. B. (2002). Sequence plasticity in the antigen-binding site of a therapeutic anti-HER2 antibody. J Mol Biol, 321(5), 851–862.

Gordon, D. B., Marshall, S. A., & Mayo, S. L. (1999). Energy functions for protein design. Curr Opin Struct Biol, 9(4), 509–513.

Hayes, R. J., Bentzien, J., Ary, M. L., et al. (2002). Combining computational and experimental screening for rapid optimization of protein properties. Proc Natl Acad Sci U S A, 99(25), 15926–15931.

Humphris, E. L., & Kortemme, T. (2008). Prediction of protein-protein interface sequence diversity using flexible backbone computational protein design. Structure, 16(12), 1777–1788.

Humphris-Narayanan, E., Akiva, E., Varela, R., et al. (2012). Prediction of mutational tolerance in HIV-1 protease and reverse transcriptase using flexible backbone protein design. PLoS Comput Biol, 8(8), e1002639.

Jiang, L., Althoff, E. A., Clemente, F. R., et al. (2008). De novo computational design of retro-aldol enzymes. Science, 319(5868), 1387–1391.

Kapp, G. T., Liu, S., Stein, A., et al. (2012). Control of protein signaling using a computationally designed GTPase/GEF orthogonal pair. Proc Natl Acad Sci U S A, 109(14), 5277–5282.

Keedy, D. A., Georgiev, I., Triplett, E. B., et al. (2012). The role of local backrub motions in evolved and designed mutations. PLoS Comput Biol, 8(8), e1002629.

Khatib, F., Cooper, S., Tyka, M. D., et al. (2011). Algorithm discovery by protein folding game players. Proc Natl Acad Sci U S A, 108(47), 18949–18953.

Koehl, P., & Levitt, M. (2002). Protein topology and stability define the space of allowed sequences. Proc Natl Acad Sci U S A, 99(3), 1280–1285.

Koga, N., Tatsumi-Koga, R., Liu, G., et al. (2012). Principles for designing ideal protein structures. Nature, 491(7423), 222–227.

Kuhlman, B., Dantas, G., Ireton, G. C., et al. (2003). Design of a novel globular protein fold with atomic-level accuracy. Science, 302(5649), 1364–1368.

Kundert, K., & Kortemme, T. (2019). Computational design of structured loops for new protein functions. Biol Chem, 400(3), 275–288.

Larson, S. M., England, J. L., Desjarlais, J. R., et al. (2002). Thoroughly sampling sequence space: large-scale protein design of structural ensembles. Protein Sci, 11(12), 2804–2813.

Larson, S. M., Garg, A., Desjarlais, J. R., et al. (2003). Increased detection of structural templates using alignments of designed sequences. Proteins, 51(3), 390–396.

Leaver-Fay, A., Tyka, M., Lewis, S. M., et al. (2011). ROSETTA3: an object-oriented software suite for the simulation and design of macromolecules. Methods Enzymol, 487, 545–574.

Mandell, D. J., Coutsias, E. A., & Kortemme, T. (2009). Sub-angstrom accuracy in protein loop reconstruction by robotics-inspired conformational sampling. Nat Methods, 6(8), 551–552.

Martin, A. C. (1996). Accessing the Kabat antibody sequence database by computer. Proteins, 25(1), 130–133.

Ollikainen, N., de Jong, R. M., & Kortemme, T. (2015). Coupling Protein Side-Chain and Backbone Flexibility Improves the Re-design of Protein-Ligand Specificity. PLoS Comput Biol, 11(9), e1004335.

Ollikainen, N., & Kortemme, T. (2013). Computational protein design quantifies structural constraints on amino acid covariation. PLoS Comput Biol, 9(11), e1003313.

Ollikainen, N., Smith, C. A., Fraser, J. S., et al. (2013). Flexible backbone sampling methods to model and design protein alternative conformations. Methods Enzymol, 523, 61–85.

Pal, G., Kouadio, J. L., Artis, D. R., et al. (2006). Comprehensive and quantitative mapping of energy landscapes for protein-protein interactions by rapid combinatorial scanning. J Biol Chem, 281(31), 22378–22385.

Park, H., Bradley, P., Greisen, P., Jr., et al. (2016). Simultaneous Optimization of Biomolecular Energy Functions on Features from Small Molecules and Macromolecules. J Chem Theory Comput, 12(12), 6201–6212.

Ponder, J. W., & Richards, F. M. (1987). Tertiary templates for proteins. Use of packing criteria in the enumeration of allowed sequences for different structural classes. J Mol Biol, 193(4), 775–791.

Procko, E., Berguig, G. Y., Shen, B. W., et al. (2014). A computationally designed inhibitor of an Epstein-Barr viral Bcl-2 protein induces apoptosis in infected cells. Cell, 157(7), 1644–1656.

Rocklin, G. J., Chidyausiku, T. M., Goreshnik, I., et al. (2017). Global analysis of protein folding using massively parallel design, synthesis, and testing. Science, 357(6347), 168–175.

Rothlisberger, D., Khersonsky, O., Wollacott, A. M., et al. (2008). Kemp elimination catalysts by computational enzyme design. Nature, 453(7192), 190–195.

Saunders, C. T., & Baker, D. (2005). Recapitulation of protein family divergence using flexible backbone protein design. J Mol Biol, 346(2), 631–644.

Silva, D. A., Yu, S., Ulge, U. Y., et al. (2019). De novo design of potent and selective mimics of IL-2 and IL-15. Nature, 565(7738), 186–191.

Simons, K. T., Bonneau, R., Ruczinski, I., et al. (1999). Ab initio protein structure prediction of CASP III targets using ROSETTA. Proteins, Suppl 3, 171–176.

Simons, K. T., Kooperberg, C., Huang, E., et al. (1997). Assembly of protein tertiary structures from fragments with similar local sequences using simulated annealing and Bayesian scoring functions. J Mol Biol, 268(1), 209–225.

Smith, C. A., & Kortemme, T. (2008). Backrub-like backbone simulation recapitulates natural protein conformational variability and improves mutant side-chain prediction. J Mol Biol, 380(4), 742–756.

Smith, C. A., & Kortemme, T. (2010). Structure-based prediction of the peptide sequence space recognized by natural and synthetic PDZ domains. J Mol Biol, 402(2), 460–474.

Smith, C. A., & Kortemme, T. (2011). Predicting the tolerated sequences for proteins and protein interfaces using RosettaBackrub flexible backbone design. PLoS One, 6(7), e20451.

Tinberg, C. E., Khare, S. D., Dou, J., et al. (2013). Computational design of ligand-binding proteins with high affinity and selectivity. Nature, 501(7466), 212–216.

Treynor, T. P., Vizcarra, C. L., Nedelcu, D., et al. (2007). Computationally designed libraries of fluorescent proteins evaluated by preservation and diversity of function. Proc Natl Acad Sci U S A, 104(1), 48–53.

Tyka, M. D., Keedy, D. A., Andre, I., et al. (2011). Alternate states of proteins revealed by detailed energy landscape mapping. J Mol Biol, 405(2), 607–618.

Voigt, C. A., Mayo, S. L., Arnold, F. H., et al. (2001). Computational method to reduce the search space for directed protein evolution. Proc Natl Acad Sci U S A, 98(7), 3778–3783.

