## Supplementary Infomation for "Comparison of Rosetta flexible-backbone computational protein design methods on binding interactions"

### SUPPORTING INFORMATION

### SUPPLEMENTAL TABLES

**Table S1. Cofactor dataset ligands.**
Ligand name and 3-letter PDB ligand identifier are shown for each protein family of the Cofactor dataset.

| **Protein** | **Ligand** |
|---|---|
| Acetyl Transferase | Coenzyme A (COA) |
| Alcohol Dehydrogenase | nicotinamide-adenine-dinucleotide phosphate (NADP) (NAP) |
| Amino-transferase | 4'-deoxy-4'-aminopyridoxal-5'-phosphate (PMP) |
| Flavodoxin | flavin mono-nucleotide (FMN) |
| Glutathione S-Transferase | glutathione (GSH) |
| Methyl-transferase | S-adenosyl-methionine (SAM) |
| Glutathione Reductase | flavin-adenine dinucleotide (FAD) |

**Table S2. Cofactor benchmark structures and positions.**
Protein names and PDB codes are shown, along with designable and packable positions. Position numbering corresponds to PDB numbering.

| Protein | PDB | Designable positions | Packable positions |
|---|---|---|---|
| Acetyl Transferase | 3S6F | 78, 79, 80, 85, 86, 87, 88, 90, 91, 114, 115, 118, 119, 121 (n = 14) | 4, 6, 26, 27, 28, 52, 60, 62, 63, 75, 77, 81, 84, 89, 93, 94, 95, 109, 111, 112, 116, 117, 122, 124, 126, ligand (n = 26) |
| Alcohol Dehydrog-enase | 1ZK4 | 13, 14, 15, 16, 17, 18, 19, 36, 37, 38, 62, 63, 89, 90, 91, 92, 112, 140, 142, 155, 159, 187, 189, 190, 192, 194, 195, 205 (n = 28) | 11, 12, 22, 23, 34, 35, 39, 42, 45, 46, 58, 59, 61, 64, 69, 72, 87, 93, 94, 108, 109, 113, 117, 138, 143, 144, 149, 152, 156, 158, 162, 185, 186, 191, 197, 198, 201, 202, 206, 210, 211, 216, 219, 221, 222, 244, 248, ligand (n = 48) |
| Amino-transferase | 2XBN | 73, 133, 134, 135, 138, 159, 161, 202, 231, 233, 234, 262, 264, 265 (n = 14) | 75, 136, 137, 139, 141, 142, 155, 158, 162, 164, 200, 204, 205, 206, 236, 260, 270, 273, 347, 349, 390, 392, ligand (n = 23) |
| Flavodoxin | 1F4P | 10, 11, 12, 13, 14, 15, 58, 59, 60, 61, 62, 68, 93, 94, 95, 98, 100, 101, 102 (n = 19) | 8, 16, 17, 18, 19, 57, 65, 66, 69, 70, 71, 91, 96, 97, 105, 106, 125, 126, 127, 130, ligand (n = 21) |
| Glutathione S-Transferase | 3R2Q | 9, 10, 11, 33, 34, 48, 49, 50, 62, 63, 64, 98, 105 (n = 13) | 3, 4, 6, 12, 13, 14, 15, 31, 38, 40, 43, 44, 46, 51, 52, 61, 66, 67, 68, 91, 94, 95, 101, 102, 108, 109, 112, 157, 160, 164, 168, ligand (n = 32) |
| Methyl-transferase | 3DLC | 8, 16, 20, 28, 48, 49, 50, 51, 52, 53, 55, 72, 73, 74, 77, 100, 101, 102, 117, 118, 119, 122, 123 (n = 23) | 7, 13, 17, 19, 21, 24, 27, 31, 32, 46, 47, 56, 57, 58, 59, 68, 70, 71, 76, 78, 80, 81, 84, 103, 104, 114, 115, 116, 120, 121, 125, 126, 128, 129, 130, 132, 133, 144, 207, ligand (n = 40) |
| Glutathione Reductase | 3DK9 | 26, 27, 28, 29, 30, 31, 49, 50, 51, 52, 56, 57, 58, 62, 63, 66, 129, 130, 155, 156, 157, 177, 181, 197, 198, 201, 202, 291, 294, 298, 330, 331, 337, 338, 339, 340, 342, 372 (n = 38) | 24, 25, 33, 35, 47, 48, 54, 61, 64, 65, 67, 70, 103, 114, 125, 126, 127, 131, 132, 140, 142, 147, 153, 154, 159, 160, 180, 192, 200, 205, 206, 223, 226, 286, 288, 295, 297, 300, 329, 332, 336, 341, 343, 344, 369, 370, 371, 373, 376, 377, 441, ligand (n = 52) |

**Table S3. Enzyme dataset.**
Ligand names are shown for wild-type and mutant proteins of the Enzyme specificity dataset.

| **Protein** | **Wild-type ligand** | **Mutant ligand** |
|---|---|---|
| 2-5-diketo-D-gluconic acid reductase A | dihydro-nicotinamide-adenine-dinucleotide phosphate (NADPH) (NDP) | nicotinamide-adenine-dinucleotide (NAD) |
| Alcohol dehydrogenase | nicotinamide-adenine-dinucleotide (NAD) | NADP nicotinamide-adenine-dinucleotide phosphate (NAP) |
| Alpha-galactosidase A | N-actyl-2-deoxy-2-amino-galactose (A2G) | alpha D-galactose (GLA) |
| Cytosine deaminase | (4S)-5-fluoro-4-hydroxy-3,4-dihydropyrimidin-2(1H)-one (FPY) | 4-hydroxy-3,4-dihydro-1H-pyrimidin-2-one (HPY) |
| Farnesyltransferase | geran-8-yl geran (GER) | farnesyl (FAR) |
| Flavocytochrome b(2) | benzoyl-formic ACID (173) | pyruvic acid (PYR) |
| Histidine ammonialyase | phenylethylene-carboxylic acid (TCA) | para-coumaric acid (HC4) |
| N-acetylornithine carbamoyltransferase | N-(3-carboxypropanoyl-L-norvaline (SN0) | N-acetyl-L-norvaline (AN0) |
| Proline dehydrogenase | 4-hydroxyproline (HYP) | proline (PRO) |
| Purine nucleoside phosphorylase | 9-(6-deoxy-alpha-L-talofuranosyl)-6-methylpurine (TAL) | adenosine (ADN) |

**Table S4. Enzyme specificity benchmark structures and positions.**
Proteins (wild-type and mutant) and ligand names and PDB codes are shown, along with designable and packable positions. Position numbering corresponds to PDB numbering. **Continued on next page.**

| Protein | PDB | Designable positions | Packable positions |
|---|---|---|---|
| 2-5-diketo-D-gluconic acid reductase A | 1M9H (mutant) | 232, 233, 234, 235, 238 (n = 5) | 19, 22, 23, 24, 25, 28, 32, 41, 190, 215, 231, 237, 239, 241, 242, ligand (n = 16) |
| | 1A80 (wild-type) | 232, 233, 234, 235, 238 (n = 5) | 19, 22, 23, 24, 25, 28, 32, 41, 190, 215, 231, 237, 239, 241, 242, ligand (n = 16) |
| Alcohol dehydrogenase | 1ZK1 (mutant) | 13, 14, 15, 16, 36, 37, 38, 42 (n = 8) | 10, 11, 12, 23, 33, 34, 35, 39, 41, 46, 58, 59, 61, 62, 63, 64, 69, 72, 89, 90, 112, 192, 193, 194, ligand (n = 25) |
| | 1ZK4 (wild-type) | 13, 14, 15, 16, 36, 37, 38 (n = 7) | 10, 11, 12, 23, 33, 34, 35, 39, 42, 46, 56, 58, 59, 61, 62, 63, 69, 72, 89, 90, 192, 193, 194, ligand (n = 24) |
| Alpha-galactosidase A | 3LX9 (mutant) | 170, 203, 206, 207, 227, 229, 231 (n = 7) | 47, 92, 93, 134, 136, 137, 141, 142, 168, 172, 174, 177, 180, 184, 201, 204, 208, 209, 228, 241, 242, 245, 246, 249, 253, 264, 266, 267, ligand (n = 29) |
| | 3HG5 (wild-type) | 170, 203, 206, 207, 227, 229, 231 (n = 7) | 47, 51, 92, 93, 134, 136, 137, 141, 142, 168, 172, 174, 177, 180, 184, 201, 204, 208, 209, 211, 228, 241, 242, 245, 246, 249, 264, 265, 266, 267, ligand (n = 31) |
| Cytosine deaminase | 1K70 (wild-type) | 63, 313, 314, 319 (n = 4) | 61, 65, 66, 81, 85, 88, 122, 124, 154, 214, 217, 246, 273, 275, 278, 279, 282, 317, 318, 320, ligand (n = 21) |
| | 1RA5 (mutant) | 63, 313, 314, 319 (n = 4) | 61, 65, 66, 81, 85, 88, 122, 124, 154, 156, 214, 217, 246, 273, 275, 278, 279, 282, 317, 318, 320, ligand (n = 22) |
| | 1RAK (mutant) | 63, 313, 314, 319 (n = 4) | 61, 65, 66, 81, 85, 88, 122, 124, 154, 156, 214, 217, 246, 273, 275, 278, 279, 282, 317, 318, 320, ligand (n = 22) |
| | 1RA0 (mutant) | 63, 313, 314, 317, 319 (n = 5) | 61, 65, 66, 69, 81, 85, 88, 122, 124, 154, 214, 217, 246, 273, 275, 278, 279, 282, 318, 320, ligand (n = 21) |
| Farnesyltransferase | 2H6G (mutant) | 602, 605, 606, 651, 654, 655, 706, 803, 865 (n = 9) | 596, 599, 603, 609, 649, 650, 652, 658, 662, 693, 702, 703, 705, 709, 710, 748, 753, 799, 800, 802, 860, 861, 862, 864, 868, 902, 903, ligand (n = 28) |
| | 2H6F (wild-type) | 602, 605, 606, 654, 655, 705, 706, 803, 865 (n = 9) | 596, 599, 603, 609, 650, 651, 658, 662, 693, 702, 703, 709, 748, 753, 754, 757, 761, 799, 800, 802, 860, 861, 862, 864, 868, 902, 903, ligand (n = 28) |

**Table S4, continued: Enzyme specificity benchmark structures and positions.**
Proteins (wild-type and mutant) and ligand names and PDB codes are shown, along with designable and packable positions. Position numbering corresponds to PDB numbering.

| Protein | PDB | Designable positions | Packable positions |
|---|---|---|---|
| Flavocytochrome b(2) | 1SZE (mutant) | 143, 198, 230, 254, 286, 325, 326 (n = 7) | 139, 144, 199, 202, 228, 229, 252, 256, 280, 283, 289, 292, 296, 323, 324, 373, 377, ligand (n = 18) |
| | 1FCB (wild-type) | 143, 198, 230, 254, 326 (n = 5) | 139, 144, 199, 202, 228, 229, 252, 280, 283, 289, 292, 296, 323, 325, 373, 376, ligand (n = 17) |
| Histidine ammonia-lyase | 2O78 (mutant) | 89, 90, 405, 406 (n = 4) | 66, 68, 69, 86, 87, 153, 154, 157, 202, 432, 381, 391, 392, 402, 408, 409, 503, ligand (n = 18) |
| | 2O7B (wild-type) | 89, 90, 405, 406 (n = 4) | 66, 68, 69, 86, 87, 153, 154, 157, 202, 432, 391, 392, 402, 408, 409, 503, ligand (n = 17) |
| N-acetylornithine carbamoyltransferase | 3L05 (mutant) | 180, 296, 298, 77, 92 (n = 5) | 48, 50, 51, 181, 182, 184, 252, 253, 270, 293, 301, 302, 78, 93, 98, ligand (n = 16) |
| | 3L06 (mutant) | 180, 184, 298, 302, 77, 92 (n = 6) | 48, 50, 51, 112, 178, 181, 182, 251, 252, 253, 270, 291, 293, 296, 297, 301, 303, 308, 78, 93, ligand (n = 21) |
| | 3L04 (mutant) | 180, 184, 298, 77, 92 (n = 5) | 48, 50, 51, 181, 182, 252, 253, 270, 296, 301, 302, 78, 93, ligand (n = 14) |
| | 3L02 (mutant) | 180, 298, 77, 92 (n = 4) | 48, 50, 51, 181, 182, 184, 252, 253, 270, 293, 296, 301, 302, 78, 93, 98, ligand (n = 17) |
| | 3KZO (wild-type) | 180, 184, 298, 77, 92 (n = 5) | 48, 50, 51, 181, 182, 252, 253, 270, 293, 296, 302, 78, 93, 98, ligand (n = 15) |
| Proline dehydrogenase | 2FZN (wild-type) | 513, 540 (n = 2) | 259, 283, 285, 327, 370, 431, 485, 487, 511, 516, 538, 542, 552, 556, 559, 560, ligand (n = 17) |
| | 3E2Q (mutant) | 285, 513, 540 (n = 3) | 259, 283, 287, 327, 329, 370, 431, 485, 487, 511, 516, 542, 552, 556, 559, 560, ligand (n = 17) |
| Purine nucleoside phosphorylase | 1OUM (mutant) | 64, 180, 181 (n = 3) | 62, 69, 73, 87, 159, 179, 185, 198, ligand (n = 9) |
| | 1PK7 (wild-type) | 64, 159, 180 (n = 3) | 62, 156, 160, 181, ligand (n = 5) |

**Table S5. Digoxigenin benchmark structures and positions.**
PDB codes are shown for the source of the protein and ligand structure. Position and starting side chain identity are shown for designable and packable positions. Table lists positions packed by CoupledMoves methods; other methods pack all positions. Positions are numbered as in (Tinberg et al., 2013).

| **Protein** | **PDB** | **Designable positions** | **Packable positions (coupled methods)** |
|---|---|---|---|
| Designed digoxigenin binder DIG10.1 | 1Z1S (protein), 4J8T (ligand) | A10, L11, L14, W22, C23, F26, L32, Y34, A37, P38, G40, H41, F45, H54, M55, F58, Y61, M62, I64, F66, F84, G86, G88, H90, V92, S93, G95, L97, A99, Y101, S103, L105, I112, Y115, L117, F119, V124, P127, L128 (n = 39) | I6, L7, V8, H9, R12, L13, E15, A19, R20, L25, P39, K42, T43, R48, E49, T50, I51, W52, L57, P59, E60, V69, F71, A80, T91, T107, P121, R123, L125, I6, L7, V8, H9, R12, L13, E15, A19, R20, L25, P39, K42, T43, R48, E49, T50, I51, W52, L57, P59, E60, V69, F71, A80, T91, T107, P121, R123, L125, DIG (n = 30) |

**Table S6. Allowed design for digoxigenin dataset.**

Shown are amino acids (one letter codes) to which positions were allowed to design. Amino acids were included only if they had high enough sequencing counts to be included in the enrichment and depletion calculations in (Tinberg et al., 2013).

| Position | Allowed amino acids |
|---|---|
| 10 | ACDEFGILMNPRSTVY |
| 11 | ACDFGHILMNPQRSTVY |
| 14 | AFHIKLMPQRSTVW |
| 22 | ACFGLMPQRSTVWY |
| 23 | ACDFGHILNPRSTVWY |
| 26 | CFILMSTVWY |
| 32 | FHILMPQRSTV |
| 34 | ACDEFHIKLNPQRSTVY |
| 37 | AEGIKLPQRSTV |
| 38 | AEGHKLMPQRSTVW |
| 40 | ACDEFGHILNPRSTVWY |
| 41 | ACDEFGHKLNPQRSTVY |
| 45 | ACDFGHILRSTVWY |
| 54 | ACDEFGHIKLNPQRSTVY |
| 55 | AEFGIKLMNRSTVW |
| 58 | ACDFGHILMNPRSTVWY |
| 61 | ACDEFGHIKLNPQRSVWY |
| 62 | AFGIKLMNPRSTVW |
| 64 | ADFGIKLMNPRSTVY |
| 66 | ACFILMNPRSTVY |

| Position | Allowed amino acids |
|---|---|
| 84 | ACDFGHILMNPRSTVWY |
| 86 | ACDEFGHILNPRSTVWY |
| 88 | ACDEFGHILNPRSTVWY |
| 90 | ACDEFGHIKLNPQRSTVY |
| 92 | ADEFGIKLMPQRSTVW |
| 93 | ACDFGHIKLMNPRSTVWY |
| 95 | ACDEFGHILNPRSTVWY |
| 97 | AEFGHIKLMPQRSTVWY |
| 99 | ACDEFGHILNPRSTVY |
| 101 | ACDEFGHIKLNPQRSTVWY |
| 103 | ACDFHILNPRSTVWY |
| 105 | AFGHIKLMPQRSTVW |
| 112 | ACFHIKLMNPRSTV |
| 115 | ACDEFHIKLNQRSTVWY |
| 117 | ACDFGHILMNPQRSTVY |
| 119 | ACDFGHILMNPRSTVWY |
| 124 | ACDEFGHILMNPRSTVWY |
| 127 | AGHIKLPQRSTV |
| 128 | AEFGHIKLMPQRSTVW |

**Table S7. Fentanyl specificity benchmark structures and positions.**
PDB codes are shown for the source of the protein and ligand structure. Position and starting side chain identity are shown for designable positions, and position is shown for packable positions. Position numbering corresponds to PDB numbering as in (Bick et al., 2017).

| Protein | PDB | Designable positions | Packable positions (coupled methods) |
|---|---|---|---|
| Designed fentanyl binder Fen49 | 2QZ3 (protein), 5TZO (ligand) | Q7, W9, N35, V37, N63, Y65, T67, Y69, W71, E78, Y80, P90, R112, P116, W129, Y166, A170, A172 (n = 18) | Y5, D11, T43, R73, L76, V82, W85, Y88, Y108, T110, A115, S117, I118, D121, F125, Q127, V131, A165, V168, Y174, Q175, FEN (n = 22) |

**Table S8. Allowed design for fentanyl dataset.**
Amino acids were allowed in design only if they had high enough sequencing counts to be included in the enrichment and depletion calculations in (Bick et al., 2017). Shown are the amino acid side chains (one letter codes) to which positions were allowed to design. Because Fen49 wild-type identities are disallowed during design (see Methods), positions marked with (†) were mutated to alanine with the FixBB application during preparation of the input structure for design.

| Position | Allowed side chains |
|---|---|
| 9 | ACDEFGHIKLMNPRSTVWY |
| 35 | ACDEFGIKLMNPQRTVWY |
| 37† | ACDEFGHILMNQRSTWY |
| 65† | ACEGLMRSTV |
| 67 | ACDEFGIKLMNPQRSTVWY |
| 69† | ACDFGHIKLNRSTVW |
| 71† | ACDEFGHIKLMNPRSTV |

| Position | Allowed side chains |
|---|---|
| 78 | ACDEFGHKLMPQRSTVWY |
| 90 | ACDEFGHILMNPQRSTVWY |
| 112 | ACDFGHILMPRSTVWY |
| 116 | AEGKLMPQRSTVWY |
| 129 | ACEFGIKLMPQRSTVW |
| 170 | ACEGLPQRSTV |

**Table S9. hGH/hGHr specificity benchmark structures and positions.**
For each library, position and starting side chain identity are shown for designable positions, and position is shown for packable positions. Position numbering corresponds to PDB numbering.

| Library | Designable positions | Packable positions |
|---|---|---|
| A | M14, Y28, N47, P61, D171, I179 (n = 6). | 17, 21, 32, 41, 48, 49, 50, 60, 66, 67, 68, 70, 75, 78, 160, 163, 164, 167, 174, 175, 176, 177, 178, 181, 183, 202, 254, 276, 315, 365, (n = 30) |
| B | H18, Y42, S62, E65, Y164, T175 (n = 6). | 22, 28, 38, 41, 44, 45, 46, 51, 53, 63, 66, 69, 160, 165, 167, 168, 171, 174, 176, 179, 202, 248, 252, 254, 255, 270, 271, 272, 277, 315, 363, 364, (n = 32) |
| C | H21, N29, L45, T60, T67, R178 (n = 6). | 14, 24, 25, 33, 41, 42, 44, 51, 58, 61, 66, 68, 75, 78, 82, 164, 167, 170, 171, 172, 174, 176, 179, 181, 182, 189, 226, 256, 272, 315, 317, 364, 365, (n = 33) |
| D | Q22, S43, E66, R167, F176, R183 (n = 6). | 18, 19, 21, 23, 24, 25, 26, 28, 40, 60, 61, 62, 63, 67, 72, 75, 78, 79, 82, 164, 172, 175, 179, 184, 254, 276, 277, 364, (n = 28) |
| E | D26, F44, P48, R64, K168, E174 (n = 6). | 14, 17, 18, 21, 22, 25, 45, 47, 49, 50, 51, 52, 53, 56, 68, 157, 160, 164, 169, 172, 203, 221, 225, 226, 254, 256, 310, 313, 315, 363, 364, (n = 31) |
| F | F25, K41, Q46, N63, K172 (n = 5). | 21, 26, 28, 29, 32, 36, 38, 42, 45, 56, 60, 62, 65, 66, 82, 160, 164, 167, 168, 169, 176, 226, 252, 254, 258, 270, 272, 277, 364, (n = 29) |

**Table S10. Herceptin/HER2 specificity benchmark structures and positions.**
Design and packable positions are shown for each library. Design positions are listed in Kabat numbering (Martin, 1996). For packable positions, numbering corresponds to consecutive renumbering of the 312 positions in combined chain A positions 1-106, chain B positions 1-119, and chain C positions 511-607. Herceptin Library D is omitted because the experimental data were dominated by the wild-type sequence.

| Library | Designable positions | Packable positions |
| --- | --- | --- |
| A | VL94, VH33, VH50, VH56, VH58, VH95 (n = 6). | 93, 95, 138, 140, 141, 153, 155, 157, 158, 161, 164, 166, 176, 204, 211, 213, 272, 273, 275, 276, 287, 288 (n = 22). |
| B | VL30, VL91, VL92, VH50, VH95, VH99, VH100a (n = 7). | 28, 29, 31, 32, 66, 71, 90, 93, 94, 138, 139, 141, 153, 155, 157, 164, 165, 176, 204, 208, 210, 212, 213, 273, 275, 284, 285, 286, 287, 288, 296, 298, 301, 303, 305, 307 (n = 36). |
| C | VL49, VL53, VL91, VH98, VH99, VH100, VH100a (n = 7). | 32, 46, 48, 50, 52, 54, 90, 92, 205, 212, 214, 285, 286, 287, 288, 296, 298, 308 (n = 18). |
| E | VL49, VL53, VL55, VH100, VH102 (n = 5). | 46, 48, 50, 52, 54, 56, 58, 108, 110, 203, 204, 205, 211, 212, 214, 216, 308 (n = 17). |

**Table S11. Number of designed sequence positions in PPS gain/loss/neutral zones.**
Values for CoupledMoves represent averages ± standard deviation for CM-BR, CM-FKIC, and CM-WKIC. The charged or polar category includes arginine, histidine, lysine, aspartate, glutamate, serine, threonine, asparagine, glutamine, tyrosine, and cysteine. The hydrophobic category includes alanine, phenylalanine, glycine, isoleucine, leucine, methionine, valine, tryptophan, and proline.

| | | **Amino acid category** | | |
|---|---|---|---|---|
| | | **all** | **charged or polar** | **hydro-phobic** |
| **Coupled Moves** | gain | 43±3 | 16±1 | 28±2 |
| | loss | 32±4 | 25±3 | 6±1 |
| | neutral | 81±6 | 26±3 | 55±3 |
| **Backrub Ensemble** | gain | 37 | 13 | 24 |
| | loss | 41 | 28 | 13 |
| | neutral | 78 | 26 | 52 |
| **Fast Design** | gain | 28 | 10 | 18 |
| | loss | 46 | 33 | 13 |
| | neutral | 82 | 24 | 58 |
| **Fixed Backbone** | gain | 13 | 4 | 9 |
| | loss | 50 | 34 | 16 |
| | neutral | 93 | 29 | 64 |

**Table S12. Compute time.**
Total compute time for each method. Values represent the mean and standard deviation across 400 trajectories.

| Method | Time (hours) | | |
|---|---|---|---|
| CM-BR | 52 | ± | 27 |
| CM-FKIC | 75 | ± | 43 |
| CM-WKIC | 73 | ± | 27 |
| FastDesign | 1,502 | ± | 1,377 |
| Backrub<br>+ FixBB | 117<br>5 | ±<br>± | 115<br>3 |
| = Backrub Ensemble | 122 | ± | 115 |

### SUPPLEMENTAL FIGURE LEGENDS

**Figure S1. Position profile similarity and RankTop for all designed positions (n) for each protein family in the cofactor dataset.**
Distributions are shown as boxplots, while values for individual positions are overlaid as swarms of black points. For PPS (left), a value of 1 means the design method perfectly recapitulated the known sequence profile, whereas a value of zero means that the design method did not model any of the amino acid side chain identities from the known profile. For RankTop (right), a value of 1 means that the design method correctly identified the most frequent amino acid side chain observed in the known profile, whereas a RankTop of 20 means that side chain was observed with zero frequency, or that all side chains were modeled with some frequency and the top known side chain was the least frequent. Median is marked with a horizontal black line, and notches represent a 95% confidence interval (CI) around the median; when CI extends past the quartiles, notches extend beyond the box, leading to a "flipped" appearance. The boxplot covers the second and third quartiles, and the vertical whiskers mark 1.5 times the inter-quartile range.

**Figure S2. Profile similarity and rank top for all designed positions (n) for individual libraries of the Herceptin/HER2 dataset.**
Distributions are shown as boxplots, while values for individual positions are overlaid as swarms of black points. For PPS (left), a value of 1 means the design method perfectly recapitulated the known sequence profile, whereas a value of zero means that the design method did not model any of the amino acid side chain identities from the known profile. For RankTop (right), a value of 1 means that the design method correctly identified the most frequent amino acid side chain observed in the known profile, whereas a RankTop of 20 means that side chain was observed with zero frequency, or that all side chains were modeled with some frequency and the top known side chain was the least frequent. Median is marked with a horizontal black line, and notches represent a 95% confidence interval (CI) around the median; when CI extends past the quartiles, notches extend beyond the box, leading to a "flipped" appearance. The boxplot covers the second and third quartiles, and the vertical whiskers mark 1.5 times the inter-quartile range.

**Figure S3. Profile similarity and rank top for all designed positions (n) for individual libraries of the hGH/hGHR dataset.**
Distributions are shown as boxplots, while values for individual positions are overlaid as swarms of black points. For PPS (left), a value of 1 means the design method perfectly recapitulated the known sequence profile, whereas a value of zero means that the design method did not model any of the amino acid side chain identities from the known profile. For RankTop (right), a value of 1 means that the design method correctly identified the most frequent amino acid side chain observed in the known profile, whereas a RankTop of 20 means that side chain was observed with zero frequency, or that all side chains were modeled with some frequency and the top known side chain was the least frequent. Median is marked with a horizontal black line, and notches represent a 95% confidence interval (CI) around the median; when CI extends past the quartiles, notches extend beyond the box, leading to a "flipped" appearance. The boxplot covers the second and third quartiles, and the vertical whiskers mark 1.5 times the inter-quartile range.

**Figure S4. Sequence logos for predicted and known binding site sequences of the cofactor dataset.**

The height of each letter is proportional to its contribution to the column's information content. The height of each column is inversely proportional to the sequence variation at that position.

**Figure S5. Sequence logos for predicted and known binding site sequences of the DIG10 dataset.**
The height of each letter is proportional to its contribution to the information content of the column. The height of each column is inversely proportional to the sequence variation at that position. The experimental profile shows amino acid residues that were enriched in the experimental selection.

**Figure S6. Sequence logos for predicted and known binding site sequences of the Fen49 dataset.**
The height of each letter is proportional to its contribution to the information content of the column. The height of each column is inversely proportional to the sequence variation at that position. Experimental data are taken from sort 4 of the library (Bick et al., 2017).

**Figure S7. Sequence logos for predicted and known binding site sequences for the Herceptin/HER2 dataset.**
The height of each letter is proportional to its contribution to the information content of the column. The height of each column is inversely proportional to the sequence variation at that position. Different experimental libraries (Lib A, B, C, E) are indicated. Library D was omitted because the experimental data were dominated by the wild-type sequence. Residues are labeled with Kabat numbering.

**Figure S8. Sequence logos for predicted and known binding site sequences of the hGH/hGHR dataset.**
The height of each letter is proportional to its contribution to the information content of the column. The height of each column is inversely proportional to the sequence variation at that position. Different experimental libraries (Lib A, B, C, D, E, F) are indicated.

**Figure S9. Profile similarity and RankTop as a function of known sequence entropy for the hGH/hGHR, DIG10 and Fen49 datasets.**
Each point represents one sequence position. The Herceptin/HER2 and Cofactor datasets are shown in **Figure 5** in the main text. For each dataset (indicated in the header), profile similarity and RankTop are binned by entropy of the known sequence profile at each position (low: entropy $\leq 0.33$, medium: $0.33 < \text{entropy} \leq 0.67$, and high: entropy $> 0.67$). The number of low entropy positions in these three datasets is small. The median is marked with a horizontal black line.

**Figure S10. Design entropy.**
Shown are the distributions of sequence entropy of the design sequence profiles for each designed position in each benchmark. The median of the distributions is marked with a white dot. Second and third quartiles are marked by the thick black bar, and the thin bar marks 1.5 times the inter-quartile range. The width of the violins is determined by the number of observations in each bin, and bins are defined using Scott's normal reference rule.

**Figure S11. Position profile similarity and RankTop as a function of similarity between the input sequence and the known profile at each position**.

When a preferred side chain from the known sequence profiles is not present in the input sequence, methods can achieve "gain" (green) by identifying correct amino acids with high frequency or rank. Alternatively, when a preferred side chains is present in the input, inaccurate design can cause "loss" (red). Only positions with low and medium entropy ($\leq 0.67$) are considered. (**A**) Left: PPS as a function of similarity to the input sequence for all profile datasets. Each point represents one position in the protein sequence, colored by design method. Right: Quantifications of number of designed sequence positions in gain, loss, and neutral zones. Gain and loss zones are defined by a threshold of 0.1 difference between input-known PPS and design-known PPS. (**B**) Left: Boxplots of each method's RankTop as a function of similarity to the input sequence. The median of the distributions is marked with a horizontal line. Second and third quartiles are marked by the box, and the whiskers extend to 1.5 times the inter-quartile range. The top amino acid from the known profile is assigned a rank of 1 if it is present in the input sequence, or a rank of 20 if it is not. All profile datasets are shown except Fen49, which is omitted because the fentanyl deep sequencing data do not include the input sequence. For the digoxigenin dataset, there are no consensus positions for which the top experimentally selected side chain was present in the starting sequence. Right: Quantification of sequence positions in gain, loss, and neutral zones for RankTop values.

**Figure S12. Performance as a function of number of design trajectories.**
Comparison of median PPS, design entropy and RankTop as a function of number of design trajectories (*n*) for the Cofactor, Herceptin, and hGH/hGHR datasets for each method.

### SUPPLEMENTAL FIGURES

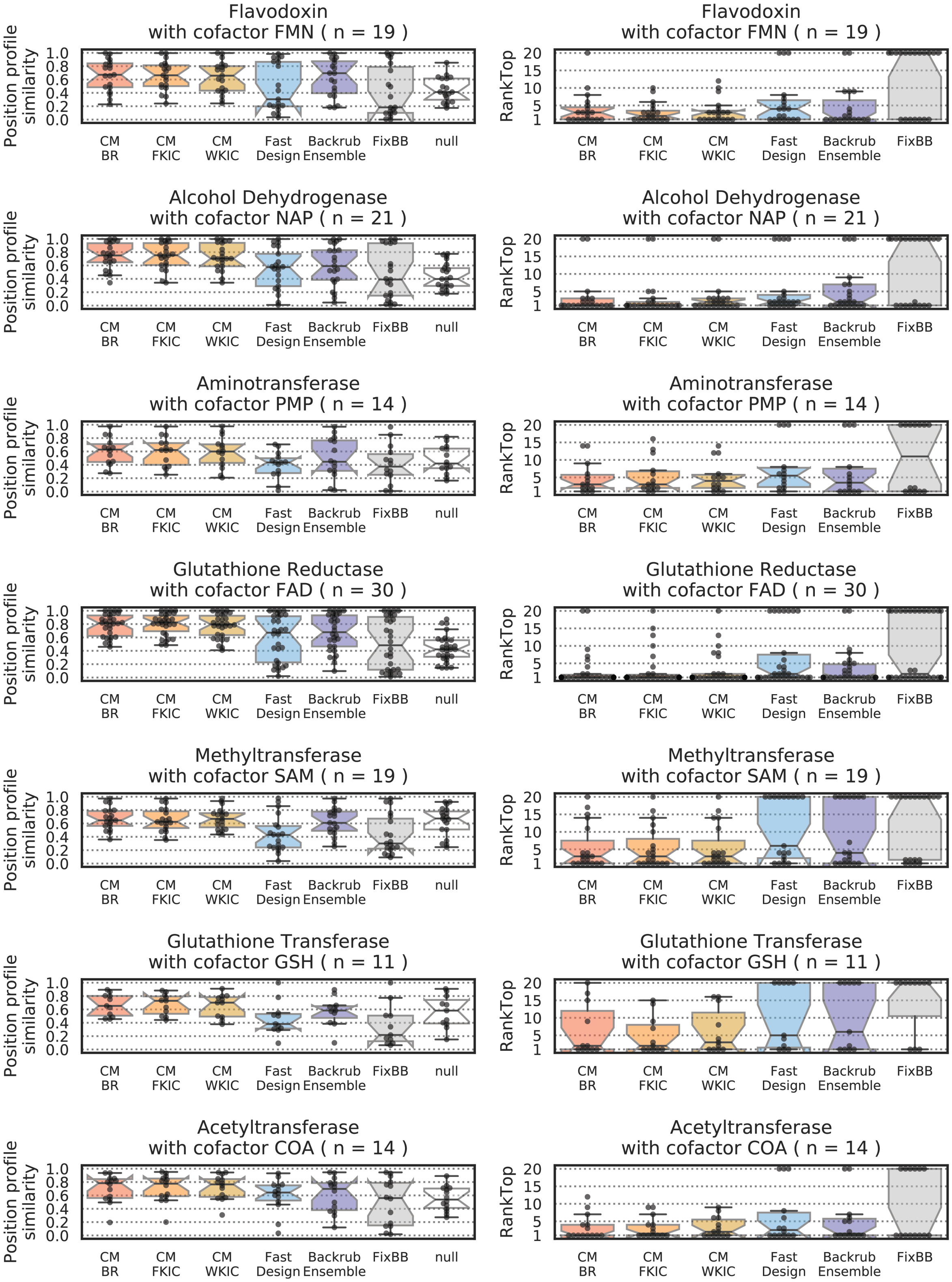


**Figure S1. Position profile similarity and RankTop for all designed positions (n) for each protein family in the cofactor dataset.**

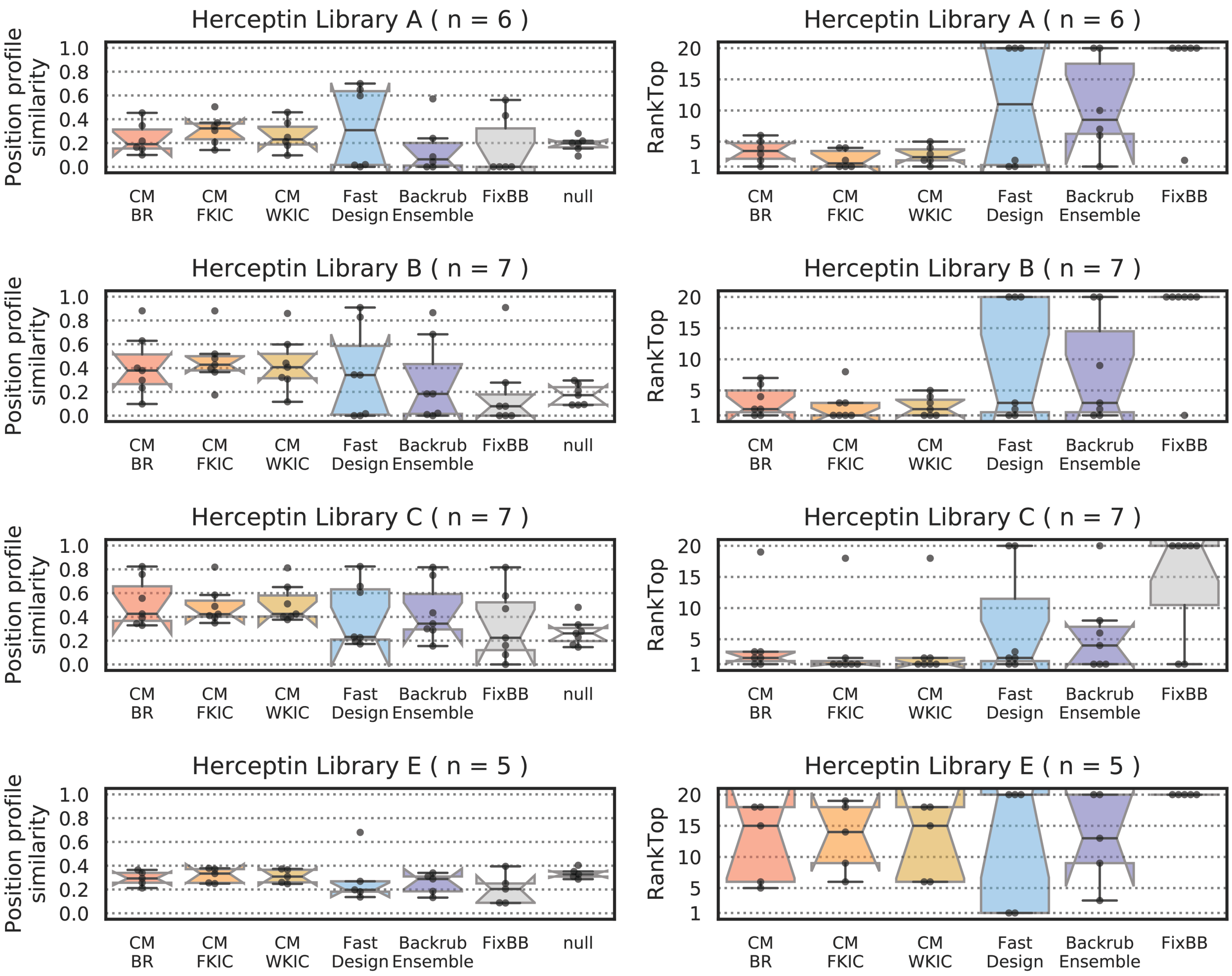


**Figure S2. Profile similarity and rank top for all designed positions (n) for individual libraries of the Herceptin/HER2 dataset.**

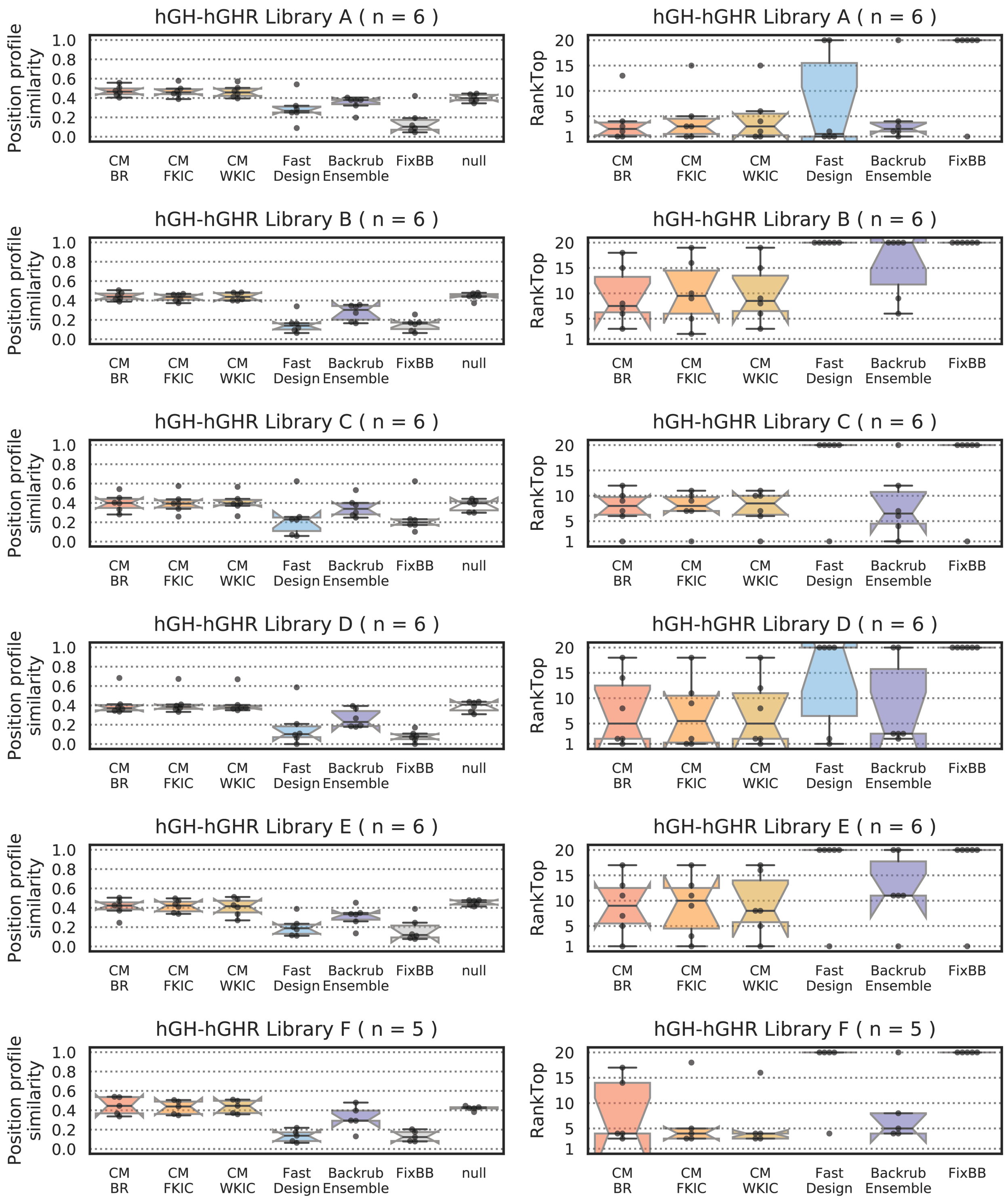


**Figure S3. Profile similarity and rank top for all designed positions (n) for individual libraries of the hGH/hGHR dataset.**

### Cofactor positions

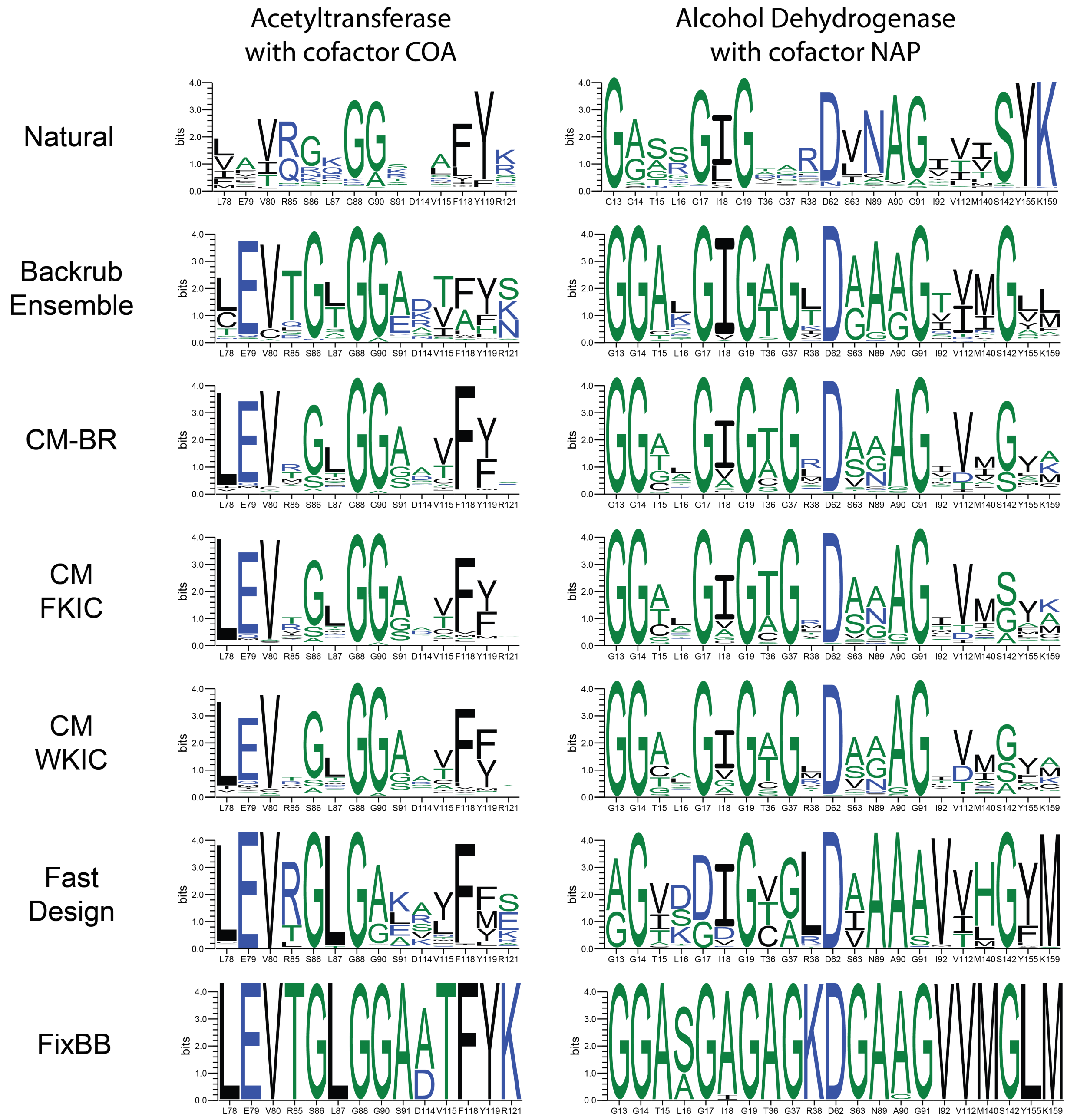


**Figure S4. Sequence logos for predicted and known binding site sequences of the cofactor dataset.**

**Continued on next page.**

### Cofactor positions

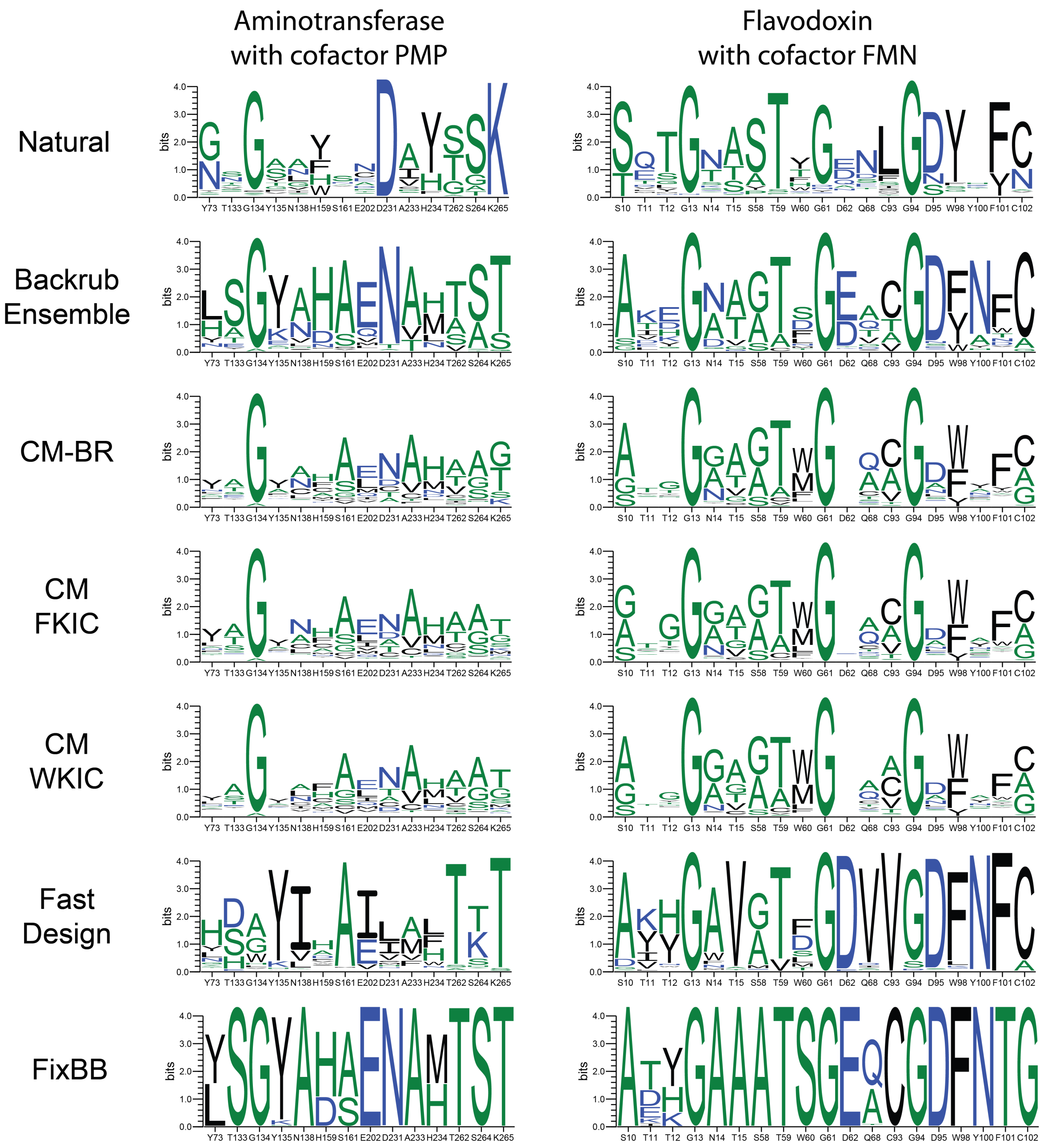


**Figure S4, continued. Sequence logos for predicted and known binding site sequences of the cofactor dataset.**

**Continued on next page.**

### Cofactor positions

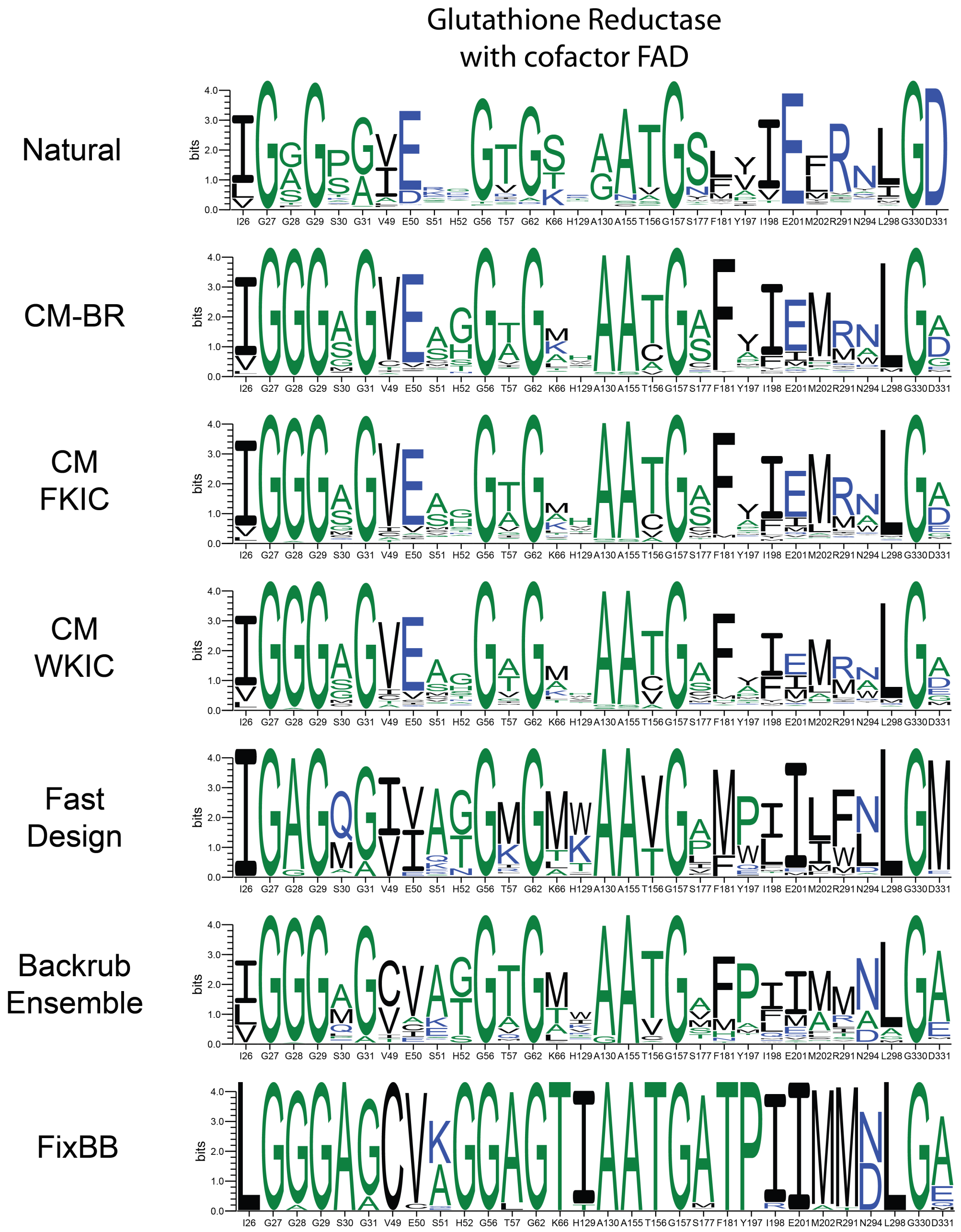


**Figure S4, continued. Sequence logos for predicted and known binding site sequences of the cofactor dataset.**
**Continued on next page.**

### Cofactor positions

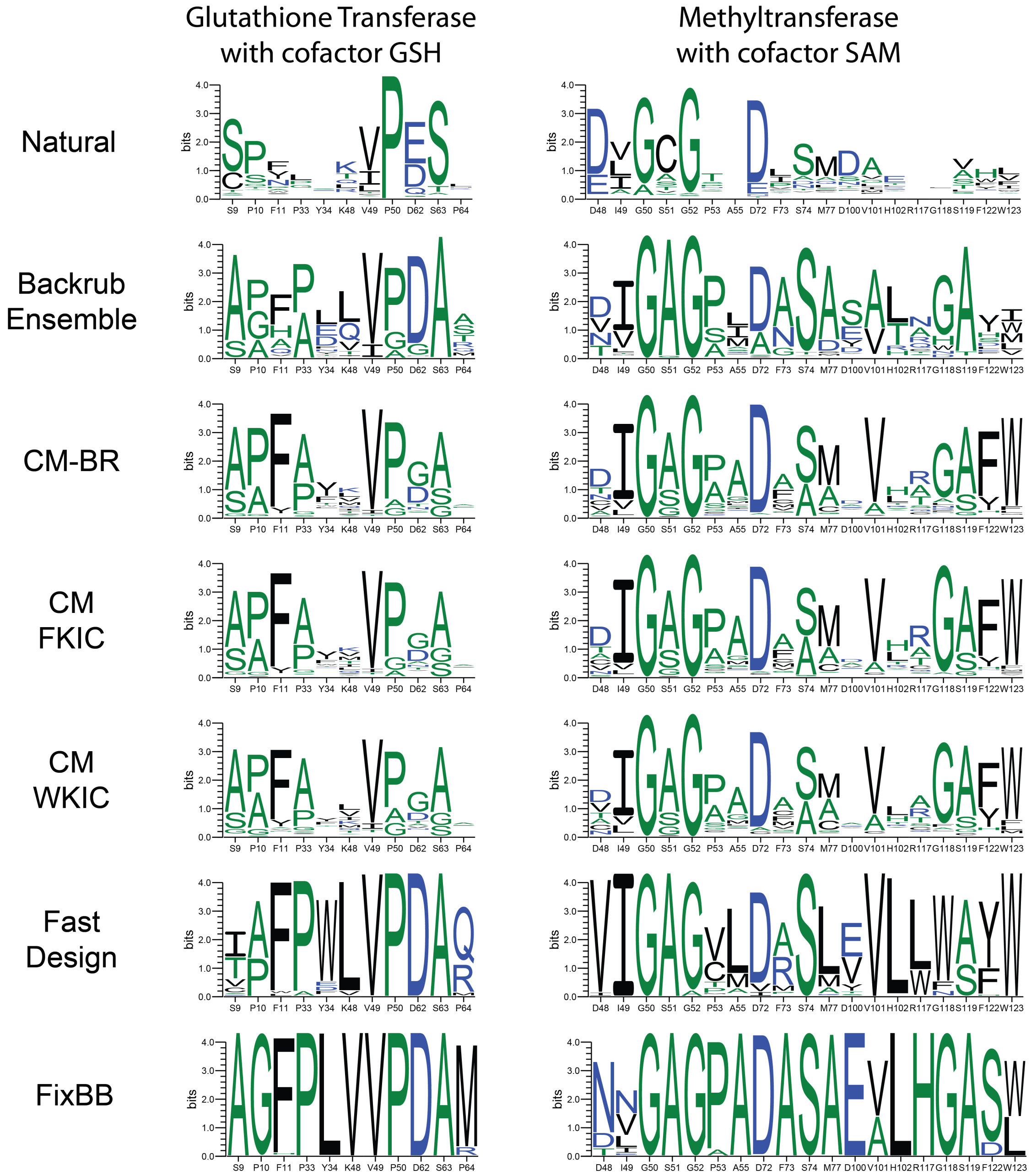


**Figure S4, continued. Sequence logos for predicted and known binding site sequences of the cofactor dataset.**

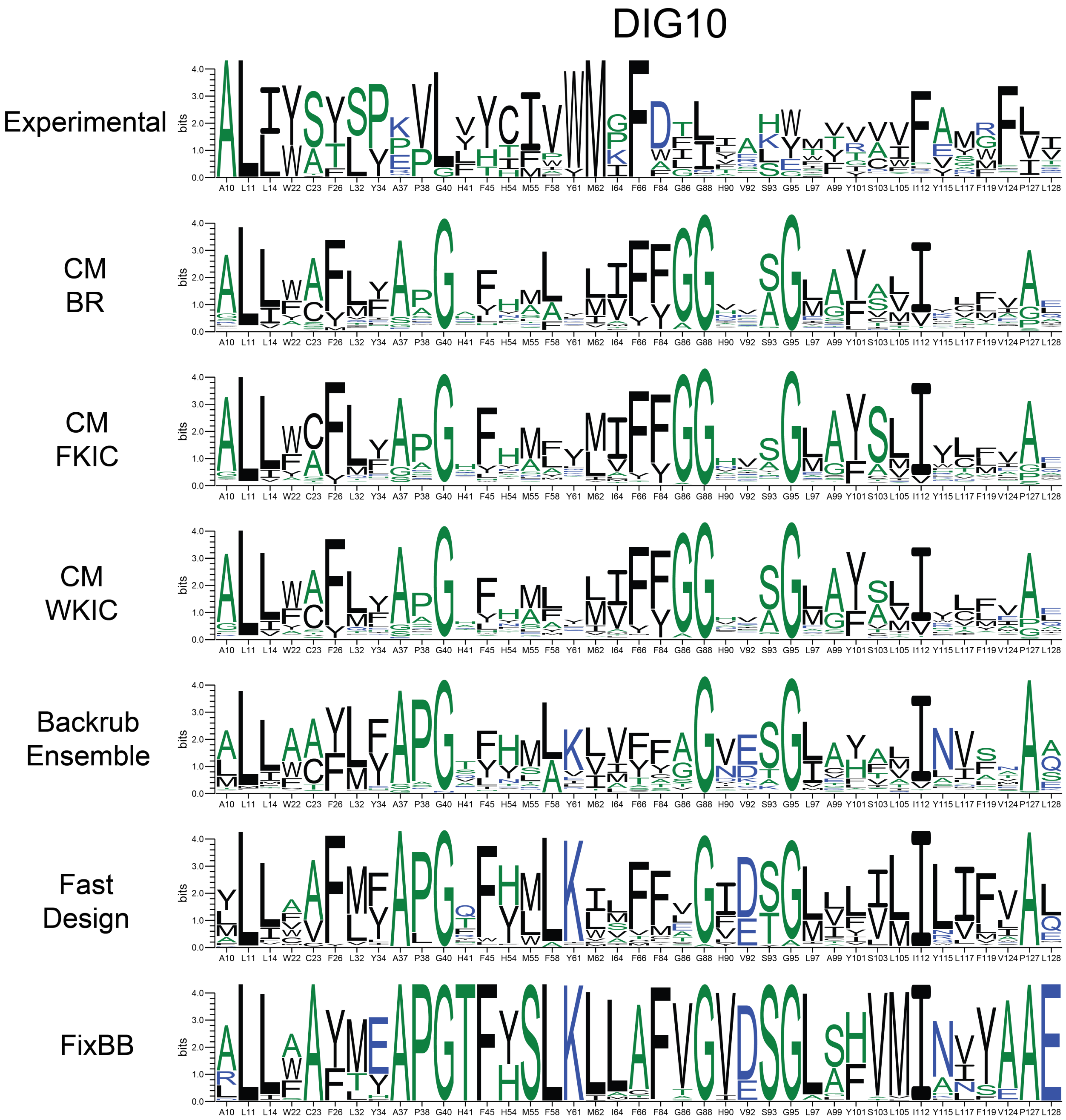


**Figure S5. Sequence logos for predicted and known binding site sequences of the DIG10 dataset.**

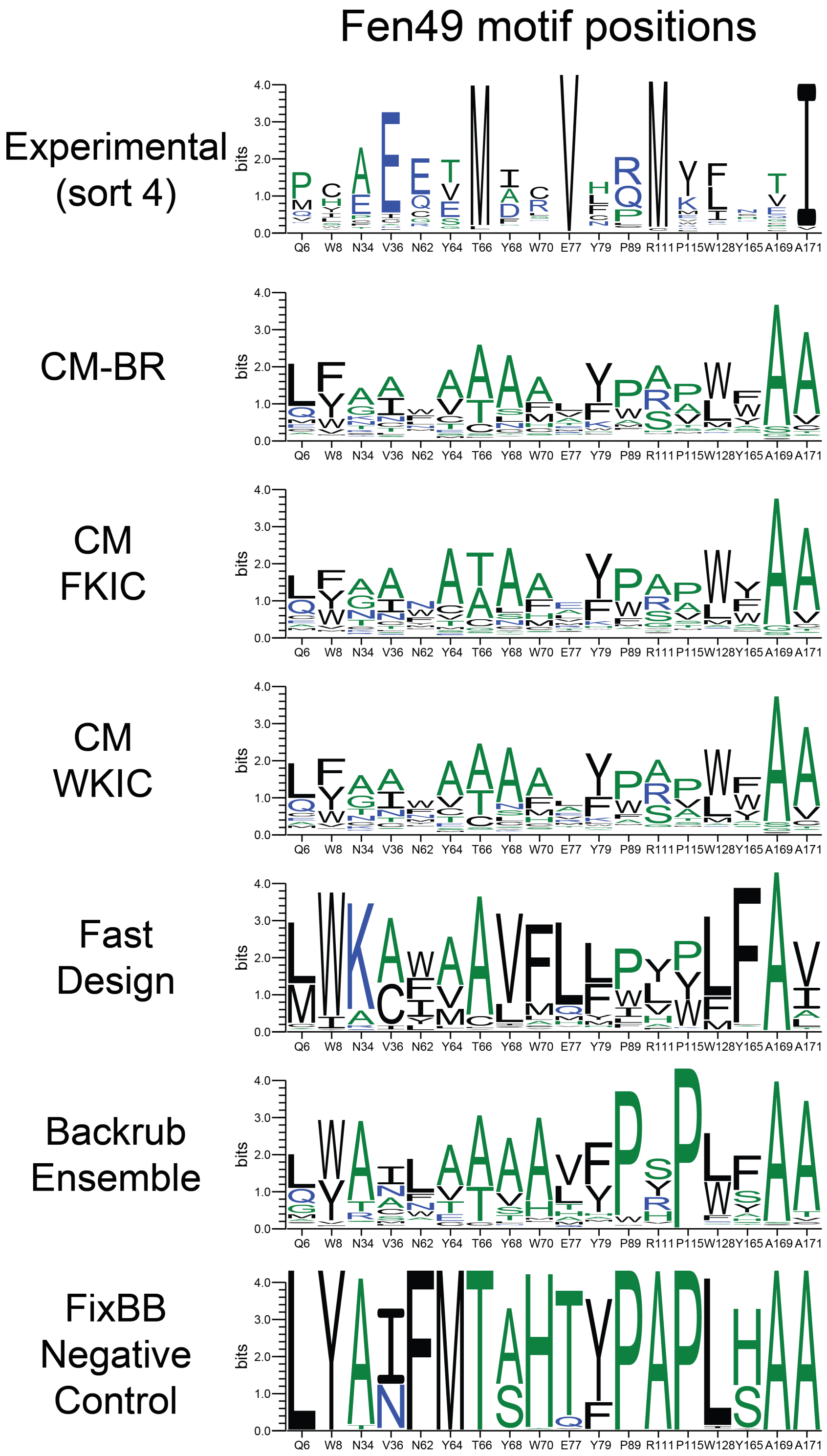


**Figure S6. Sequence logos for predicted and known binding site sequences of the Fen49 dataset.**

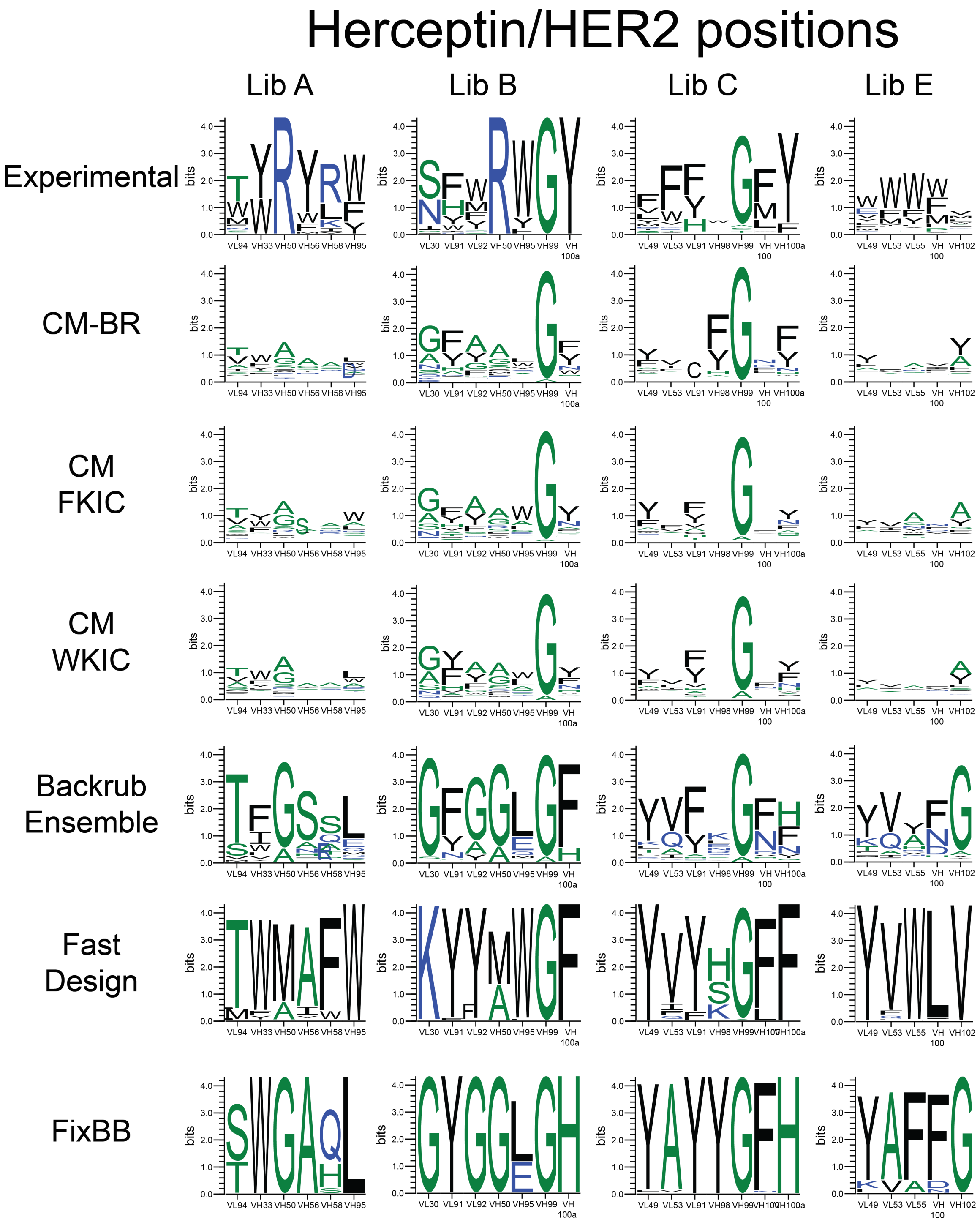


**Figure S7. Sequence logos for predicted and known binding site sequences for the Herceptin/HER2 dataset.**

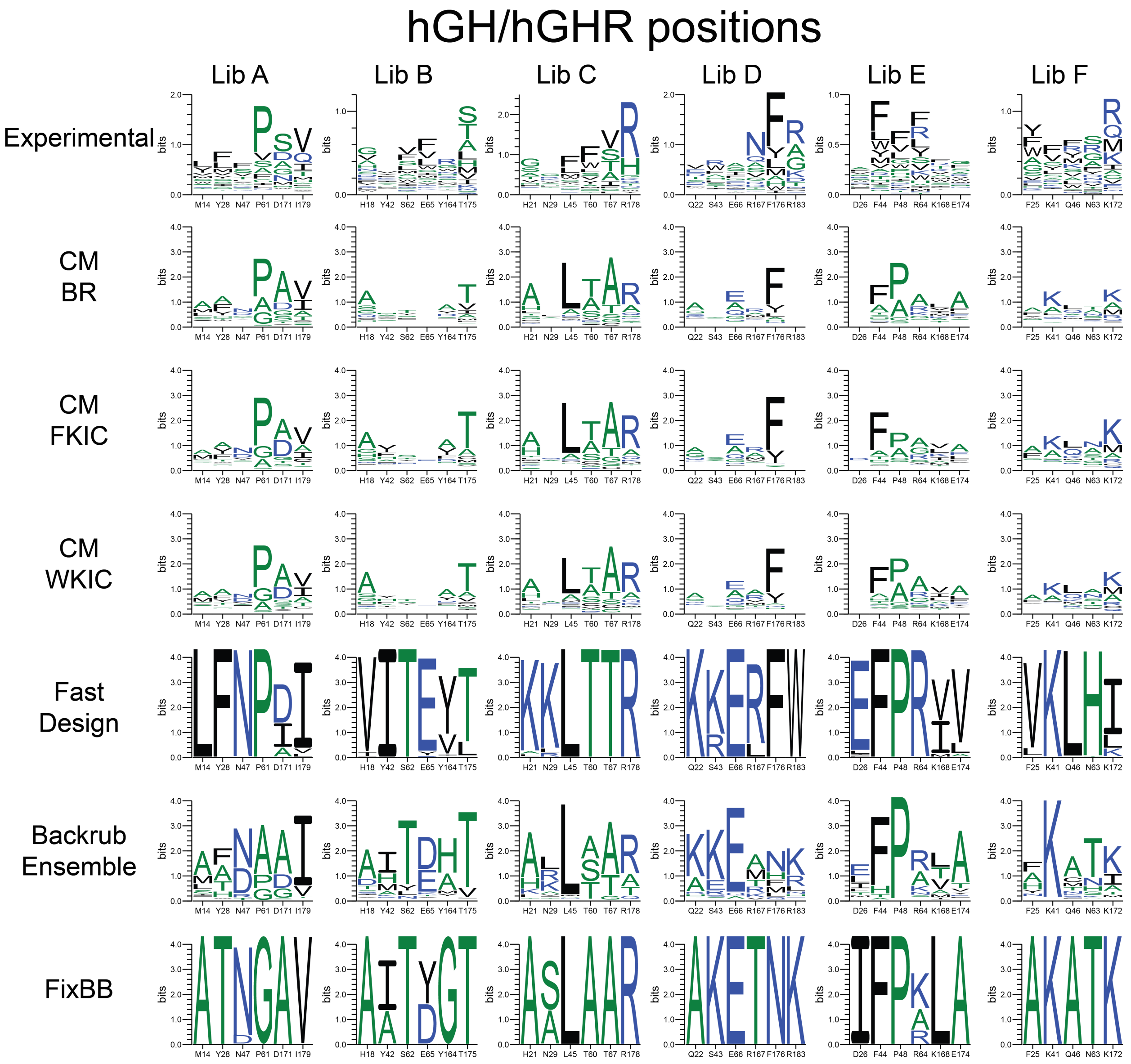


**Figure S8. Sequence logos for predicted and known binding site sequences of the hGH/hGHR dataset.**

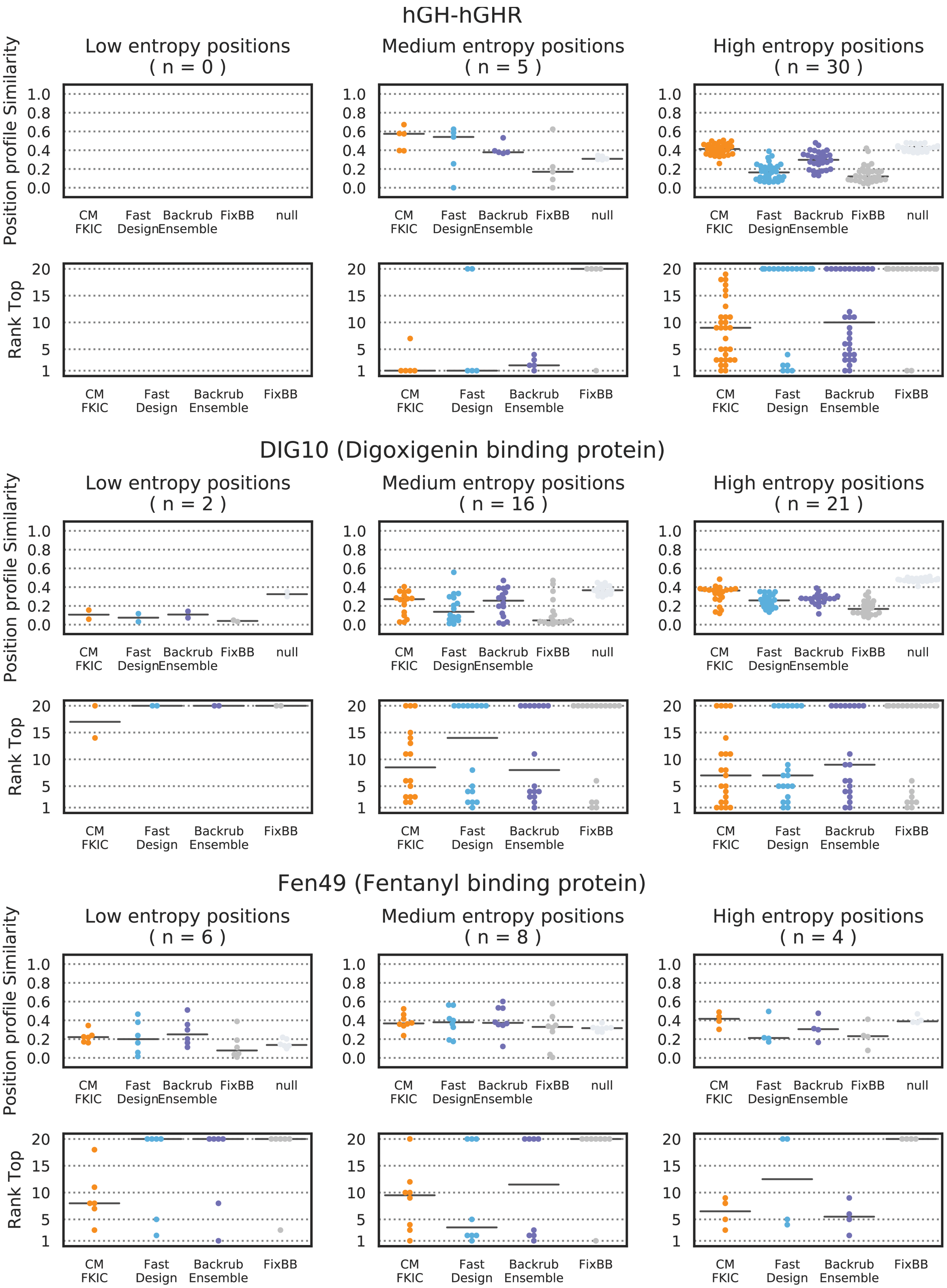


**Figure S9. Profile similarity and RankTop as a function of known sequence entropy for the hGH/hGHR, DIG10 and Fen49 datasets.**

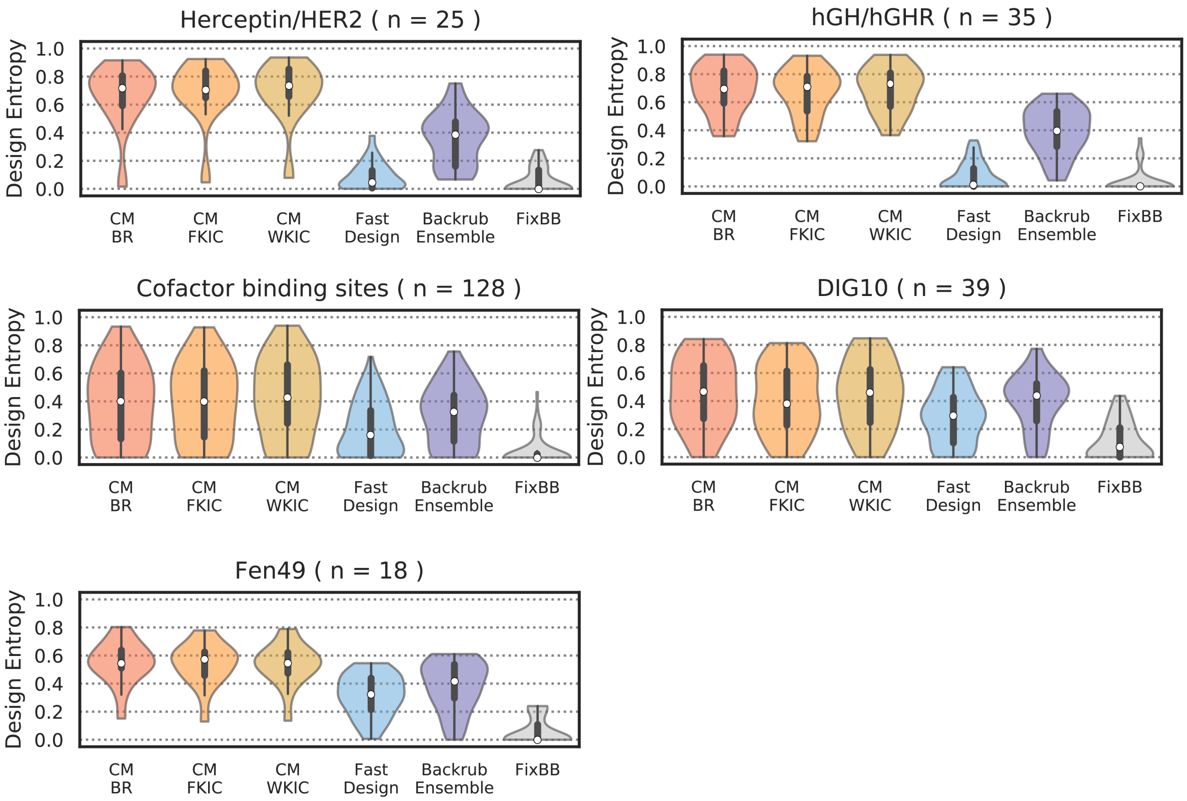


**Figure S10. Design entropy.**

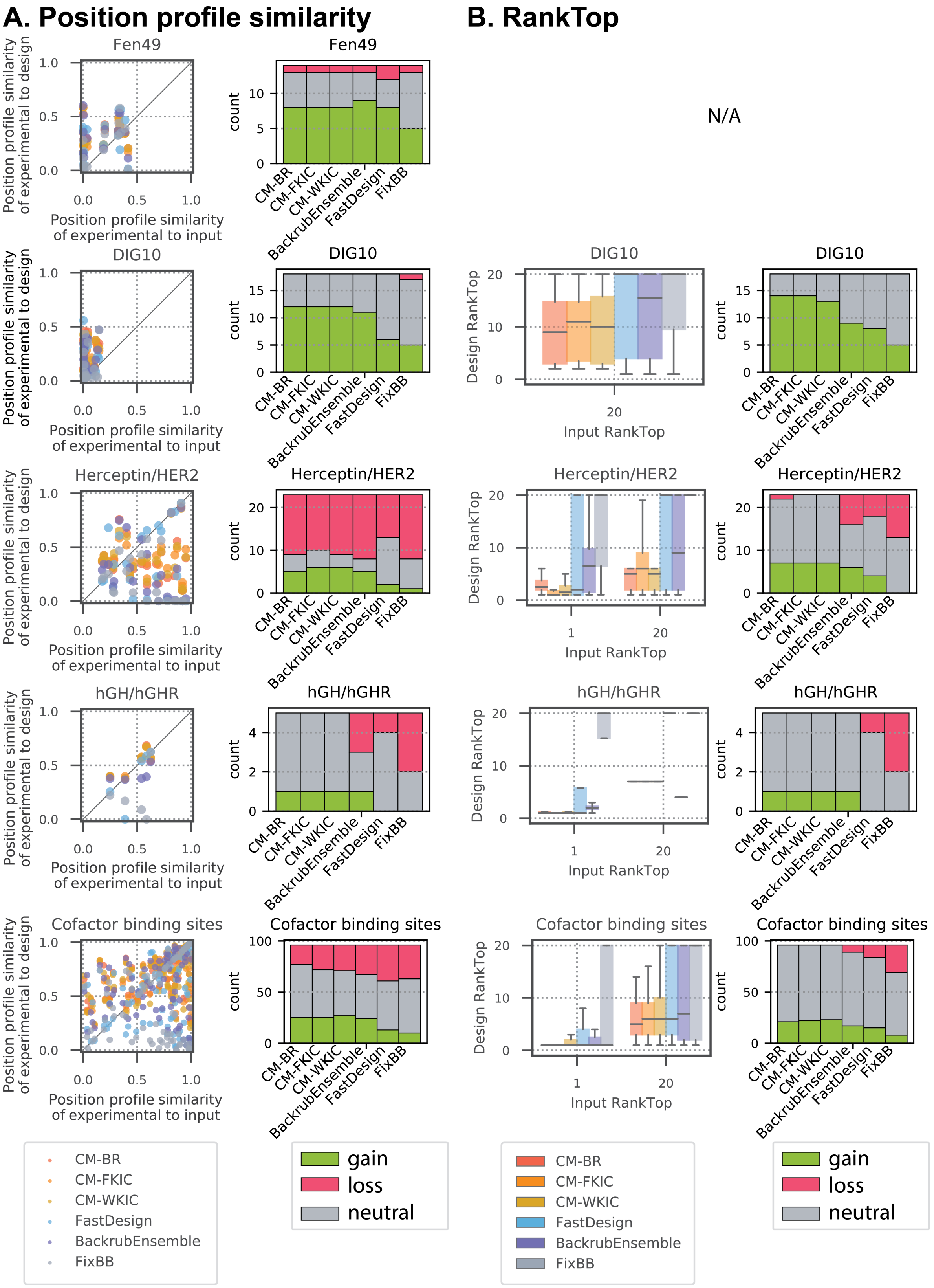


**Figure S11. Position profile similarity and RankTop as a function of similarity between the input sequence and the known profile at each position.**

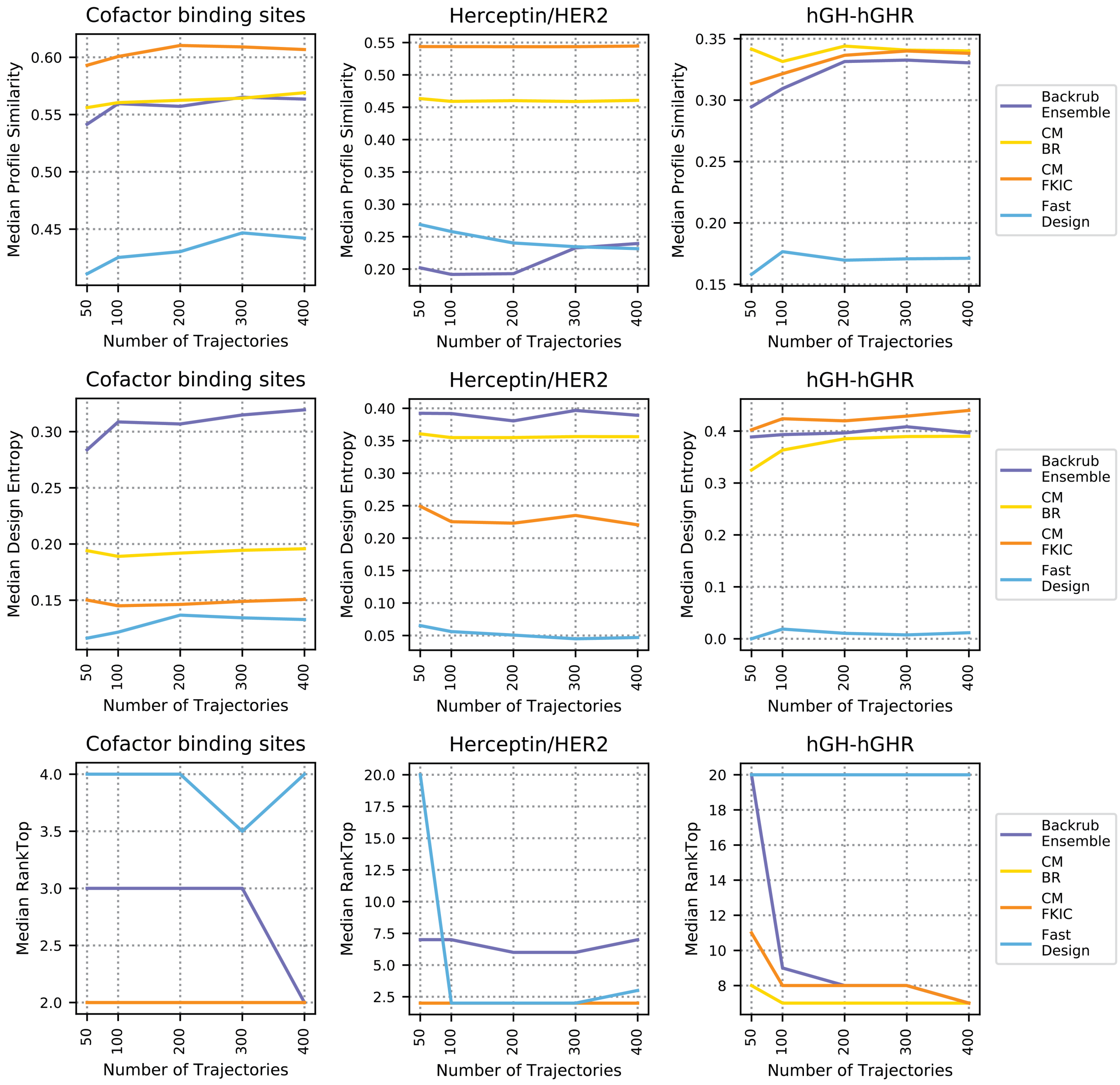


**Figure S12. Performance as a function of number of design trajectories.**

### Rosetta command lines and XML scripts

##### CM-BR (with ligand)

```
Rosetta/main/source/bin/coupled_moves.default.linuxgccrelease -s pdb -
mute protocols.backrub.BackrubMover -ex1 -ex2 -extrachi_cutoff 0 -
nstruct 1 -ignore_unrecognized_res -score::weights ref2015 -
extra_res_fa ligand_name.params -resfile resfile -coupled_moves::mc_kt
2.4 -coupled_moves::boltzmann_kt 2.4 -coupled_moves::ntrials 1000 -
coupled_moves::initial_repack false -coupled_moves::ligand_mode true -
coupled_moves::ligand_weight 2 -coupled_moves::fix_backbone false -
coupled_moves::bias_sampling true -coupled_moves::bump_check true -
coupled_moves::backbone_mover backrub -
coupled_moves::exclude_nonclashing_positions true -nstruct 400
```

##### CM-BR (without ligand)

```
Rosetta/main/source/bin/coupled_moves.default.linuxgccrelease -s pdb -
mute protocols.backrub.BackrubMover -ex1 -ex2 -extrachi_cutoff 0 -
nstruct 1 -ignore_unrecognized_res -score::weights ref2015 -resfile
resfile -coupled_moves::mc_kt 2.4 -coupled_moves::boltzmann_kt 2.4 -
coupled_moves::ntrials 1000 -coupled_moves::initial_repack false -
coupled_moves::fix_backbone false -coupled_moves::bias_sampling true -
coupled_moves::bump_check true -coupled_moves::backbone_mover backrub
-coupled_moves::exclude_nonclashing_positions true -nstruct 400
```

##### CM-FKIC (with ligand)

```
Rosetta/main/source/bin/coupled_moves.default.linuxgccrelease -s
name.pdb -mute protocols.backrub.BackrubMover -ex1 -ex2 -
extrachi_cutoff 0 -nstruct 1 -ignore_unrecognized_res -score::weights
ref2015 -extra_res_fa name.params -resfile name.resfile -
coupled_moves::mc_kt 2.4 -coupled_moves::boltzmann_kt 2.4 -
coupled_moves::ntrials 1000 -coupled_moves::initial_repack false -
coupled_moves::ligand_mode true -coupled_moves::ligand_weight 2 -
coupled_moves::fix_backbone false -coupled_moves::bias_sampling true -
coupled_moves::bump_check true -
coupled_moves::exclude_nonclashing_positions true -
coupled_moves::backbone_mover kic -coupled_moves::kic_perturber
fragment -loops:frag_sizes 9 3 -loops:frag_files name.200.9mers.gz
name.200.3mers.gz -nstruct 400
```

**CM-FKIC (without ligand)**

```
Rosetta/main/source/bin/coupled_moves.default.linuxgccrelease -s
name.pdb -mute protocols.backrub.BackrubMover -ex1 -ex2 -
extrachi_cutoff 0 -nstruct 1 -ignore_unrecognized_res -score::weights
ref2015 -resfile name.resfile -coupled_moves::mc_kt 2.4 -
coupled_moves::boltzmann_kt 2.4 -coupled_moves::ntrials 1000 -
coupled_moves::initial_repack false -coupled_moves::ligand_mode false
-coupled_moves::fix_backbone false -coupled_moves::bias_sampling true
-coupled_moves::bump_check true -
coupled_moves::exclude_nonclashing_positions true -
coupled_moves::backbone_mover kic -coupled_moves::kic_perturber
fragment -loops:frag_sizes 9 3 -loops:frag_files name.200.9mers.gz
name.200.3mers.gz -nstruct 400
```

**CM-WKIC (with ligand)**

```
Rosetta/main/source/bin/coupled_moves.default.linuxgccrelease -s
name.pdb -mute protocols.backrub.BackrubMover -ex1 -ex2 -
extrachi_cutoff 0 -nstruct 1 -ignore_unrecognized_res -score::weights
ref2015 -extra_res_fa name.params -resfile name.resfile -
coupled_moves::mc_kt 2.4 -coupled_moves::boltzmann_kt 2.4 -
coupled_moves::ntrials 1000 -coupled_moves::initial_repack false -
coupled_moves::ligand_mode true -coupled_moves::ligand_weight 2 -
coupled_moves::fix_backbone false -coupled_moves::bias_sampling true -
coupled_moves::bump_check true -
coupled_moves::exclude_nonclashing_positions true -
coupled_moves::backbone_mover kic -coupled_moves::kic_perturber
walking -nstruct 400
```

**CM-WKIC (without ligand)**

```
Rosetta/main/source/bin/coupled_moves.default.linuxgccrelease -s
name.pdb -mute protocols.backrub.BackrubMover -ex1 -ex2 -
extrachi_cutoff 0 -nstruct 1 -ignore_unrecognized_res -score::weights
ref2015 -extra_res_fa name.params -resfile name.resfile -
coupled_moves::mc_kt 2.4 -coupled_moves::boltzmann_kt 2.4 -
coupled_moves::ntrials 1000 -coupled_moves::initial_repack false -
coupled_moves::fix_backbone false -coupled_moves::bias_sampling true -
coupled_moves::bump_check true -
coupled_moves::exclude_nonclashing_positions true -
coupled_moves::backbone_mover kic -coupled_moves::kic_perturber
walking -nstruct 400
```

**FastDesign (with ligand)**

```
Rosetta/main/source/bin/relax.default.linuxgccrelease -s name.pdb -
resfile name.resfile -extra_res_fa ligand_name.params -ex1 -ex2 -
extrachi_cutoff 0 -nstruct 400 -in:file:fullatom -relax:fast -
relax:respect_resfile -relax:constrain_relax_to_start_coords -
relax:coord_cst_stdev .5
```

**FastDesign (without ligand)**

```
Rosetta/main/source/bin/relax.default.linuxgccrelease -s name.pdb -
resfile name.resfile -ex1 -ex2 -extrachi_cutoff 0 -nstruct 400 -
in:file:fullatom -relax:fast -relax:respect_resfile -
relax:constrain_relax_to_start_coords -relax:coord_cst_stdev .5
```

**BackrubEnsemble step 1: Backrub ensemble generation (with ligand)**

```
Rosetta/main/source/bin/backrub.default.linuxgccrelease -
score::weights ref2015 -s name.pdb -nstruct 400 -
ignore_unrecognized_res -extra_res_fa ligand_name.params -
backrub:ntrials 10000 -mc_kt 1.2 -max_atoms 12
```

**BackrubEnsemble step 1: Backrub ensemble generation (without ligand)**

```
Rosetta/main/source/bin/backrub.default.linuxgccrelease -
score::weights ref2015 -s name.pdb -nstruct 400 -
ignore_unrecognized_res -backrub:ntrials 10000 -mc_kt 1.2 -max_atoms
12
```

**BackrubEnsemble step 2: Design on backrub ensemble (with ligand)**

```
Rosetta/main/source/bin/rosetta_scripts.default.linuxgccrelease -
parser:protocol FBBRS.xml -parser:script_vars res_file=name.resfile -s
name_ensemble_member.pdb -nstruct 400 -ignore_unrecognized_res -
extra_res_fa ligand_name.params
```

**BackrubEnsemble step 2: Design on backrub ensemble (without ligand)**

```
Rosetta/main/source/bin/rosetta_scripts.default.linuxgccrelease -
parser:protocol FBBRS.xml -parser:script_vars res_file=name.resfile -s
name_ensemble_member.pdb -nstruct 400 -ignore_unrecognized_res
```

**BackrubEnsemble step 2: Design on backrub ensemble, file name FBBRS.xml**

```
<ROSETTASCRIPTS>
    <SCOREFXNS>
    </SCOREFXNS>
    <RESIDUE_SELECTORS>
    </RESIDUE_SELECTORS>
    <TASKOPERATIONS>
<ReadResfile name="resfile" filename="%%res_file%%" />
      <ExtraRotamers name="ex1" chi="1" />
      <ExtraRotamers name="ex2" chi="2" />
      <ExtraChiCutoff name="exchi0" extrachi_cutoff="0" />
    </TASKOPERATIONS>
    <FILTERS>
    </FILTERS>
    <MOVERS>
      <PackRotamersMover name="pack_rot"
task_operations="resfile,ex1,ex2,exchi0" />
    </MOVERS>
    <APPLY_TO_POSE>
    </APPLY_TO_POSE>
    <PROTOCOLS>
      <Add mover="pack_rot" />
    </PROTOCOLS>
    <OUTPUT/>
</ROSETTASCRIPTS>
```

**FixBB control (with ligand)**

```
Rosetta/main/source/bin/rosetta_scripts.default.linuxgccrelease -
parser:protocol FBBRS.xml -parser:script_vars res_file=name.resfile -s
name.pdb -nstruct 400 -ignore_unrecognized_res -extra_res_fa
ligand_name.params
```

**FixBB control (without ligand)**

```
Rosetta/main/source/bin/rosetta_scripts.default.linuxgccrelease -
parser:protocol FBBRS.xml -parser:script_vars res_file=name.resfile -s
name.pdb -nstruct 400 -ignore_unrecognized_res -extra_res_fa
ligand_name.params
```
